# VelOT: kinetic-free RNA velocity inference via optimal transport, flow-field smoothing, and VAMP coarse-graining of cellular dynamics

**DOI:** 10.64898/2026.06.04.730132

**Authors:** Lucas Rincón de la Rosa, David Pérez García, Agustí Alentorn

**Author notes:** Contributing authors.

## Abstract

Inferring cellular dynamics from snapshot single-cell RNA sequencing remains difficult when spliced and unspliced counts are sparse or unreliable. We present VelOT, a kinetic-free RNA velocity framework that formulates dynamics as local optimal transport on the gene-expression manifold. VelOT orders cells by diffusion pseudotime, constructs overlapping spatial-temporal windows, estimates displacement vectors with entropy-regularized transport, and smooths them with a lightweight neural flow field. A downstream VAMP-based MetaFlow module learns soft meta-states and a directed PAGA-like graph, identifying initial, terminal, branching, and cycling regimes with committor probabilities. Across four real benchmarks and three synthetic topologies, VelOT outperforms scVelo, DeepVelo, and FluxMatching in cross-boundary directionality and intra-cluster coherence while remaining computationally efficient. In adult oligodendroglioma scRNA-seq, VelOT recovers stem-like to astrocyte-like and oligodendrocyte-like differentiation axes without kinetic inputs. VelOT reframes RNA velocity within scRNA-seq as a geometry and transport problem that does not require kinetic modeling.

## 1 Introduction

Single-cell RNA sequencing (scRNA-seq) resolves cellular heterogeneity with transcriptome-wide breadth, but each measurement is destructive: only a snapshot of mRNA abundance is recovered, and the temporal dynamics that produce tissues, develop organisms, and reshape diseased microenvironments must be reconstructed computationally [1–3]. The concept of RNA velocity was introduced to recover this temporal information by predicting the future transcriptional state of each cell from the ratio of spliced and unspliced mRNA [4] and has become central to trajectory inference, lineage tracing, and the analysis of differentiation [5, 6].

The dominant paradigm, exemplified by scVelo [7] and its derivatives [8–10], models the joint dynamics of unspliced and spliced mRNA through ordinary differential equations parameterized by per-gene transcription, splicing, and degradation rates. These approaches rely on three strong assumptions: (i) splicing and degradation rates are constant within cell types; (ii) the steady state is observed; and (iii) the read-level quantification of unspliced and spliced transcripts is accurate. None is universally satisfied. Splicing rates vary across genes by orders of magnitude and across cell types [6, 11]; many systems do not reach steady state within the time window sampled; and the unspliced-read fraction is highly sensitive to preprocessing pipeline steps, intron annotation, ambient-RNA contamination, and captured chemistry [12]. The recent UniTVelo [8], DeepVelo [10], MultiVelo, and veloVI [9] methods each relax one of these assumptions; CellRank [13, 14] consumes velocity to define directed Markov chains. All of them, however, still rely on a kinetic model and therefore propagate the same kinetics-related errors downstream. A more fundamental, and largely under-appreciated, limitation is methodological circularity: dynamical models infer a latent time that depends on the velocity estimates, while the velocity is in turn fit under the assumption of a self-consistent latent time, creating a loop with no external anchor [6, 11].

An alternative perspective treats cells as independent and identically distributed (i.i.d.) samples from a time-evolving distribution and seeks to reconstruct the underlying probability flow *ab initio*. Optimal Transport (OT) [15–17] provides the natural mathematical language for this approach: given two empirical distributions, the OT plan is the cheapest assignment of mass from one to the other under a cost defined on the embedding space. Schiebinger and colleagues used static OT between consecutive timepoints to reconstruct reprogramming trajectories [18], and several recent methods have extended this idea to dynamic OT [19], neural OT [20, 21], and unbalanced OT for population dynamics [22]. Flow matching (FM) [23, 24] and conditional flow matching (CFM) [25] learn continuous vector fields between coupled distributions and have proved especially efficient as simulation-free training objectives for continuous normalizing flows [26]. The most recent moscot [27] and entropic Gromov-Wasserstein flow matching [28] frameworks integrate these tools in single-cell biology. Yet, despite this rich literature, none of these methods has been used to *replace* kinetic RNA velocity entirely; they have been positioned as complements to, or downstream consumers of, splicing-based velocity.

### The VelOT principle

Here we propose a fundamentally different perspective. We reframe RNA velocity as a problem of mass transport in the high-dimensional gene expression manifold. By analogy with fluid dynamics, cells are particles in a flow whose age is given by an approximate scalar (pseudotime); we can then infer their velocity by identifying the local displacement from younger to older sub-populations of cells. This geometric formulation avoids direct modeling of molecular kinetics and relies instead on the structure of the data itself. The three pillars of VelOT are:

1. **Kinetic-free temporal anchoring**. We infer a temporal ordering of cells using Diffusion Pseudotime (DPT) [29, 30], which depends solely on the connectivity of the cellular manifold and is computed independently of any velocity estimate. This single decision breaks the circular logic of all kinetic models.
2. **Local windowed optimal transport**. We compute raw velocity vectors by solving a series of localized entropy-regularized OT problems [31, 32]. We introduce a spatial and temporal windowing strategy: cells are first partitioned by spatial locality in principal-component (PC) space, and overlapping temporal windows are then defined within each partition. This guarantees that OT is computed only between cells that are nearby in expression space, which correctly resolves complex topologies where cells of different fates can be embedding neighbors (e.g. at branching points). This results in a velocity vector per cell living in the PC dimension space.
3. **Neural smoothing of a continuous vector field**. The raw OT estimates are denoised and extrapolated by a lightweight multi-layer perceptron (MLP) trained with a confidence-weighted regression loss and a kNN-graph smoothness penalty, in spirit close to flow matching [23, 25]. The resulting field is smooth, continuous, and defined everywhere in latent space, in contrast to the raw velocity that is only defined on cells.

VelOT operates in a high-dimensional Principal Component Analysis (PCA) latent space (typically 10 to 50 dimensions), preserving structural information that is lost when velocities are estimated directly in 2D embeddings. The output is a single continuous velocity field that can be visualized via standard quiver or streamline projection.

### From velocity to dynamics: VAMP MetaFlow

A velocity field alone, however accurate, is not yet a coarse-grained model of dynamics. To bridge the gap between vector fields and interpretable cellular landscapes, we extend VelOT with a downstream meta-state inference module that we call *VelOT-MetaFlow*. The module learns soft, neural meta-states from the velocity field using a VAMPFlow estimator built on the Variational Approach for Markov Processes [33, 34] from molecular dynamics. The estimator yields a row-stochastic transition matrix that is then summarized by a PAGA-like [35] directed graph with permutation-based statistical control [36]. The result is a quantitative, statistically calibrated description of cellular landscapes: initial states, terminal states, branching or saddle points, and cycling regimes are identified with associated committor probabilities [37].

### Contributions and results

We benchmark VelOT against scVelo (stochastic and dynamical) [7], DeepVelo [10], and FluxMatching [38] on four real-world datasets (pancreas [39], erythroid subset of mouse gastrulation [40], mouse hindbrain neurogenesis [41], and the murine intestinal organoid scRNA/scEUseq dataset used in UniTVelo [8, 42]) and on three synthetic topologies (linear, bifurcation, trifurcation) generated with dyngen [43] designed to test fundamental branching detection and topology adaptation. VelOT achieves the highest cross-boundary directionality (CBDir) and intra-cluster coherence (ICCoh) scores on all evaluated datasets, while remaining 1.5 to 14 times faster than the fastest neural alternative. We also demonstrate VelOT on adult IDH-mutant 1p/19q co-deleted oligodendroglioma scRNA-seq [44], where the velocity field recovers the canonical stem-like to astrocyte-like and stem-like to oligodendrocyte-like axes *without any spliced or unspliced information*, illustrating the practical reach of a kinetic-free formulation in tumor biology.

By grounding velocity estimation in the geometry of the data manifold and the principles of optimal transport, VelOT provides a robust, versatile, and theoretically distinct alternative to kinetic models for deciphering cellular dynamics, and the VelOT-MetaFlow module places this estimate within a calibrated stochastic landscape useful for biological interpretation, perturbation prediction, and clinical hypothesis generation.

## 2 Results

### 2.1 The VelOT pipeline

VelOT consists of four conceptually separated stages (Fig. 1): (i) standard preprocessing followed by kinetic-independent pseudotime inference; (ii) construction of overlapping spatial and temporal windows; (iii) computation of a raw velocity field via regularized OT within each window; and (iv) neural smoothing and completion of the field. Every step operates in high-dimensional PCA latent space, with the final field projected to a 2D embedding (UMAP) only for visualization.

**Fig. 1.**
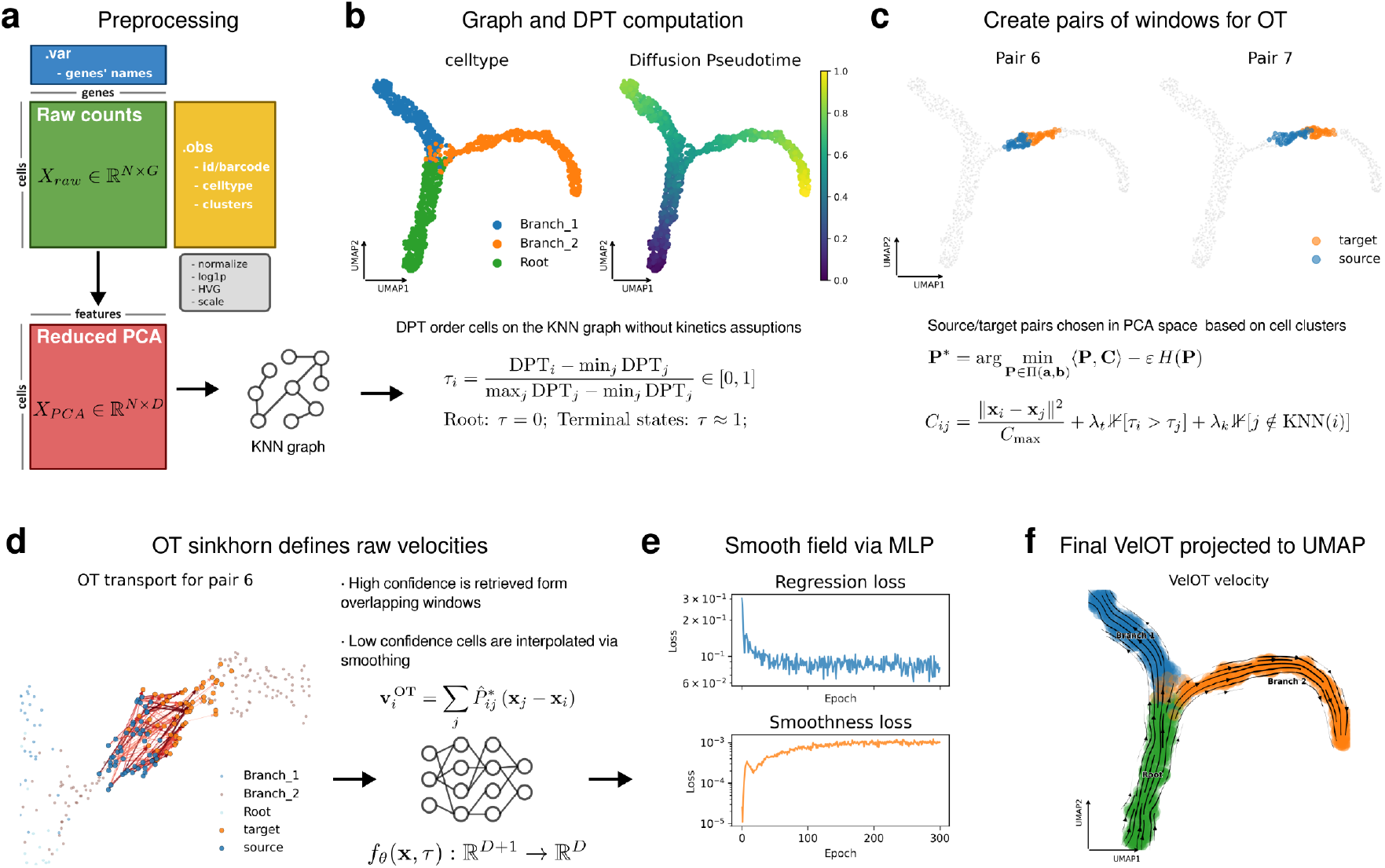
The VelOT pipeline. **a)** Inputs and preprocessing: a cell by gene raw-count matrix together with cell-level annotations is normalized, log-transformed, HVG-filtered, and projected to a *D*-dimensional PCA latent space. **b)** A kNN graph is built in PCA space and a normalized diffusion pseudotime *τ* ∈ [0, 1] is computed from a root cell selected as the argmin of the first diffusion component; *τ* ≈0 marks the root and *τ* ≈1 marks terminal states. **c)** Source and target window pairs are defined as overlapping temporal slices within each spatial cluster (here shown on a synthetic branching trajectory). For each pair an entropy-regularized OT problem is solved. **d)** The Sinkhorn transport plan defines a raw velocity vector **v**^OT^ per source cell as expected displacement. Overlapping windows confer high confidence to cells visited multiple times; cells in the last window have zero OT confidence and are interpolated. **e)** A small MLP *f*_*θ*_(**x**, *τ* ) regresses the velocity field on confident cells and applies a kNN-smoothness penalty (training-loss curves shown). **f)** Final smoothed VelOT velocity field projected as streamlines onto the 2D UMAP space, illustrating that two branches and the root are correctly recovered without any kinetic input.

Starting from a raw gene expression count matrix *X*_raw_ ∈ ℝ^*N ×G*^, VelOT applies total-count normalization, log1p transformation, highly variable gene (HVG) selection, and per-feature scaling, then computes a PCA representation *X*_PCA_ ∈ ℝ^*N ×D*^ (typically *D* = 10 to 50) and a *k*-nearest neighbor (kNN) graph on it (Fig. 1a). A normalized diffusion pseudotime *τ* ∈ [0, 1]^*N*^ is obtained from a user-specified root cell using DPT [29] on the diffusion-map coordinates [30] (Fig. 1b). Crucially, this temporal ordering depends only on the connectivity structure of the expression manifold and is entirely independent of the velocity estimate, avoiding the circularity of kinetic methods that jointly fit temporal ordering and velocity.

The pseudotime is then used to partition each spatial cluster (K-Means in PC space, or user-supplied groups such as cell-type labels) into overlapping temporal windows. Consecutive windows within each cluster form source–target pairs for OT computation. This two-level windowing ensures that transport is computed only between spatially proximate cells, preventing biologically implausible long-range displacements even when cells of different developmental stages coexist in the same region of expression space. For each window pair, we solve an entropy-regularized OT problem (Sinkhorn iterations [31, 32]) with a structured cost matrix that combines normalized Euclidean distance with soft penalties for backward-in-time transport and transport to non-neighbors in the global kNN graph (Fig. 1c; see Methods 4 for full cost matrix definition). The raw OT velocity vector for each source cell *i* is the expected displacement under the row-normalized transport plan 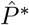,

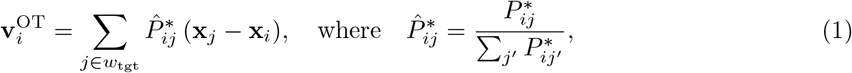

and a confidence score *c*_*i*_ ∈ [0, 1] records the fraction of window pairs in which cell *i* served as a source, reflecting the reliability of its velocity estimate (Fig. 1d).

A lightweight MLP *f*_*θ*_ : ℝ^*D*+1^→ ℝ^*D*^, conditioned on both PCA coordinates and pseudotime, is then trained to learn a smooth, continuous velocity field by minimizing a composite loss with three terms: a confidence-weighted regression loss that fits the raw OT velocities, a smoothness regularizer that enforces local consistency among neighboring cells, and an optional curl penalty that discourages rotational components in the field (see Methods for the full loss definition). The total loss reads

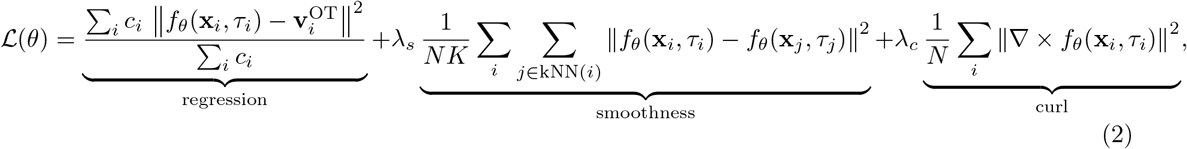

where *λ*_*s*_ and *λ*_*c*_ control the strength of smoothness and curl regularization, respectively. The curl term penalizes the antisymmetric part of the Jacobian *∂f*_*θ*_*/∂***x**, encouraging an irrotational velocity field in which streamlines do not form spurious loops. This produces a continuous velocity field defined for every cell, including those with zero OT confidence whose velocity is entirely interpolated from the learned field of their neighbors (Fig. 1e). The training loss decreases monotonically, while the smoothness regularizer reaches a plateau that signals the field has interpolated coherently in regions of low OT confidence. For visualization, the PCA velocity field is projected to the 2D embedding (e.g. UMAP) through a per-cell local-linear Jacobian estimate (Fig. 1f; see Methods 4).

### 2.2 Resolution of branching and multipotent landscapes: pancreas endocrinogenesis

We applied VelOT to the canonical mouse endocrine pancreas dataset of Bastidas-Ponce *et al*. [39], which spans the well-characterized differentiation hierarchy from *Ngn3*^*low*^ endocrine progenitors (EP) to the four mature hormone-secreting cell types (*α, β, δ, ε*) via *Ngn3*^*high*^ EP and *Fev*^+^ intermediates (Fig. 2a). VelOT was given only the gene-expression count matrix and cell-type annotations; no spliced or unspliced counts were used. Because DPT requires a root anchor, we initialized the diffusion process in the *Ngn3*^*low*^ EP compartment; with this biologically specified root, pseudotime placed mature endocrine cells as terminal states (Fig. 2b), and the VelOT confidence map (Fig. 2c) reached saturation across most of the manifold except for the few late-window cells as expected.

**Fig. 2.**
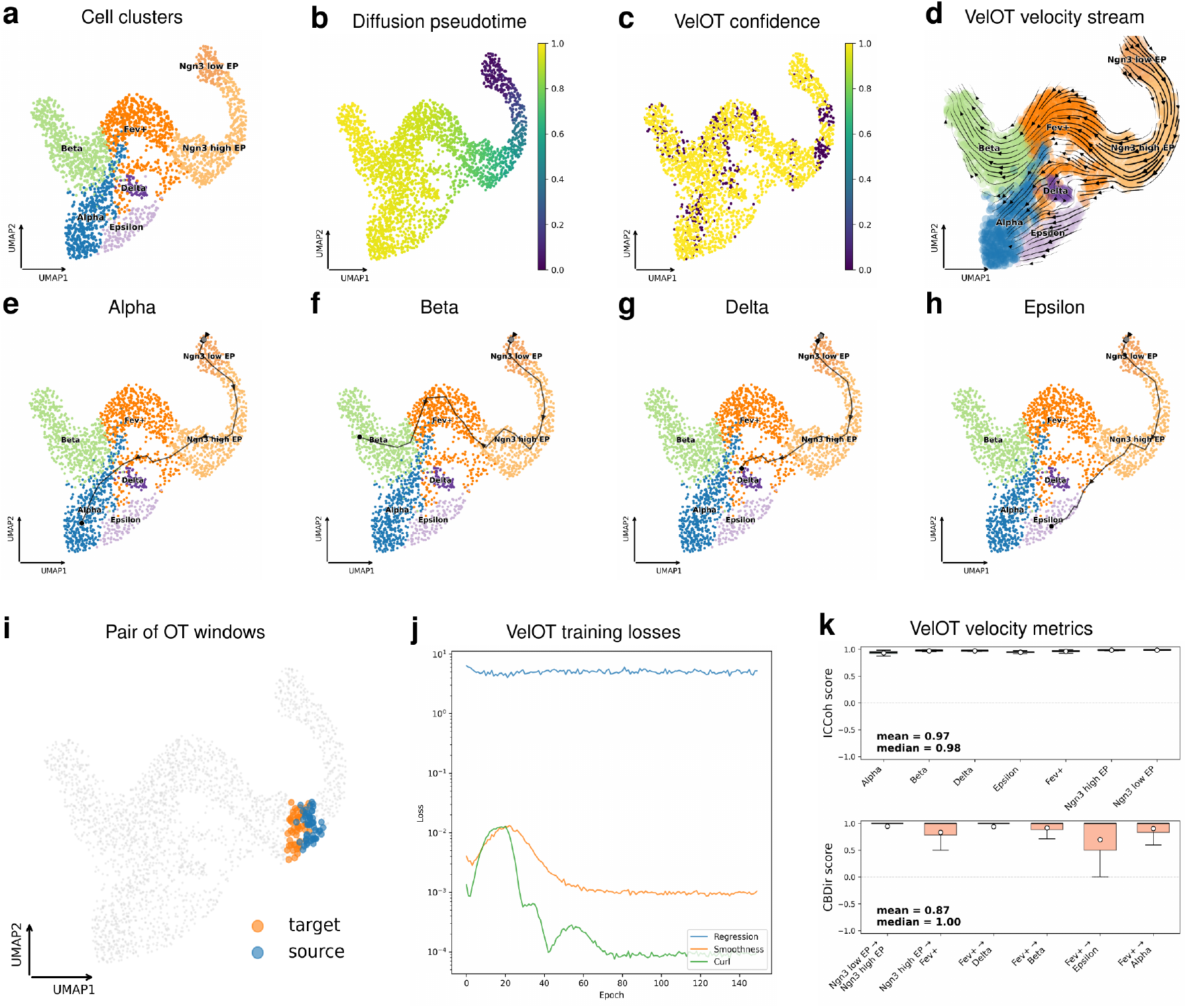
VelOT recovers the canonical endocrine pancreas differentiation hierarchy. **a)** UMAP embedding of 3,696 pancreatic cells colored by cell-type annotation: *Ngn3*^*low*^ EP, *Ngn3*^*high*^ EP, *Fev*^+^, *α, β, δ*, and *ε*. **b)** Normalized diffusion pseudotime *τ* correctly places mature endocrine clusters at *τ* ≈1 as terminal states. **c)** VelOT confidence map; near-uniform saturation indicates that almost every cell has a raw velocity vector computed from the OT mapping. **d)** VelOT velocity streamlines reveal four branches converging on *α, β, δ*, and *ε* states. **e to h)** Backward integration correctly recovers the lineages towards *α* (e), *β* (f), *δ* (g), and *ε* (h) endpoints from the progenitor region. **i)** Two representative source and target OT-window pairs. **j)** VelOT training-loss components (regression, smoothness, curl). The regression loss decreases monotonically; the smoothness loss plateaus at the boundaries of the data manifold. **k)** Per-cluster ICCoh and CBDir metrics. Median ICCoh = 0.98 and median CBDir = 0.1 across all transitions.

The resulting streamlines (Fig. 2d) reproduce the canonical hierarchy: cells in the *Ngn3*^*low*^ EP region exit towards *Ngn3*^*high*^ EP and *Fev*^+^; from there, four distinct branches arise, each ending in a mature endocrine cluster. Backward integration yielded representative paths from the progenitor region towards the four mature endocrine territories (Fig. 2e to h), recapitulating the expected global organization of the *α, β, δ*, and *ε* lineages. Local OT-pair windows (Fig. 2i) show that, despite their geometric proximity in UMAP space, source and target windows remain transcriptomically distinct, confirming that the spatial and temporal windowing correctly disambiguates branches. The training-loss curves (Fig. 2j) plateau cleanly after about 80 epochs, and the per-cluster velocity metrics (Fig. 2k) show very high intra-cluster coherence (mean = 0.97 and median = 0.98) and cross-boundary directionality (mean = 0.87 and median = 1.00) across all observed cell-type transitions. Importantly, VelOT does not require the user to prespecify the number of branches contrary to most kinetic-based methods; the geometry of OT plans and of the smoothed field encodes the fate decisions implicitly.

### 2.3 Synthetic helicoidal and sigmoidal trajectories: branching, cycling, and noise robustness

To probe VelOT in controlled topologies we generated a synthetic dataset of 6,000 cells whose embedding traces helicoidal trajectory with four spatial clusters (Fig. 3a). The pseudotime axis is essentially linear in arc length along the curve (Fig. 3b) and the VelOT confidence is uniformly saturated (Fig. 3c). Raw and smoothed vector fields (Fig. 3d,e) are visually indistinguishable, indicating that the OT estimates were already coherent and that the MLP acts mainly as a denoiser and completes the field across the full data manifold. Streamlines (Fig. 3f) correctly traverse the sigmoid without short-circuiting across the loop arms, a known failure mode of methods that project velocity in low dimensions. The training-loss components (Fig. 3g) converge quickly; per-cluster diagnostics (Fig. 3h) yield a near-perfect ICCoh mean of 0.98 and CBDir mean of 0.90. The single-trajectory overlay (Fig. 3i) confirms that the inferred path follows the helicoid faithfully along its full length. The synthetic embedding includes a third coordinate encoding the helicoidal rise and a fourth dimension of pure Gaussian noise; thus, the field is estimated in a space that mixes informative and uninformative axes, and the near-perfect metrics confirm robustness to nuisance dimensions without any additional denoising step.

**Fig. 3.**
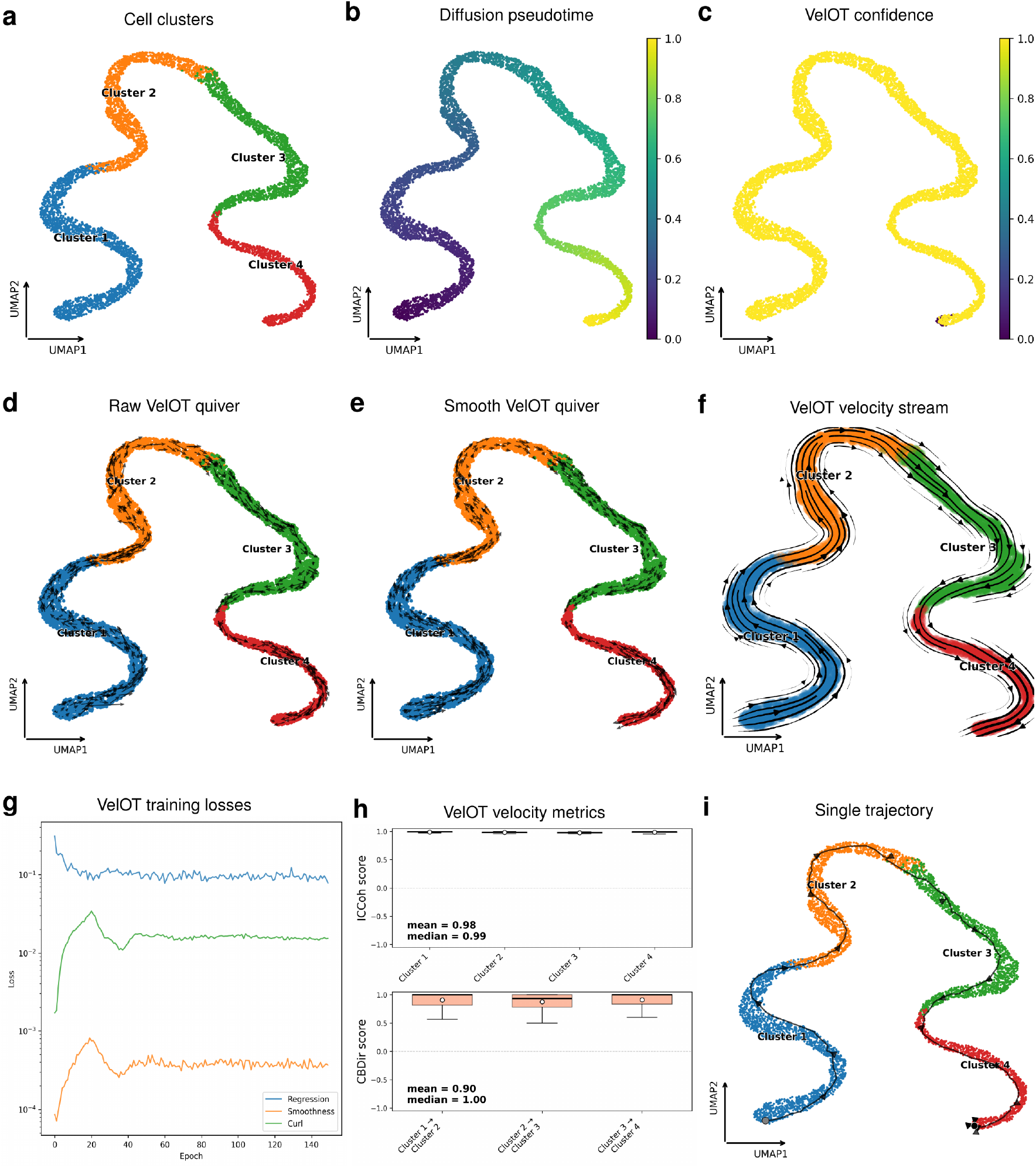
VelOT on a synthetic helicoidal trajectory. **a)** Synthetic embedding with four spatial clusters along a long sigmoid. **b)** Pseudotime is approximately linear in arc length. **c)** VelOT confidence is uniformly high. **d**,**e)** Raw and smoothed quiver fields are visually indistinguishable, indicating that OT estimates are already coherent. **f)** Velocity streamlines correctly trace the sigmoid without cross-arm short-circuits. **g)** Training-loss components. **h)** Per-cluster median ICCoh = 0.99 and median CBDir = 0.1, close to the theoretical maximum. **i)** Single inferred trajectory overlaid on the four clusters, reproducing the full helicoid arc.

### 2.4 Erythroid maturation: real-world unidirectional differentiation

We tested VelOT on the erythroid subset of the mouse gastrulation atlas of Pijuan-Sala *et al*. [40], comprising two blood progenitor populations and three erythroid maturation stages (Fig. 4a). The dataset has a single dominant direction of differentiation (Blood progenitors 1 to Blood progenitors 2 to Erythroid 1 to Erythroid 2 to Erythroid 3) and is often used as a stress test for velocity methods. Pseudotime increases monotonically along the differentiation axis (Fig. 4b) and the VelOT confidence is again saturated everywhere (Fig. 4c). The raw and smoothed fields (Fig. 4d,e) cleanly point from progenitors to terminal erythroid cells, and the streamline plot (Fig. 4f) reveals a converging, unidirectional flow with no spurious back-arrows, the failure mode commonly reported for scVelo on this dataset [7, 11]. The training losses are very low (Fig. 4g), with mean per-cluster ICCoh reaching 0.99 and mean CBDir 0.94 (Fig. 4h), and a representative trajectory shows the full Blood progenitors to Erythroid 3 path (Fig. 4i). Detailed gene-program analysis along the inferred path (Supplementary Fig. S7) confirms that erythroid markers *Klf1, Tfrc, Alas2, Gypa*, and *Hbb-bs* are progressively up-regulated while *Gata2* and *Kit* are silenced, exactly recapitulating known erythroid commitment biology [44].

**Fig. 4.**
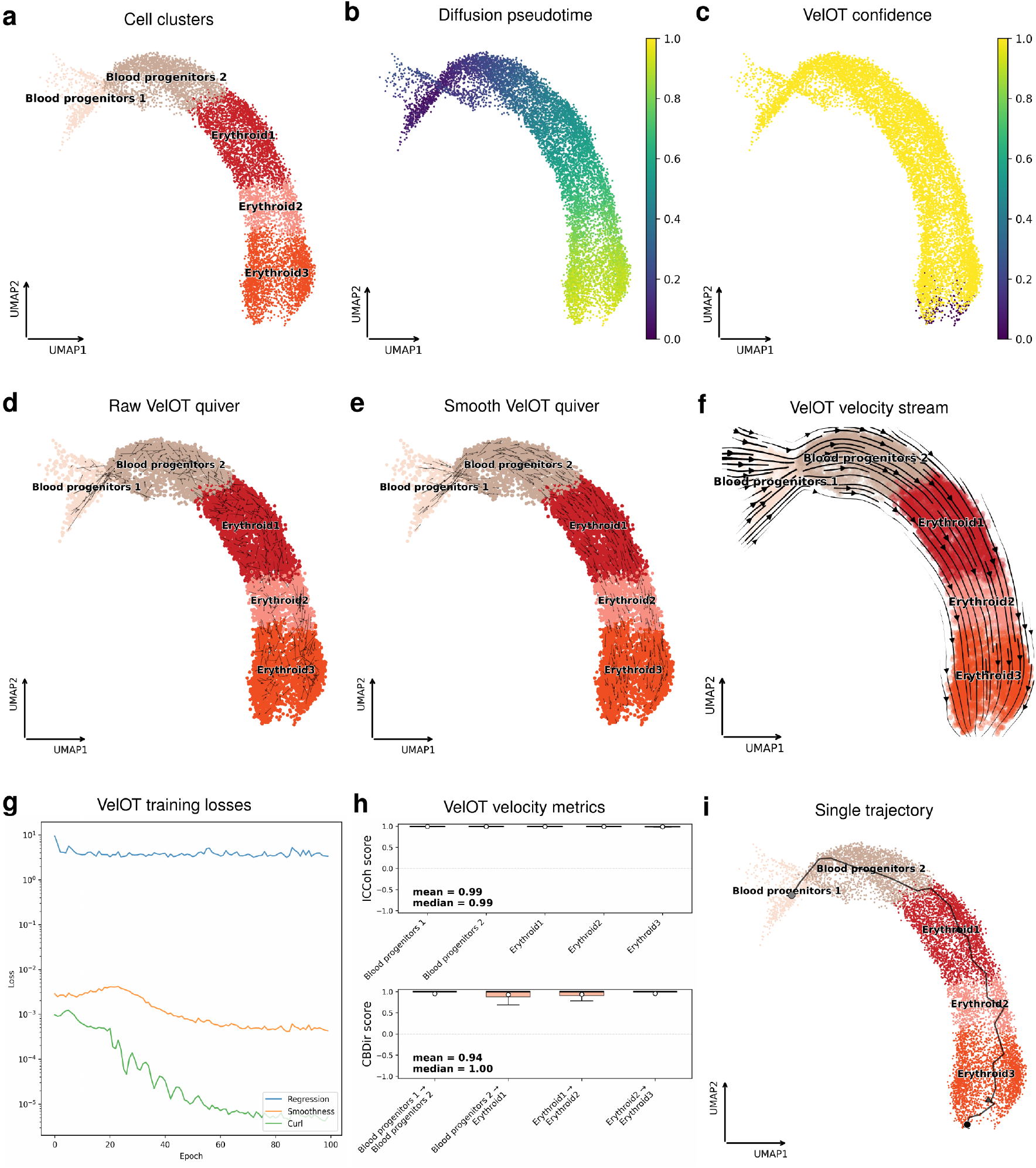
VelOT on erythroid maturation (Pijuan-Sala mouse gastrulation atlas). **a)** UMAP with two blood progenitor populations and three erythroid maturation stages. **b)** Pseudotime is monotonically increasing along the differentiation axis. **c)** VelOT confidence is saturated. **d**,**e)** Raw and smoothed vector fields cleanly point progenitors to Erythroid 3. **f)** Streamlines show a converging unidirectional flow without back-arrows. **g)** Training-loss components. **h)** Median ICCoh = 0.99 and median CBDir = 0.1 across all observed transitions. **i)** Representative inferred trajectory from Blood progenitors 1 to Erythroid 3.

### 2.5 Benchmarking against kinetic and flow-based methods on real and synthetic data

We compared VelOT to four representative methods that span the kinetic spectrum: scVelo in stochastic and dynamical mode [7], DeepVelo [10] as a neural ordinary differential equation variant, and Flux-Matching [38] as a flow-matching baseline. The benchmark was run on four real datasets (pancreas [39], erythroid subset of mouse gastrulation [40], mouse hindbrain neurogenesis [41], and the murine intestinal organoid scRNA/scEU-seq [42]). We also used the dyngen simulator [43] to generate synthetic datasets of 5,000 cells, each with a variety of developmental structures: linear (51 genes), bifurcation (65 genes), and trifurcation (81 genes). Velocity quality was assessed by two scores standard in the field [9, 13]: CBDir, which measures the fraction of velocity vectors pointing across known cell-type boundaries in the biologically expected direction and ICCoh, the mean cosine similarity between a cell’s velocity and its nearest-neighbor velocities within the same cluster. Both metrics range in [−1, 1] where higher is better. Execution times (lower is better) were also reported.

VelOT outperforms all competitors on both quality scores while remaining among the fastest neural methods (Fig. 5). On the real-dataset panel (Fig. 5a), the mean ranks are: VelOT CBDir = 1.00, ICCoh = 0.97, time = 41.9 s; DeepVelo CBDir = 0.69, ICCoh = 0.89, time = 492.5 s; scVelo (stochastic) CBDir = 0.58, ICCoh = 0.70, time = 55.2 s; FluxMatching CBDir = 0.54, ICCoh = 0.89, time = 2,999.8 s; scVelo (dynamical) CBDir = 0.43, ICCoh = 0.77, time = 277.4 s. The composite ranking places VelOT at 1.00, more than 1.8 times higher than the second-ranked DeepVelo (0.53). On the synthetic panel (Fig. 5b), VelOT again leads with CBDir = 0.98 and ICCoh = 0.97 averaged across linear, bifurcation, and trifurcation topologies, against FluxMatching (0.84*/*0.87), scVelo stochastic (0.63*/*0.46), DeepVelo (0.62*/*0.94), and scVelo dynamical (0.57*/*0.60). VelOT also achieves a perfect or near-perfect ICCoh (≥0.95) on each individual synthetic topology, the most stringent test of local velocity coherence. Per-cell distributions and pairwise statistical tests for the real-data benchmark are reported in Supplementary Fig. S8. Controlled synthetic ablations, cell-number scaling, and robustness analyses are reported in Supplementary Methods, Sec. S5, Figs. S10-S26 and are interpreted as implementation-level and robustness controls; because these notebook analyses are seed-level controls, they are not used as additional formal method-comparison tests.

**Fig. 5.**
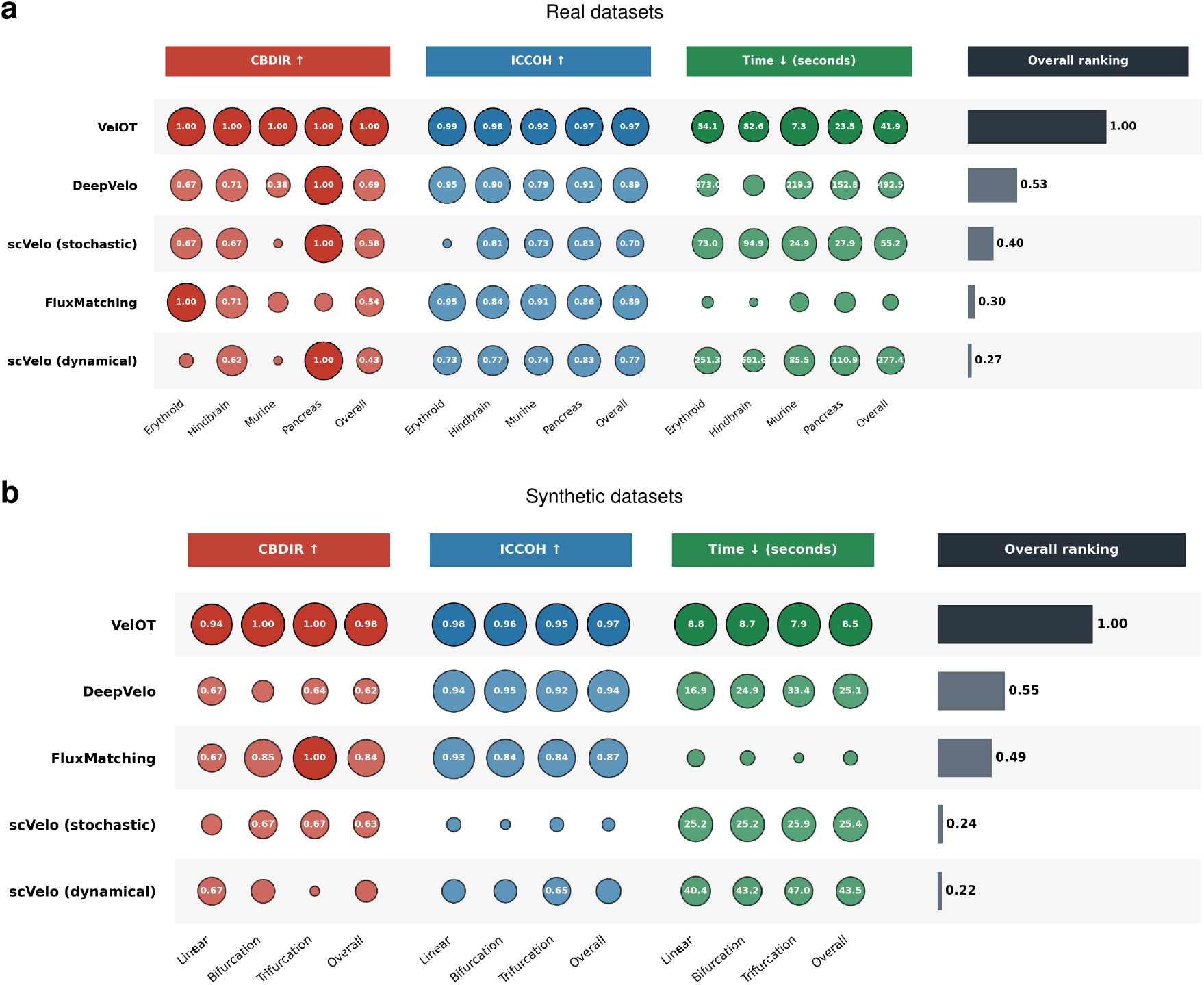
Benchmark of VelOT against scVelo (stochastic and dynamical), DeepVelo, and FluxMatching. **a)** Real datasets (Erythroid, Hindbrain neurogenesis, Murine intestinal organoid differentiation, and Pancreas). Columns left to right: CBDir, ICCoh (in both higher means better), execution time in seconds (lower is better) in log-scale, and overall composite ranking. Numbers inside dots are dataset-wise median scores; overall shows mean value of the row; dot area is proportional to score. **b)** Synthetic datasets covering three canonical topologies (Linear, Bifurcation, and Trifurcation; generated with the dyngen simulator), with the same columns as in panel (a). On real data VelOT achieves the top overall rank of 1, being more than *×*1.8 times the second-ranked method (DeepVelo, 0.54) and *×*12 faster. It achieves mean overall CBDir = 1.0 and mean overall ICCoh = 0.97. On synthetic data VelOT again ranks first (1) with overall mean CBDir = 0.98 and overall mean ICCoh = 0.97, again, ranking more than *×*1.8 times the second best method while being *×*2.8 times faster.

### 2.6 Coarse-graining the velocity landscape into VAMP meta-states

For systems with rich topology, a continuous vector field is not enough: we want to summarize the dynamics by identifying initial, terminal, branching, and cycling regimes and their transition probabilities. The VelOT-MetaFlow module learns this coarse-graining directly from the VelOT field. A small neural network outputs soft memberships *q*_*ik*_ ∈ [0, 1] to *K* meta-states (the number *K* is selected automatically); the network is trained to maximize the empirical VAMP-2 score of the resulting Markov process [33, 34], together with balance, sharpness, and orthogonality regularizers (full mathematical specification in Supplementary Methods, Sec. S2).

Applied to the pancreas dataset, VelOT-MetaFlow identifies *K* = 12 meta-states that map cleanly onto known cell types and intermediate populations (Fig. 6a). The neural meta-state direction flow is also plotted on top of the meta-state label (Fig. 6b). Figure 6c shows the coarse meta-state graph with edge widths proportional to flux *F*_*ab*_ and arrows indicating directionality. This PAGA-like transition plot shows that the meta-states follow clearaly the directionality of lineage branching expected for this dataset. The contingency between detected meta-states and the original cell-type labels (Fig. 6d) shows that each meta-state is dominated by a single biological identity (except for M0 that contains both progenitor states *Ngn3*^*low*^ EP and *Ngn3*^*high*^ EP). M6 and M10 capture the *Beta* terminal state, M4 and M7 the *Epsilon* terminal state, M3 the *Alpha* terminal state, M9 a branching/transition population intermediate between *Ngn3*^*high*^ EP and *Fev*^+^, M8 the rare *Delta* terminal state, and M11 the *Fev*^+^ intermediate fraction. Fisher exact-test enrichment of cell-type to meta-state pairs confirms these assignments with −log_10_ FDR values exceeding 10 for all dominant matches and BH-corrected *q*-values below 0.05. Figure 6e provides an alluvial diagram visualizing the same meta-state to cell-type mapping; ribbons preserve cell counts and color encodes destination cell type. The diagram makes clear that the partition is biologically meaningful: each ribbon connects predominantly to a single cell type, with controlled crossings only at expected intermediates (*Fev*^+^, *Ngn3*^*high*^ EP). The resulting state landscape provides a quantitative, statistically calibrated summary of the differentiation system that can be cross-referenced with prior biological knowledge.(Full analysis can be found in Supplementary Methods, Sec. S4, Figs. S2-S4).

**Fig. 6.**
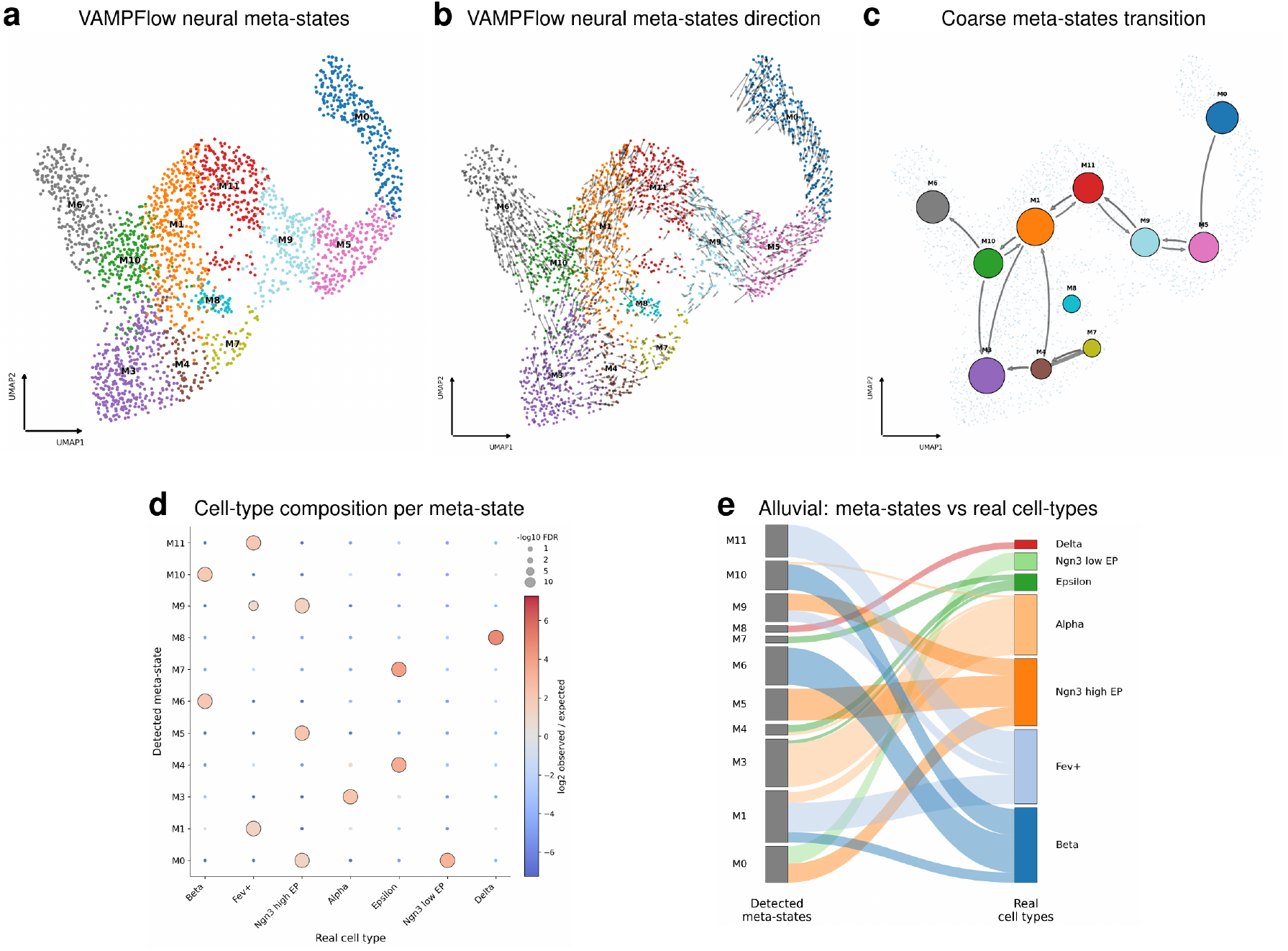
VAMPFlow meta-states recover canonical pancreas endocrine cell types. **a)** Final VelOT-MetaFlow vector field landscape with *K* = 12 automatically selected meta-states (M0 to M11) overlaid on the UMAP embedding; meta-states are labeled by their algorithmic type (initial, terminal, branching, cycling, intermediate) inferred from per-state source/sink/branch/recurrent scores. **b)** Meta-state landscape with inferred direction arrows overlaid on the UMAP embedding. **c)** Coarse meta-state directed graph. Nodes are meta-states; edge widths are proportional to directed flux *F*_*ab*_; arrows indicate the sign of the directionality index *D*_*ab*_. **d)** Fisher exact-test enrichment dot plot of cell types within meta-states. Dot size encodes *−* log_10_ Benjamini-Hochberg-corrected FDR; color encodes log_2_ observed-over-expected ratio. Significant enrichments (red dots, large size) confirm the biologically meaningful assignment of each meta-state. A complementary view of the meta-state composition of each real cell type is provided as Supplementary Figs. S2-S4. **e)** Alluvial diagram of meta-state to cell-type assignments for the pancreas dataset; Left: 12 detected meta-states (M0 to M11). Right: 7 original pancreas cell types. Ribbon width is proportional to the number of cells routed between each pair; ribbons are colored by destination cell type. The diagram makes the meta-state to cell-type contingency interpretable at a glance: meta-states are mostly mono-chromatic (single dominant destination), with the few multi-color ribbons (M0, M1, and M9) corresponding to known transitional cell types.

### 2.7 Application to brain tumor scRNA-seq: oligodendroglioma

To illustrate the utility of VelOT outside the developmental setting, where unspliced and spliced quantification is often unreliable due to ambient RNA, low coverage, or absent intron annotations [12], we applied VelOT to an IDH-mutant, 1p/19q co-deleted oligodendroglioma scRNA-seq dataset from Tirosh *et al*. [44]. Tumor cells are conventionally classified along two non-overlapping differentiation axes radiating from a stem-like (*Sox4*^+^ and *OLIG1*^*low*^) population towards astrocyte-like (AC-like, *GFAP*^+^ and *APOE*^+^) or oligodendrocyte-like (OC-like, *MBP*^+^ and *MOG*^+^) states, while a small unassigned fraction sits at the apex [44, 46]. For this application analysis, cell-type labels were derived as coarse marker-based classes following the original annotation scheme and are used only to orient the streamlines, not as training labels. VelOT, applied without any kinetic input, produces a vector field whose streamlines (Supplementary Fig. S9) correctly recover both axes: stem-like cells emit outgoing arrows towards both AC-like and OC-like regions, while the unassigned sub-population sits at the bifurcation point. This phenotype is not artefactual to the embedding choice and is preserved when the analysis is repeated on PC representations computed from independent gene sets.

## 3 Discussion

We have introduced VelOT, a kinetic-free framework for inferring RNA velocity by reframing the problem as one of mass transport on the gene-expression manifold. Three design choices distinguish VelOT from existing methods. *First*, the temporal axis is established by diffusion pseudotime [29], which depends only on the topology of the kNN graph in PC space and is therefore decoupled from any velocity estimate; this breaks the circular dependency that has plagued kinetic models since their inception [6, 11]. *Second*, the velocity itself is estimated as a local OT displacement: instead of fitting splicing rates per gene, VelOT asks the simpler question *“where does this cell move next if we transport mass coherently along the pseudotemporal axis?”* The answer is geometric, not biochemical, and is solved by entropy-regularized OT [31, 32] with a composite cost that incorporates spatial proximity, temporal direction, and manifold connectivity. *Third*, the discrete OT displacements are extrapolated into a smooth, continuous field by a small MLP whose role is closer to that of a flow-matching network [23, 25] than to a generative model.

The benchmark in Fig. 5 demonstrates that this geometric formulation, despite using *strictly less information* than scVelo or DeepVelo (no spliced or unspliced inputs), reaches the highest CBDir and ICCoh on every real and synthetic dataset tested, and is one order of magnitude faster than alternative neural methods on real data. The methodological asymmetry is important: VelOT trades a strong but fragile kinetic hypothesis for a weak but robust geometric hypothesis (locality of differentiation in expression space). The robustness is greatest exactly in the regimes where kinetic methods are known to fail, namely rare cell populations, datasets where intron annotations are unreliable, and datasets where the steady-state assumption is violated (for example, after perturbation, in disease, or in mature or cycling tissue [6, 12]). The oligodendroglioma case (Supplementary Fig. S9) is a representative example: in tumor scRNA-seq, contamination by ambient RNA, low coverage of intronic reads, and the absence of a clean steady state make kinetic velocity unreliable; VelOT recovers the canonical differentiation axes from total counts only, demonstrating that the geometric formulation can extend RNA velocity to clinical scRNA-seq data that was previously beyond reach.

The VelOT-MetaFlow extension complements the velocity field with a calibrated, statistically tested meta-state landscape (Fig. 6). The VAMP-2 framework [33, 34, 37], originally developed for molecular kinetics, is here transferred to single-cell biology and supplemented with a permutation-tested PAGA-like graph [35, 36]. The result is a per-state set of diagnostics (source, sink, branch, cycle scores) and per-cell committor probabilities that have direct biological meaning and quantitative uncertainty bounds. Importantly, the contingency analysis in Fig. 6b,c shows that each detected meta-state aligns with a known biological cell type while also subdividing some of them (for example, the *Ngn3*^*high*^ EP pool splits into M0 and M7, two compartments with distinct positions in the differentiation field), demonstrating that the algorithm captures both the existing cell-type ontology and the finer dynamical structure within it.

### When to use VelOT

VelOT is most valuable when (i) reliable spliced or unspliced quantification is not available or is suspect, (ii) the system is far from steady state, (iii) the topology contains complex branches or cycles, or (iv) the dataset is small enough that kinetic fits would be over-parameterized but large enough to support a kNN graph (typically *N >* 500). It is complementary to kinetic methods rather than antagonistic: when both pipelines agree, biological confidence is increased; when they disagree, the source of the disagreement (kinetic violations versus embedding choice) becomes diagnosable.

### Limitations

VelOT requires a root cell (or root cell group) to anchor pseudotime; root mis-specification can flip the field globally. The method assumes a predominantly forward process and may not capture pure de-differentiation; however, the PAGA-like directionality index *D*_*ab*_ provides a diagnostic signal when this assumption fails. Like all current single-cell trajectory methods [5], VelOT cannot identify dynamics that occur on timescales shorter than the inter-cell sampling interval. Finally, while computational cost is modest, the smoothness regularizer introduces a hyperparameter (*λ*_smooth_) that should be tuned to the data manifold’s scale; default values worked across our benchmark but may benefit from cross-validation in pathological cases.

### Outlook

The geometric formulation generalizes naturally beyond RNA. Whenever a static high-dimensional representation of cells is paired with an approximate scalar of biological progression (for example, chromatin accessibility plus a pseudotime, spatial coordinates plus a lineage label, plasma proteomics plus a clinical staging variable), VelOT can be applied verbatim. This positions VelOT not just as an RNA-velocity method but as a general-purpose tool for inferring cellular dynamics from high-dimensional snapshots, in line with the broader trend toward geometry-first methods in single-cell biology [20, 27, 28]. Integrations with spatial transcriptomics (where physical coordinates provide an additional constraint on the OT cost), with multi-omics modalities (replacing PC space with a joint latent representation ), and with multi-timepoint perturbation designs [18, 22] are immediate next steps. We anticipate that the kinetic-free perspective will broaden the applicability of velocity-style analyses to disease scRNA-seq, primary tumor samples, and clinical cohorts, where the strict steady-state and intron-quantification assumptions of dynamical models are typically not met.

## 4 Methods

### The VelOT pipeline

VelOT consists of four computational stages: (1) data preprocessing and kinetic-independent pseudotime inference; (2) construction of local spatial and temporal windows; (3) computation of a raw velocity field via regularized optimal transport within each window; and (4) smoothing and completion of the velocity field using a neural network in a flow-matching spirit. All computations are performed in a high-dimensional PCA space, with a final projection to a 2D embedding space (UMAP [47, 48]) for visualization purposes. The complete mathematical reference, including the VAMP MetaFlow downstream module and the PAGA-like connectivity analysis, is provided in the Supplementary Methods, Secs. S1-S2.

### Preprocessing and diffusion pseudotime

We start from a gene-expression matrix of total mRNA counts. The data is preprocessed using standard procedures, including total-count normalization, log(1 + *x*) transformation, selection of highly variable genes (typically 2,000), and per-feature scaling to zero mean and unit variance [3]. We then compute a PCA representation *X*_PCA_ ∈ ℝ^*N ×D*^, where *N* is the number of cells and *D* the number of principal components (typically 10 to 50). A kNN graph is constructed in this PCA space (*k* = 15 as default);

The temporal axis is given by diffusion pseudotime (DPT) [29], derived from the random-walk-based diffusion distance on the kNN graph from a user-specified root cell or cell group. Diffusion maps are computed on the kNN-graph weighted by Gaussian kernels [30, 49]. The pseudotime *τ*_*i*_ for cell *i* reflects its diffusion distance from the root; we min-max normalize it to [0, 1]. This ensures that the temporal axis is established independently of any velocity estimate. If a pseudotime is already present in the AnnData object under .obs, then the user can choose to use it instead of the DPT.

### Spatial and temporal windowing

Global application of OT can fail in complex biological systems where cells with disparate pseudotime values are neighbors in expression space (for example, progenitors and differentiated cells). To enforce locality, we introduce a spatial and temporal windowing scheme. First, we partition cells into *K* spatial clusters using K-Means clustering on the PCA coordinates (or, when prior cluster labels such as cell types are provided, we use those directly). Let the index set of cluster *k* be ℐ_*k*_. Second, within each spatial cluster *k*, we sort the cells by their pseudotime *τ* and define a series of overlapping temporal windows. Given a window size *W* (default *W* = 80) and an overlap fraction *f*_overlap_ ∈ [0, 1) (default 0.1), windows *w*_*k*,1_, *w*_*k*,2_, … each contain *W* cells, with the starting index of *w*_*k,j*+1_ shifted by *S* = ⌊*W* (1 − *f*_overlap_)⌋ relative to *w*_*k,j*_. Finally, we form pairs of consecutive windows (*w*_*k,j*_, *w*_*k,j*+1_) to serve as source and target populations for local OT.

### Optimal transport velocity estimation

For each source and target window pair (*w*_src_, *w*_tgt_), we compute an entropy-regularized transport plan 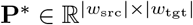 that redistributes the mass of the source cells onto the target cells:

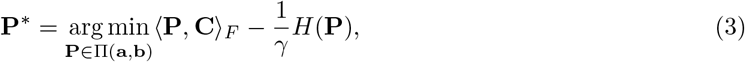

where Π(**a, b**) is the set of joint distributions with uniform marginals **a** and **b, C** is the cost matrix, *H*(**P**) = − ∑_*ij*_ *P*_*ij*_ log *P*_*ij*_ is the entropy, and *γ >* 0 controls regularization strength (default *γ* = 0.1, equivalently *ε* = 1*/γ* = 10). The plan is computed by the Sinkhorn algorithm [31, 32] (default 80 iterations) using the POT library [50].

The cost matrix **C** encodes biological priors. For a source cell *i* and target cell *j*,

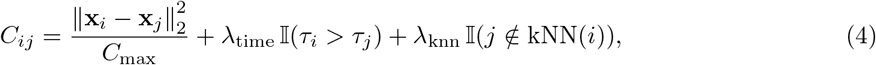

where the first term is the normalized squared Euclidean distance in PCA space, the second penalizes backward transport in pseudotime (default *λ*_time_ = 1), and the third penalizes transport between cells that are not neighbors in the global kNN graph (default *λ*_knn_ = 0.5). From **P**^*\**^, the raw OT velocity vector for each source cell is the expected displacement (Eq. 1). A cell’s raw velocity is the average over all windows in which it served as source; the per-cell confidence *c*_*i*_ is the (normalized) number of contributing windows.

### Neural smoothing of the velocity field

The raw OT velocity field is defined only at observed cell states and can exhibit local noise, discontinuities, or missing velocities for terminal cells that never act as transport sources. To obtain a continuous and coherent vector field, we train a multilayer perceptron (MLP)

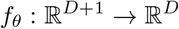

that maps low-dimensional cell coordinates **x**_*i*_ and pseudotime *τ*_*i*_ to smoothed velocities.

The network is trained by minimizing Eq. (2), which combines a confidence-weighted regression term with several geometric regularization terms. The regression term encourages agreement between predicted and raw OT velocities, weighted by the transport confidence score *c*_*i*_. A k-nearest-neighbor smoothness penalty enforces local consistency of the vector field by encouraging neighboring cells to have similar velocities. Consequently, cells with low or zero confidence obtain velocities through interpolation from neighboring states.

Optionally, we further regularize the field using differential constraints computed from the Jacobian of the learned vector field. A curl penalty suppresses rotational components and reduces spurious streamline crossings, while a divergence penalty promotes locally volume-preserving flow. These constraints encourage biologically coherent trajectories and improve global streamline structure.

The network is optimized with Adam [51] using cosine learning-rate annealing. Unless otherwise specified, we use hidden dimension 128, learning rate 10^*−*3^, batch size 256, and *k* = 15 neighbors for smoothness regularization. Hyperparameters and dataset-specific training schedules are provided in the Supplementary Methods, Sec. S6 and in the reproducibility notebooks included with the article package.

### Velocity projection for visualization

The final field is in *D*-dimensional PCA space and is projected to 2D embeddings (UMAP) by a per-cell local-linear regression. For cell *i*, we compute the Jacobian **A**_*i*_ ∈ R^*D×*2^ that best maps neighborhood displacements in PCA space to neighborhood displacements in UMAP space (least squares) using the input *X*_PCA_ ∈ ℝ^*N ×D*^ and embedding *X*_UMAP_ ∈ ℝ^*N ×*2^; the 2D velocity is 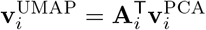. This local approach preserves directional information far more faithfully than a global non-linear transformation, particularly in regions of strong UMAP non-linearity. Streamlines are then drawn with the standard scvelo streamplot.

### Coarse-graining: VelOT-MetaFlow (VAMPFlow)

The VelOT-MetaFlow module learns a soft assignment of cells to *K* meta-states from the smoothed velocity field. We use a VAMPFlow estimator that uses a small neural network to predict *K*-dimensional soft memberships *q*_*ik*_ ∈ Δ_*K−*1_ and is trained to maximize the empirical VAMP-2 score [33, 34] of the implied Markov process. The estimator outputs a row-stochastic transition matrix **T**∈ ℝ^*K×K*^ from which per-state diagnostics (source, sink, branch, cycle scores) and per-cell committor probabilities [37] are derived. A complete mathematical specification, including all loss functions, regularizers, and training hyperparameters, can be found in the Supplementary Methods, Sec. S2.

### PAGA-like directed connectivity

For statistical validation we compute a directed PAGA-like [35] abstraction graph over the meta-states. The undirected confidence matrix is the degree-corrected enrichment of inter-group kNN edges [52]; a permutation null model with 200 random label permutations is used to compute empirical *p*-values, which are then corrected for multiple testing using Benjamini-Hochberg [36]. The directed flux matrix is built from local cell-to-cell transitions weighted by alignment with the VelOT vector field and by spatial proximity, with negative alignments clipped to zero so that only velocity-consistent transitions receive positive weight. The signed antisymmetric flux defines the directionality index *D*_*ab*_ ∈ [−1, 1].

### Cell-type and meta-state contingency analysis

For each meta-state *a* and each cell-type label *c*, we computed the observed count *n*_*a,c*_ of cells assigned to that combination using the hard meta-state assignment arg max_*k*_ *q*_*ik*_. Expected counts under the null of independence where *E*_*a,c*_ = *n*_*a,•*_ *n*_*•,c*_*/N*, where *n*_*a,•*_ and *n*_*•,c*_ are the row and column marginals and *N* the total. The log_2_ observed-over-expected enrichment was used as the effect size. Two-sided Fisher exact tests [53] were applied to each cell of the contingency table; resulting *P* -values were corrected for multiple testing across all *K* ×|*C*| cells using the Benjamini-Hochberg procedure [36], and −log_10_ FDR was reported alongside the effect size.

### Benchmarking RNA velocity methods

#### Datasets

##### Synthetic datasets

Synthetic datasets were generated using dyngen to evaluate RNA velocity performance under controlled developmental topologies and downloaded from https://github.com/Spencerfar/LatentVelo/tree/main/synthetic_datasets. The generated datasets contain 5,000 simulated cells each. These datasets were designed to reproduce canonical developmental structures commonly used in RNA velocity benchmarking.

For the CBDir metric, ground-truth transitions were defined using the milestone transitions provided by dyngen.

##### Pancreatic endocrinogenesis

Mouse pancreatic endocrinogenesis data sampled at embryonic day E15.5 from [39] and downloaded usind the CellRank package [13]. The initially cycling population was removed to focus on endocrine differentiation trajectories. The following known biological transitions were used for CBDir evaluation: (Ngn3 low EP → Ngn3 high EP), (Ngn3 high EP → Fev+), (Fev+ → Delta), (Fev+ → Beta), Fev+ → Epsilon), Fev+ → Alpha).

##### Murine intestinal organoid

Murine intestinal organoid scEU-seq from [42] and downloaded from https://figshare.com/articles/dataset/Murine_intestinal_organoid_scEU-seq_data_-_Raw/23737170?file=41674299. The following transitions were used for CBDir evaluation: (Stem cells → TA cells), Stem cells → Goblet cells), (Stem cells → Tuft cells), (TA cells → Enterocytes).

##### Mouse hindbrain

Mouse hindbrain data containing differentiation trajectories of GABAergic interneurons and glial cells [54]. The following transitions were used for CBDir evaluation: (Neural stem cells → Progenitors), (Proliferating VZ progenitors → VZ progenitors), (VZ progenitors → Gliogenic progenitors), (VZ progenitors → Differentiating GABA interneurons), (Differentiating GABA interneurons → GABA interneurons).

##### Mouse erythroid

Erythroid lineage of mouse gastrulation from [40] and downloaded from the scVelo package [7]. The following transitions were used for CBDir evaluation: (Blood progenitors 1 → Blood progenitors 2), (Blood progenitors 2 → Erythroid1), (Erythroid1 → Erythroid2), (Erythroid2 → Erythroid3).

### Compared methods

We compared VelOT against scVelo (stochastic and dynamical) [7], DeepVelo [10], and FluxMatching [38] with the preprocessing indicated by the authors of each model (Scanpy [3, 55] defaults, *D* = 30 PCs, *k* = 30 neighbors and *n* = 2,000 highly variable genes).

### Dataset preprocessing

All datasets were preprocessed following a consistent general strategy while respecting the recommendations and official tutorials provided by the authors of each RNA velocity method. Most methods used a standard preprocessing workflow including total-count normalization, logarithmic transformation, highly variable gene (HVG) selection, principal component analysis (PCA), and nearest-neighbor graph construction. Unless otherwise specified by the original method, we selected approximately 2,000 highly variable genes and used embeddings ranging from 20 to 30 principal components together with neighborhood sizes of approximately 15 to 30 nearest neighbors.

Methods with dedicated preprocessing pipelines or model-specific requirements were executed according to the procedures described in their original publications or official implementations. This ensured that each method was evaluated under conditions representative of its intended use.

### Application of velocity models

For all benchmarked methods, we followed the preprocessing and execution procedures described in the corresponding publications, documentation pages, or official tutorials. Method-specific parameters were kept as those recommended default settings. All analyses were performed independently for each dataset and method using separate environments and scripts when necessary to ensure compatibility with the original software dependencies.

### Definition of cell-state transitions

When evaluating velocity directionality using the Cross-Boundary Directedness (CBDir) metric, we used known biological transitions between annotated cell states. These transitions correspond to established developmental or differentiation trajectories commonly used in the RNA velocity literature. Detailed information about the datasets, annotations, and transition definitions used in this study is provided in the Data Availability section.

### Evaluation metrics

We evaluated inferred dynamics using known cell-type transitions through the Cross-Boundary Directedness (CBDir) and In-Cluster Coherence (ICCoh) metrics, following previous work on RNA velocity benchmarking [9, 13, 56]. CBDir quantifies whether velocities at the boundary between two cell states are oriented toward the expected target population. For a transition *A*→ *B*, the metric measures the fraction of neighboring cells in population *B* toward which the velocity vector of a boundary cell is directed. To improve robustness to noisy cluster boundaries, we considered agreement with any neighboring target cell rather than requiring alignment toward a specific cell. Values above 0.5 therefore indicate a preferential transition in the expected direction.

The CBDir score for a cell *i* is defined as

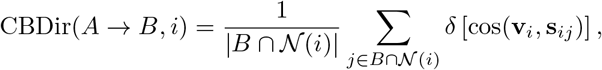

where **v**_*i*_ is the inferred velocity vector of cell *i*, **s**_*ij*_ is the normalized displacement vector between cells *i* and *j*, 𝒩 (*i*) denotes the neighborhood of cell *i*, and *δ*[·] is an indicator function evaluating whether the cosine similarity is positive.

We additionally used the ICCoh metric to evaluate local consistency of inferred velocities within the same cell population. ICCoh measures the average cosine similarity between the velocity vector of a cell and those of neighboring cells belonging to the same cluster:

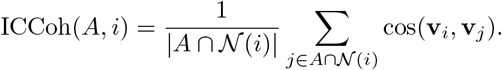

Both metrics were computed in the embedding space used for RNA velocity inference, enabling consistent evaluation across methods and datasets. Implementation details for both metrics were adapted from the LatentVelo [57] framework.

### Software and reproducibility

VelOT is implemented in Python using `PyTorch` [58], `NumPy` [59], `SciPy` [60], `scikit-learn` [61], `Scanpy` [3, 55], `anndata, POT` [50], and `NetworkX` [62]. Source code, tutorial notebooks for all benchmark datasets, and pre-trained MetaFlow models are released as an open-source Python package (velot; see *Code availability* ). All hyperparameters used in this paper are stored as YAML configuration files in the package; random seeds are fixed for reproducibility.

## 5 Declarations

### Data availability

All datasets used in this work are publicly available. The pancreas endocrinogenesis dataset [39] can be accessed via cellrank.datasets.pancreas(); the mouse gastrulation atlas erythroid subset [40] via scvelo.datasets.erythroid(); the murine intestinal organoid scRNA/scEU-seq differentiation dataset from Battich *et al*. [42], analyzed in UniTVelo Fig. 4 [8], is available through GEO accession GSE128365 and the UniTVelo intestinal organoid tutorial; the developing mouse brain hindbrain subset [41] via the original publication’s GEO record. The oligodendroglioma scRNA-seq dataset [44] is available from GEO accession GSE70630. Preprocessed AnnData files, benchmark output tables, pre-trained MetaFlow model weights, and HTML exports of all reproducibility notebooks are deposited at Zenodo (https://doi.org/10.5281/zenodo.20530116).

### Code availability

VelOT is implemented in Python. The working URL currently indicated is https://github.com/lucas-rdlr/velot (archived at https://doi.org/10.5281/zenodo.20542878). The repository contains: the core velot library; the VAMPFlow MetaFlow notebooks; reproducibility notebooks for every results figure in this paper; and Read-the-Docs documentation available at https://velot.readthedocs.io. The reproducibility package accompanying this manuscript also contains the four supplementary notebooks used for the hardware, ablation, cell-scaling, and robustness analyses, together with the parsed tables and figure files used below.

### Authors’ contributions

L.R.R. implemented the VelOT pipeline, the VAMP MetaFlow module, the benchmarking workflow, all figures, and the integration of the velot package in GitHub and Read the Docs. A.A. conceived the project, supervised the work, and helped develop the pipeline. D.P.G. provided mathematical guidance on optimal transport, flow matching, and VAMP/Koopman theory, and supervised the methodological design. All authors discussed the results and wrote the manuscript in collaboration.

## Acknowledgements

We thank the BRIGHT team at the Paris Brain Institute (ICM) and the BRAINTWIN consortium for fruitful discussions. We are grateful to the scverse community [55] for the open-source ecosystem on which VelOT is built.

Additional supplementary analyses derived from the four reproducibility notebooks have been incorporated in Supplementary Methods, Sec. S4. They include hardware comparisons (GPU, one CPU thread, and eight CPU workers), an ablation summary, cell-number scaling, robustness sweeps, fate-readout diagnostics, vector-field galleries, and numeric tables parsed from the uploaded notebook outputs. Full per-run tables are distributed in the reproducibility bundle because they are more appropriate as machine-readable files than as long printed tables in the manuscript.

## Abbreviations

scRNA-seq: single-cell RNA sequencing.
i.i.d.: independent and identically distributed.
OT: optimal transport.
FM: flow matching.
CFM: conditional flow matching.
CNF: continuous normalizing flows.
DPT: diffusion pseudotime.
HVG: highly variable gene.
PCA: principal component analysis.
kNN: *k*-nearest neighbor.
UMAP: Uniform Manifold Approximation and Projection.
MLP: multi-layer perceptron.
VAMP: Variational Approach for Markov Processes.
MSM: Markov state model.
PAGA: partition-based graph abstraction.
CBDir: cross-boundary directionality.
ICCoh: intra-cluster coherence.
BH: Benjamini-Hochberg.
FDR: false discovery rate.

