## Supplementary material and figures for "VelOT: kinetic-free RNA velocity inference via optimal transport, flow-field smoothing, and VAMP coarse-graining of cellular dynamics"

**Contents.** This Supplementary Information provides extended mathematical background, a full specification of the VelOT-MetaFlow module, benchmarking and statistical testing details, biological supplementary results, and a dedicated synthetic validation section. The synthetic validation section reports hardware timing, cell-number scaling, ablation families, perturbation robustness, terminal routing, and per-cell confidence diagnostics. Supplementary references appear at the end of this standalone document.

**Guide to Supplementary Figures.** The figure guide below is provided at the beginning of the Supplementary Information to make the standalone file easier to inspect. It includes the original biological and benchmark figures retained from the previous supplementary version, followed by the expanded synthetic validation and ablation figures added in this revision.

**Table S1**: Overview of all supplementary figures included in this document.

| **Figure** | **Topic** | **Purpose** |
|---|---|---|
| Fig. S2 | Soft meta-state memberships | Per-cell membership maps showing local ambiguity and state overlap. |
| Fig. S3 | Meta-state versus cell-type contingency | Count, fraction, enrichment summaries and flow representation linking meta-states to cell types. |
| Fig. S4 | Bidirectional composition bars | Complementary meta-state-by-cell-type and cell-type-by-meta-state summaries. |
| Fig. S7 | Erythroid full analysis | Gene programs, vector-field flow, and feature associations along erythroid trajectories. |
| Fig. S5 | Pancreas path gene heatmaps | Gene expression along source-to-terminal MetaFlow paths. |
| Fig. S6 | Meta-state gene heatmap | Compact marker-gene signatures for each pancreas meta-state. |
| Fig. S8 | Benchmark significance | Per-cell CBDir, ICCoh, and runtime distributions with statistical tests. |
| Fig. S9 | Oligodendroglioma streamlines | Additional neuro-oncology application showing AC-like and OC-like differentiation axes. |
| Fig. S10 | Hardware timing | GPU, CPU-1, and CPU-8 runtime comparison on the synthetic bifurcation. |
| Fig. S11 | Hardware field equivalence | Quiver and streamline fields confirming that hardware changes do not alter the inferred dynamics. |
| Fig. S12 | Cell-number scaling | Runtime, memory, OT pair count, amortized cost, and quality versus cell number. |
| Fig. S13 | Stage-wise log-log scaling | Per-stage scaling of window construction, OT, smoothing, projection, and metrics. |
| Fig. S14 | Scaling exponents and speed-up | Fitted empirical exponents and GPU speed-up factors across cell numbers. |
| Fig. S15 | Speed-quality Pareto ablation | Joint comparison of runtime and composite quality across ablation families. |
| Fig. S16 | Per-family ablation metrics | Mean and seed-level variability of CBDir, ICCoh, and composite quality by ablation family. |
| Fig. S17 | Paired ablation tests | Exact paired sign-flip tests against baseline for quality and runtime metrics. |
| Fig. S18 | Ablation metric heatmap | Normalized multi-metric summary of quality, confidence, low-confidence fraction, and speed. |
| Fig. S19 | Ablation vector-field gallery | Representative streamline fields for baseline and selected ablations. |
| Fig. S20 | Smoothing convergence | Training curves for total, regression, smoothness, curl, and divergence losses. |
| Fig. S21 | Perturbation robustness | Quality, terminal reach, and fate entropy under geometry noise, imbalance, and pseudotime corruption. |
| Fig. S22 | Terminal reach matrices | Terminal-routing probabilities across perturbation sweeps. |
| Fig. S23 | Monotonic trend tests | Spearman trend tests for quality, reach, confidence, and fate entropy. |
| Fig. S24 | Forward trajectories | Representative fields with integrated trajectories from root cells under perturbations. |
| Fig. S25 | Routing alluvial diagrams | Root-to-terminal route proportions for representative perturbation regimes. |
| Fig. S26 | Confidence diagnostics | Mean confidence and zero-confidence fractions across perturbation sweeps. |

### S1 Extended background: optimal transport and flow matching for single-cell biology

#### S1.1 Optimal transport

Optimal transport (OT) is a mathematical theory dating back to Monge (1781) and Kantorovich [1] that formalizes the cheapest way to morph one probability distribution into another [2, 3]. Let $\mu$ and $\nu$ be two probability measures on a metric space $(\mathcal{X}, c)$, where $c : \mathcal{X} \times \mathcal{X} \to \mathbb{R}_{\geq 0}$ is a cost function. The Kantorovich formulation seeks a coupling $\gamma \in \Pi(\mu, \nu)$ (a joint distribution with marginals $\mu$ and $\nu$) that minimizes

$$\mathcal{W}_c(\mu, \nu) = \min_{\gamma \in \Pi(\mu,\nu)} \int_{\mathcal{X}\times\mathcal{X}} c(x, y)\, d\gamma(x, y). \tag{S1}$$

When $c(x, y) = \|x - y\|_2^p$, $\mathcal{W}_p(\mu, \nu) = \mathcal{W}_c^{1/p}$ is the $p$-Wasserstein distance.

In the discrete case, $\mu = \sum_i a_i \delta_{x_i}$ and $\nu = \sum_j b_j \delta_{y_j}$, and $\gamma$ is a non-negative matrix $\mathbf{P} \in \mathbb{R}_{\geq 0}^{N_\mu \times N_\nu}$ with row sums $\mathbf{a}$ and column sums $\mathbf{b}$. Computing $\mathbf{P}$ exactly requires solving a linear program of complexity $O(N^3 \log N)$ for $N = \max(N_\mu, N_\nu)$, which is prohibitive at single-cell scale (typically $N \sim 10^5$ to $10^6$).

#### S1.2 Entropy-regularized OT and the Sinkhorn algorithm

Cuturi [4] proposed regularizing the Kantorovich problem with the entropy of the coupling:

$$\mathbf{P}_\varepsilon^* = \operatorname*{arg\,min}_{\mathbf{P}\in\Pi(\mathbf{a},\mathbf{b})} \langle \mathbf{P}, \mathbf{C}\rangle_F - \varepsilon H(\mathbf{P}), \qquad H(\mathbf{P}) = -\sum_{ij} P_{ij} \log P_{ij}, \tag{S2}$$

where $\varepsilon > 0$ controls the smoothness of the optimal coupling. The Lagrangian dual decomposes into matrix-vector operations and the optimum admits the form $P_{ij}^* = u_i K_{ij} v_j$ with $K_{ij} = \exp(-C_{ij}/\varepsilon)$. The Sinkhorn algorithm [5] alternates row and column normalizations to obtain $\mathbf{u}$ and $\mathbf{v}$ in $O(N^2 I)$ for $I \sim 50$ to 200 iterations, making OT tractable on modern GPU hardware. Theoretical convergence properties and finite-sample guarantees are reviewed in [6, 7]. Connections between entropic OT and statistical estimation, including the Sinkhorn divergence (which is symmetric, positive, and convex), are developed in [7–9]. Variants relevant to single-cell biology include unbalanced OT [10], the Wasserstein barycenter [8, 11], the Gromov-Wasserstein distance for non-aligned domains [12], and convolutional Wasserstein distances on geometric grids [13].

#### S1.3 Static and dynamic OT for single-cell trajectories

Schiebinger *et al.* [14] introduced the Waddington-OT framework, which solves static OT between consecutive experimental timepoints to reconstruct ancestor-descendant relationships during cellular reprogramming. The static formulation, while powerful, models trajectories as straight-line interpolations and cannot capture intrinsic curvature of the manifold. Dynamic OT [15–17] replaces straight lines with continuous paths $\gamma_t$ that minimize an action functional, and has been operationalized through neural ODEs and continuous normalizing flows. The TrajectoryNet model [15] learns such continuous paths in scRNA-seq data; MIOFlow [18] couples manifold-aware regularization; moscot [17] provides a unified framework with multimodal extensions; and [19, 20] extend OT to perturbation prediction and learned cost functions.

VelOT differs from all these methods in three respects: (i) it solves OT between *intra-dataset* windows defined along a pseudotemporal axis, not between experimentally distinct timepoints; (ii) it uses OT to estimate *instantaneous velocity*, not to interpolate distributions; and (iii) the smoothing step is decoupled from the OT step, so that the discrete OT plans can be inspected and validated independently of the smoothing.

#### S1.4 Flow matching and conditional flow matching

Flow matching (FM) [21] is a simulation-free training objective for continuous normalizing flows [22]. Given a probability path $\{p_t\}_{t\in[0,1]}$ that interpolates between a source $p_0$ and a target $p_1$, and a vector field $u_t$ that generates this path (i.e. $\partial_t p_t + \nabla \cdot (p_t u_t) = 0$), FM trains a neural vector field $v_\theta(x, t)$ to minimize

$$\mathcal{L}_{\mathrm{FM}}(\theta) = \mathbb{E}_{t, x\sim p_t} \left\| v_\theta(x, t) - u_t(x) \right\|^2. \tag{S3}$$

Crucially, $u_t$ is generally intractable, but the conditional flow matching (CFM) trick [21, 23, 24] replaces $u_t$ by a conditional vector field $u_t(x|z)$ for a latent $z$ (for example, the endpoints of a coupling), and the loss reduces to a simple regression. When the coupling is the optimal-transport plan, the resulting OT-CFM [23] approximates dynamic OT.

The smoothing step of VelOT is conceptually a CFM step where the source and target pairs are sampled from the OT plans of Sec. S1.1: the network regresses on local OT displacements at $\lambda = 0$ (source $x_0$) with an added smoothness regularizer. Unlike full CFM, VelOT does not require the network to be invertible nor to define a normalizing flow; it acts purely as a smoother of the OT field. This choice trades a small modeling expressivity loss for substantial gains in training stability and computational efficiency.

##### S1.5 Other transport-based and flow-based tools in single-cell biology

The single-cell community has accumulated a rich toolbox built on related ideas, all of which are complementary to VelOT. Waddington-OT [14] reconstructs ancestor-descendant relationships across experimental timepoints during reprogramming. Yang *et al.* [25] use autoencoders combined with OT to predict cell lineages. SCOT [26] aligns multi-omic measurements through Gromov-Wasserstein distances. CellOT and unbalanced parameterized Monge maps [10, 20] extend OT to perturbation responses. PHATE and MAGIC [27] use diffusion operators to denoise gene expression. Generative single-cell models such as scVI [28], DCA [29], and the recent foundation models scGPT [30], Geneformer [31], and scELMo [32] provide alternative latent representations that could in principle replace PCA in VelOT.

### S2 Mathematical specification of the VelOT-MetaFlow module

The VelOT-MetaFlow module takes the smoothed VelOT velocity field and produces a coarse-grained, statistically calibrated landscape of cell-state transitions. It combines (i) preprocessing of latent representations, pseudotime, and velocity lifting; (ii) OT-flow matching with the VelOT field as soft anchor; (iii) the VAMPFlow meta-state estimator; and (iv) a PAGA-like directed connectivity analysis. We provide here the complete mathematical specification; the reader is referred to [33–37] for further background.

#### S2.1 Preprocessing for MetaFlow

Let $N$ be the number of cells and $G$ the number of genes; let $\mathbf{Z}_{\text{raw}} \in \mathbb{R}^{N\times d}$ be a latent representation (typically the top $d$ PCs of the log-normalized count matrix). We standardize each dimension,

$$\mathbf{Z} = \frac{\mathbf{Z}_{\text{raw}} - \hat{\mu}}{\hat{\sigma}}, \qquad \hat{\mu}_j = \frac{1}{N}\sum_i Z_{ij}^{\text{raw}}, \qquad \hat{\sigma}_j = \sqrt{\frac{1}{N}\sum_i (Z_{ij}^{\text{raw}} - \hat{\mu}_j)^2 + \epsilon}, \tag{S4}$$

with $\epsilon = 10^{-8}$ for numerical stability.

A monotone pseudotime $\tau : \{1,\dots,N\} \to [0,1]$ orders cells along the developmental trajectory; we reuse the value computed in the main VelOT pipeline when present, otherwise we compute DPT [38]. The interval $[0,1]$ is partitioned into $B = 8$ quantile-equal bins, giving a bin assignment $b_i \in \{0,\dots,B-1\}$. These bins serve as marginals for the downstream OT couplings.

***Lifting the velocity to latent space.***

The VelOT vector field $\mathbf{V}^{\text{emb}} \in \mathbb{R}^{N\times 2}$ is defined in 2D UMAP space. To lift it to the latent space we use a per-cell ridge regression. For each cell $i$, let $\mathcal{N}(i) = \{i_1,\dots,i_k\}$ be its $k$-nearest neighbors in embedding space, and define

$$\mathbf{X}^{(i)} = \left[\mathbf{E}_{i_1} - \mathbf{E}_i, \dots, \mathbf{E}_{i_k} - \mathbf{E}_i\right]^\top \in \mathbb{R}^{k\times 2}, \quad \mathbf{Y}^{(i)} = \left[\mathbf{Z}_{i_1} - \mathbf{Z}_i, \dots, \mathbf{Z}_{i_k} - \mathbf{Z}_i\right]^\top \in \mathbb{R}^{k\times d}. \tag{S5}$$

The ridge solution is

$$\hat{\mathbf{B}}^{(i)} = \left(\mathbf{X}^{(i)\top}\mathbf{X}^{(i)} + \lambda \mathbf{I}_2\right)^{-1}\mathbf{X}^{(i)\top}\mathbf{Y}^{(i)}, \qquad \lambda = 10^{-3}, \tag{S6}$$

and the lifted velocity is $\mathbf{V}^{\text{VelOT}}_{\text{latent},i} = \mathbf{V}^{\text{emb}}_i \hat{\mathbf{B}}^{(i)} \in \mathbb{R}^d$. If a direct latent-space velocity is already stored in `adata.obsm`, it is used verbatim.

#### S2.2 OT couplings between adjacent pseudotime bins

For adjacent bins $(t, t+1)$, let $\mathcal{S}_t = \{i : b_i = t\}$ and $\mathcal{S}_{t+1} = \{i : b_i = t+1\}$, capped at $M = 800$ cells per side. The Sinkhorn-regularized plan is

$$\gamma^{(t,t+1)} = \underset{\gamma\in\Pi(\mu_t,\mu_{t+1})}{\arg\min} \sum_{i,j} \gamma_{ij} \left\|\mathbf{Z}_i - \mathbf{Z}_j\right\|^2 + \varepsilon \sum_{i,j} \gamma_{ij} \log \gamma_{ij}, \tag{S7}$$

with $\varepsilon = 0.05$ and 80 Sinkhorn iterations.

#### S2.3 OT-flow-matching neural vector field

A time-conditioned neural vector field $f_\theta : \mathbb{R}^d \times [0,1] \to \mathbb{R}^d$ is trained to minimize the CFM objective [21, 23, 24] augmented with a VelOT alignment term. At each training step we sample a bin pair $(t, t+1)$, source and target indices $(i, j)$ with probability proportional to $\gamma_{ij}^{(t,t+1)}$, and an interpolation factor $\lambda \sim U(0,1)$. The interpolant is

$$\tau_\lambda = (1-\lambda)\tau_i + \lambda\tau_j, \qquad z_\lambda = (1-\lambda)x_0 + \lambda x_1 + \sigma\xi, \quad \xi \sim \mathcal{N}(0, \mathbf{I}), \tag{S8}$$

with $\sigma = 0.01$. The flow-matching loss is

$$\mathcal{L}_{\text{FM}}(\theta) = \mathbb{E}\left[\left\| f_\theta(z_\lambda, \tau_\lambda) - \tfrac{x_1 - x_0}{\tau_j - \tau_i} \right\|^2\right]. \tag{S9}$$

The VelOT alignment loss, with weight $\omega_{\text{align}} = 0.15$, ensures consistency with the input field:

$$\mathcal{L}_{\text{align}}(\theta) = \tfrac{1}{2}\left(1 - \tfrac{f_\theta(x_0,\tau_i)\cdot \mathbf{V}^{\text{VelOT}}_{\text{latent},i}}{\|f_\theta(x_0,\tau_i)\| \left\|\mathbf{V}^{\text{VelOT}}_{\text{latent},i}\right\|}\right) + \tfrac{1}{2}\text{MSE}_{\text{scale}}\left(f_\theta(x_0,\tau_i), \mathbf{V}^{\text{VelOT}}_{\text{latent},i}\right), \tag{S10}$$

where MSE is computed after rescaling the target to the median norm of the prediction. The network is a 4-residual-block time-conditioned MLP (256 hidden units, LayerNorm, SiLU); training uses AdamW ($\eta = 10^{-3}$, weight decay $10^{-4}$) for $T_{\text{flow}} = 1{,}200$ epochs.

#### S2.4 Final velocity and forward propagation

Three velocity modes are available:

- `velot_only`: $\mathbf{V}^* = \mathbf{V}^{\text{VelOT}}_{\text{latent}}$.
- `ot_only`: $\mathbf{V}^* = \mathbf{V}^{\text{OT}}$ (the CFM-learned field).
- `blend` (default, $\alpha = 0.50$): $\mathbf{V}^* = (1-\alpha)\mathbf{V}^{\text{OT}} + \alpha\tilde{\mathbf{V}}^{\text{VelOT}}$, with $\tilde{\mathbf{V}}^{\text{VelOT}}$ rescaled to the median norm of $\mathbf{V}^{\text{OT}}$.

The future state used by all meta-state estimators is

$$\mathbf{Z}_i^{\text{future}} = \mathbf{Z}_i + \Delta t \cdot \mathbf{V}_i^*, \qquad \Delta t = 0.15. \tag{S11}$$

A diagnostic per-cell alignment is computed as

$$a_i = \frac{\mathbf{V}_i^{\text{OT}} \cdot \tilde{\mathbf{V}}_i^{\text{VelOT}}}{\left\|\mathbf{V}_i^{\text{OT}}\right\| \left\|\tilde{\mathbf{V}}_i^{\text{VelOT}}\right\|} \in [-1, 1], \tag{S12}$$

with $a_i < 0$ indicating local disagreement between OT-induced and VelOT-induced directions.

#### S2.5 Vector-field diagnostics: divergence and curl

The local divergence indicates whether mass leaves ($\nabla \cdot \mathbf{V} > 0$) or enters ($\nabla \cdot \mathbf{V} < 0$) a neighborhood. For cell $i$ with $k$-NN $\mathcal{N}(i)$, we estimate the local Jacobian $\hat{\mathbf{B}}^{(i)} \in \mathbb{R}^{d\times d}$ by least squares,

$$\hat{\mathbf{B}}^{(i)} = \arg\min_{\mathbf{B}} \sum_{j\in\mathcal{N}(i)} \left\|(\mathbf{Z}_j - \mathbf{Z}_i)\mathbf{B} - (\mathbf{V}_j - \mathbf{V}_i)\right\|^2, \tag{S13}$$

and the divergence is $\hat{\nabla} \cdot \mathbf{V}_i^* = \operatorname{tr}(\hat{\mathbf{B}}^{(i)})$, z-scored across cells.

The $z$-curl in the 2D embedding,

$$\omega_i = \frac{\partial V_y^{\text{emb}}}{\partial e_x} - \frac{\partial V_x^{\text{emb}}}{\partial e_y}, \tag{S14}$$

is estimated by the 2D analogue of the Jacobian above; high $|\omega_i|$ indicates cyclic flow (cell-cycle or recurrent differentiation).

#### S2.6 VAMPFlow estimator

A stochastic process $\{z_t\}$ over cell states is described by the transfer (Koopman) operator $\mathcal{K}_\tau$ [39–45], which acts on a square-integrable observable $f$ by

$$[\mathcal{K}_\tau f](z) = \mathbb{E}[f(z_{t+\tau}) \mid z_t = z]. \tag{S15}$$

The Variational Approach for Markov Processes (VAMP) [33, 34] states that the singular value decomposition (SVD) of $\mathcal{K}_\tau$ in a weighted $L^2$ space provides the optimal low-rank approximation of the dynamics. Given a basis of $K$ observable functions $\boldsymbol{\chi} = (\chi_1, \ldots, \chi_K)$, the time-lagged covariance matrices are

$$\mathbf{C}_{00} = \mathbb{E}[\boldsymbol{\chi}(z_t)^\mathsf{T}\boldsymbol{\chi}(z_t)], \quad \mathbf{C}_{11} = \mathbb{E}[\boldsymbol{\chi}(z_{t+\tau})^\mathsf{T}\boldsymbol{\chi}(z_{t+\tau})], \quad \mathbf{C}_{01} = \mathbb{E}[\boldsymbol{\chi}(z_t)^\mathsf{T}\boldsymbol{\chi}(z_{t+\tau})]. \tag{S16}$$

The VAMP-$r$ score is

$$\mathcal{V}_r(\boldsymbol{\chi}) = \left\|\mathbf{C}_{00}^{-1/2}\mathbf{C}_{01}\mathbf{C}_{11}^{-1/2}\right\|_r^r, \tag{S17}$$

where $\|\cdot\|_r^r = \sum_i \sigma_i^r$ is the sum of $r$-th powers of the singular values. For $r = 2$, the VAMP-2 score equals $\sum_{k=1}^K \sigma_k^2$ and is equivalent to the kinetic variance captured by the basis. Maximizing $\mathcal{V}_2$ over $\boldsymbol{\chi}$ recovers the leading singular functions of $\mathcal{K}_\tau$, which encode the slowest relaxation modes and the metastable macro-states of the process.

***Finite-sample VAMP-2 score.***

In the pipeline, cells are represented at time $t$ by $\mathbf{q}_0 = \boldsymbol{\chi}(\mathbf{Z}) \in \mathbb{R}^{N\times K}$ (current-state soft memberships) and at time $t + \Delta t$ by $\mathbf{q}_1 = \boldsymbol{\chi}(\mathbf{Z}^{\text{future}}) \in \mathbb{R}^{N\times K}$. The empirical cross-covariance matrices are (after mean-centering)

$$\hat{\mathbf{C}}_{00} = \frac{\tilde{\mathbf{q}}_0^\mathsf{T}\tilde{\mathbf{q}}_0}{N-1} + \epsilon\mathbf{I}, \quad \hat{\mathbf{C}}_{11} = \frac{\tilde{\mathbf{q}}_1^\mathsf{T}\tilde{\mathbf{q}}_1}{N-1} + \epsilon\mathbf{I}, \quad \hat{\mathbf{C}}_{01} = \frac{\tilde{\mathbf{q}}_0^\mathsf{T}\tilde{\mathbf{q}}_1}{N-1}, \tag{S18}$$

with $\epsilon = 10^{-5}$. The finite-sample VAMP-2 score is

$$\hat{\mathcal{V}}_2 = \left\| \hat{\mathbf{C}}_{00}^{-1/2}\hat{\mathbf{C}}_{01}\hat{\mathbf{C}}_{11}^{-1/2} \right\|_F^2, \tag{S19}$$

where the matrix inverse square root is computed via eigendecomposition.

***VAMPMetaStateNet architecture.***

The observable functions $\boldsymbol{\chi}_\theta : \mathbb{R}^d \to \Delta_{K-1}$ are parameterized by a deep network (VAMPnets [34]):

$$\boldsymbol{\chi}_\theta(z) = \text{softmax}(f_\theta^{(L)}(z)), \tag{S20}$$

where $f_\theta^{(L)}$ is an $L$-layer MLP with 256 hidden units, LayerNorm, SiLU activations, and dropout $p = 0.05$. The output layer has $K$ units, followed by a row-wise softmax ensuring $\sum_k q_{ik} = 1$ and $q_{ik} \geq 0$.

***VAMPFlow training objective.***

The training loss combines the negative VAMP-2 score with three regularizers:

$$\mathcal{L}_{\text{VAMP}}(\theta) = -\hat{\mathcal{V}}_2(\theta) + \lambda_{\text{bal}}\mathcal{L}_{\text{balance}} + \lambda_{\text{sharp}}\mathcal{L}_{\text{sharp}} + \lambda_{\text{orth}}\mathcal{L}_{\text{orth}}, \tag{S21}$$

with $\lambda_{\text{bal}} = 0.20$, $\lambda_{\text{sharp}} = 0.02$, $\lambda_{\text{orth}} = 0.02$, and

$$\mathcal{L}_{\text{balance}} = \left\|\bar{q} - \tfrac{1}{K}\mathbf{1}\right\|^2, \quad \bar{q}_k = \tfrac{1}{N}\sum_i q_{ik}, \tag{S22}$$

$$\mathcal{L}_{\text{sharp}} = -\tfrac{1}{N}\sum_{i=1}^{N}\sum_{k=1}^{K} q_{ik}\log(q_{ik} + \epsilon), \tag{S23}$$

$$\mathcal{L}_{\text{orth}} = \left\|\hat{\mathbf{Q}}^\mathsf{T}\hat{\mathbf{Q}} - \mathbf{I}_K\right\|_F^2, \qquad \hat{\mathbf{Q}} = \mathbf{q}\,\text{diag}(\|q_k\|^{-1})_{k=1}^K. \tag{S24}$$

The balance term prevents degenerate solutions where most cells are assigned to a single state; the sharpness (negative entropy) term encourages confident assignments; the orthogonality term reduces redundancy across states. Training uses AdamW [46] ($\eta = 10^{-3}$, weight decay $10^{-4}$, mini-batch size 2048, gradient clip at 5) for $T_{\text{VAMP}} = 1{,}200$ epochs.

***Automatic selection of $K$.***

When the user does not specify a fixed number of meta-states, the pipeline trains a short warm-up ($T_{\text{auto}} = 350$ epochs) for each candidate $K \in \{5, 6, 7, 8, 9, 10, 12\}$ and selects the value of $K$ that maximizes the compound criterion

$$\mathcal{C}(K) = \hat{\mathcal{V}}_2(K) - 0.08K + 2\min_k(\bar{q}_k) + 0.50\frac{H(\bar{q})}{\log K}, \tag{S25}$$

where $H(\bar{q}) = -\sum_k \bar{q}_k \log \bar{q}_k$. The three correction terms enforce low model complexity, non-empty states, and balanced occupancy, respectively. Default $K$ for the pancreas analysis is 12 (the automatically chosen value).

***Soft transition matrix.***

The coarse-grained transition matrix $\mathbf{T}^{\text{VAMP}} \in \mathbb{R}^{K\times K}$ is

$$\tilde{T}_{ab}^{\text{VAMP}} = \frac{\sum_i q_{ia}q_{ib}^{\text{future}}}{\sum_i q_{ia}}, \qquad T_{ab}^{\text{VAMP}} = \frac{\tilde{T}_{ab}^{\text{VAMP}}}{\sum_{b'}\tilde{T}_{ab'}^{\text{VAMP}}}, \tag{S26}$$

where $q_{ib}^{\text{future}} = q_b(\mathbf{Z}_i^{\text{future}})$.

#### S2.7 Meta-state classification and committors

For each estimator the coarse transition matrix $\mathbf{T} \in \mathbb{R}^{K\times K}$ and the soft membership $\mathbf{q} \in \mathbb{R}^{N\times K}$ yield per-state diagnostic scores. All cell-level quantities (divergence $d_i$, stemness $s_i$, curl $\omega_i$, velocity alignment $a_i$) are aggregated to state level by a soft weighted mean,

$$\bar{x}_k = \frac{\sum_i q_{ik} x_i}{\sum_i q_{ik} + \epsilon}. \tag{S27}$$

The state-level quantities are

$$s_k = T_{kk} \quad \text{(self-transition)}, \tag{S28}$$

$$o_k = \sum_{b\neq k} T_{kb} \quad \text{(outgoing flux)}, \tag{S29}$$

$$\iota_k = \sum_{a\neq k} T_{ak} \quad \text{(incoming flux)}, \tag{S30}$$

$$H_k = -\frac{\sum_b T_{kb} \log T_{kb}}{\log K} \quad \text{(row entropy)}. \tag{S31}$$

Four composite scores are computed:

$$\Sigma_k^{\text{src}} = o_k - \iota_k + 0.35\bar{d}_k + 0.35\bar{s}_k, \tag{S32}$$

$$\Sigma_k^{\text{sink}} = s_k + \iota_k - o_k - 0.35\bar{d}_k, \tag{S33}$$

$$\Sigma_k^{\text{branch}} = H_k + o_k - s_k, \tag{S34}$$

$$\Sigma_k^{\text{rec}} = s_k + 0.30\bar{\omega}_k, \tag{S35}$$

and each state is classified by the type whose score most exceeds the 75th percentile across states. States not exceeding any threshold are labeled *intermediate/transient*.

***Committor probabilities.***
Given a set of terminal states $\mathcal{T} \subset \{0, \ldots, K-1\}$, the committor probability $q^+_{ab}$ that a trajectory starting in state $a$ reaches terminal $b$ before any other terminal satisfies the linear system

$$\mathbf{B}_{\mathcal{S}} = (\mathbf{I} - \mathbf{Q})^{-1}\mathbf{R}, \tag{S36}$$

where $\mathbf{Q} = \mathbf{T}[\mathcal{S}, \mathcal{S}]$ is restricted to transient states $\mathcal{S} = \{0, \ldots, K-1\} \setminus \mathcal{T}$ and $\mathbf{R} = \mathbf{T}[\mathcal{S}, \mathcal{T}]$ is the absorption block. The per-cell committor to terminal $t$ is $q^+_i(t) = \sum_k q_{ik} B_{kt}$.

#### S2.8 PAGA-like directed connectivity

The PAGA algorithm [37] provides a statistically robust abstraction graph over cell groups. The pipeline implements a custom extension that (i) uses the VelOT or MetaFlow vector field to define directed transitions, (ii) applies a permutation null model to correct for degree effects, and (iii) computes signed directionality indices.

***Undirected kNN graph.***
A symmetric weighted graph $\mathbf{A} \in \mathbb{R}^{N\times N}_{\geq 0}$ is built from the latent space $\mathbf{Z}$ using $k$-NN with Gaussian kernel weights,

$$A_{ij} = \exp\!\Big(-\frac{\|\mathbf{Z}_i - \mathbf{Z}_j\|^2}{2\hat{\sigma}_i^2}\Big), \quad \hat{\sigma}_i = \text{median}_{j\in\mathcal{N}(i)} \|\mathbf{Z}_i - \mathbf{Z}_j\|, \tag{S37}$$

for $j \in \mathcal{N}(i)$ ($k_{\text{kNN}} = 30$ neighbors).

***Undirected connectivity, enrichment, and confidence.***
For a partition of cells into $K$ groups, define

$$C_{ab} = \sum_{(i,j)\in\mathcal{E}} A_{ij}\mathbb{I}[g_i = a]\mathbb{I}[g_j = b], \tag{S38}$$

with degree-corrected expected count

$$E_{ab} = \frac{\mathrm{vol}(a)\mathrm{vol}(b)}{2W + \epsilon}, \qquad \mathrm{vol}(a) = \sum_{i:g_i=a} d_i, \qquad W = \tfrac{1}{2}\sum_i d_i, \tag{S39}$$

where $d_i = \sum_j A_{ij}$. The enrichment is $\mathrm{enr}_{ab} = C_{ab}/(E_{ab} + \epsilon)$ and the PAGA-like confidence is $\mathrm{conf}_{ab} = \min(\mathrm{enr}_{ab}, 1)$.

***Permutation null and FDR control.***
To provide calibrated $p$-values, group labels are permuted $n_{\mathrm{perm}} = 200$ times (preserving group sizes), and the distribution of $C_{ab}^{(\pi)}$ under the null is estimated. The empirical $p$-value is

$$p_{ab} = \frac{1 + \#\{\pi : C_{ab}^{(\pi)} \geq C_{ab}\}}{1 + n_{\mathrm{perm}}}, \tag{S40}$$

and a $z$-score is computed from the permutation distribution. Multiple-testing correction uses Benjamini-Hochberg [47].

***Directed transitions and directionality index.***
For cell $i$ with $k_{\mathrm{dir}}$-NN, the directed edge $i \to j$ is weighted by

$$w_{ij} \propto \exp\Big(\frac{(\mathbf{Z}_j - \mathbf{Z}_i)\cdot \mathbf{V}_i^*}{\|\mathbf{Z}_j - \mathbf{Z}_i\|\,\|\mathbf{V}_i^*\| + \epsilon}\Big/T\Big) \times \exp\Big(-\frac{\|\mathbf{Z}_j - \mathbf{Z}_i\|^2}{2\hat{\sigma}^2}\Big), \tag{S41}$$

with temperature $T = 0.25$. Negative alignment values are clipped to zero so that only velocity-consistent transitions receive positive weight. After row-normalization, the group-level flux is

$$F_{ab} = \frac{1}{n_a} \sum_{i:g_i=a} \sum_{j:g_j=b} P_{ij}, \tag{S42}$$

and the directionality index is

$$D_{ab} = \frac{F_{ab} - F_{ba}}{F_{ab} + F_{ba} + \epsilon} \in [-1, 1]. \tag{S43}$$

$D_{ab} > 0$ indicates net flow from $a$ to $b$ and $D_{ba} = -D_{ab}$ by construction.

### S3 Detailed benchmark protocol and statistical tests

***Supplementary reference scope.***

To keep the main manuscript close to the requested citation density while preserving the full methodological context, the Supplementary Methods retain additional references for trajectory inference, velocity modeling, optimal-transport geometry, latent representation learning, implementation, and neuro-oncology interpretation [48–62]. These references support the extended methodological and biological discussion but are intentionally not all repeated in the main text.

#### S3.1 Datasets and preprocessing

The four real datasets used in the benchmark were preprocessed identically: counts were normalized to 10,000 per cell, log1p-transformed, and the top 2,000 highly variable genes were retained. PCA with $D = 30$ components and a kNN graph with $k = 30$ were computed using Scanpy [63, 64] defaults. Cell-type labels were taken from the published metadata of each dataset. For the synthetic data, three topologies (linear, bifurcation, trifurcation) were generated with the dyngen simulator [65] using default biological parameters and Splatter-compatible noise.

#### S3.2 Method execution

scVelo (stochastic and dynamical) was executed via the official package with default hyperparameters and `n_top_genes=2000`. DeepVelo was run from the public reference implementation with default neural-ODE architecture and 1,000 training epochs. FluxMatching was run using its publicly available implementation, with batch size 1024 and 800 training epochs. VelOT was run with default parameters: window size $W = 80$, overlap $f_{\text{overlap}} = 0.5$, Sinkhorn iterations 80, regularization $\varepsilon = 0.05$, smoothness $\lambda_{\text{smooth}} = 0.05$, and 300 epochs of MLP training. All methods were run with identical preprocessing pipelines on a Dell Precision 7550 workstation running Ubuntu 22.04.5 LTS, equipped with an Intel Xeon W-10855M CPU (6 cores, 12 threads), 93 GB RAM, and an NVIDIA Quadro RTX 5000 Mobile GPU. GPU-accelerated computations were performed using CUDA 11.8 and PyTorch. Wall-clock times include preprocessing of velocity-specific inputs (spliced and unspliced counts for scVelo and DeepVelo).

#### S3.3 Evaluation metrics

Cross-boundary directionality (CBDir) and intra-cluster coherence (ICCoh) are the two standard scores used in recent velocity benchmarks [66–68]. For a known cell-type boundary $(c_1, c_2)$ with biologically expected direction $c_1 \to c_2$, CBDir is the fraction of $c_1$ cells whose velocity vector $\mathbf{v}_i$ projects positively on the displacement to the mean of their nearest $c_2$ neighbors:

$$\text{CBDir}(c_1, c_2) = \frac{1}{|c_1^{\text{bdr}}|} \sum_{i \in c_1^{\text{bdr}}} \mathbb{I}\Big[\mathbf{v}_i \cdot (\bar{\mathbf{x}}_{\text{NN}}^{c_2}(i) - \mathbf{x}_i) > 0\Big], \tag{S44}$$

where $c_1^{\text{bdr}}$ are cells in $c_1$ with at least one $c_2$ neighbor in the global kNN graph. ICCoh is the mean cosine similarity between a cell's velocity and the mean velocity of its kNN neighbors in the same cluster,

$$\text{ICCoh} = \frac{1}{N} \sum_{i=1}^{N} \frac{\mathbf{v}_i \cdot \bar{\mathbf{v}}_i^{\text{intra}}}{\|\mathbf{v}_i\| \, \|\bar{\mathbf{v}}_i^{\text{intra}}\|}. \tag{S45}$$

Both metrics lie in $[-1, 1]$, where higher is better and 1 corresponds to a perfectly coherent and correctly oriented field. The composite *overall ranking* reported in main Fig. 5 is the unweighted mean of normalized per-dataset ranks across CBDir, ICCoh (higher is better), and log time (lower is better).

#### S3.4 Statistical significance

Pairwise statistical tests were computed on the per-cell distributions of CBDir and ICCoh (treating each cell as an independent sample) using two-sided Wilcoxon signed-rank tests with Benjamini-Hochberg correction across the five method-pair comparisons per dataset. Synthetic topologies and implementation-level controls are summarized separately in Supplementary Figs. S10-S26; these analyses use seed-level summaries and paired or trend tests when explicitly reported. Stars in Supplementary Fig. S8 denote $^{*}P < 0.05$, $^{**}P < 0.01$, $^{***}P < 0.001$, $^{****}P < 0.0001$, and *ns* not significant.

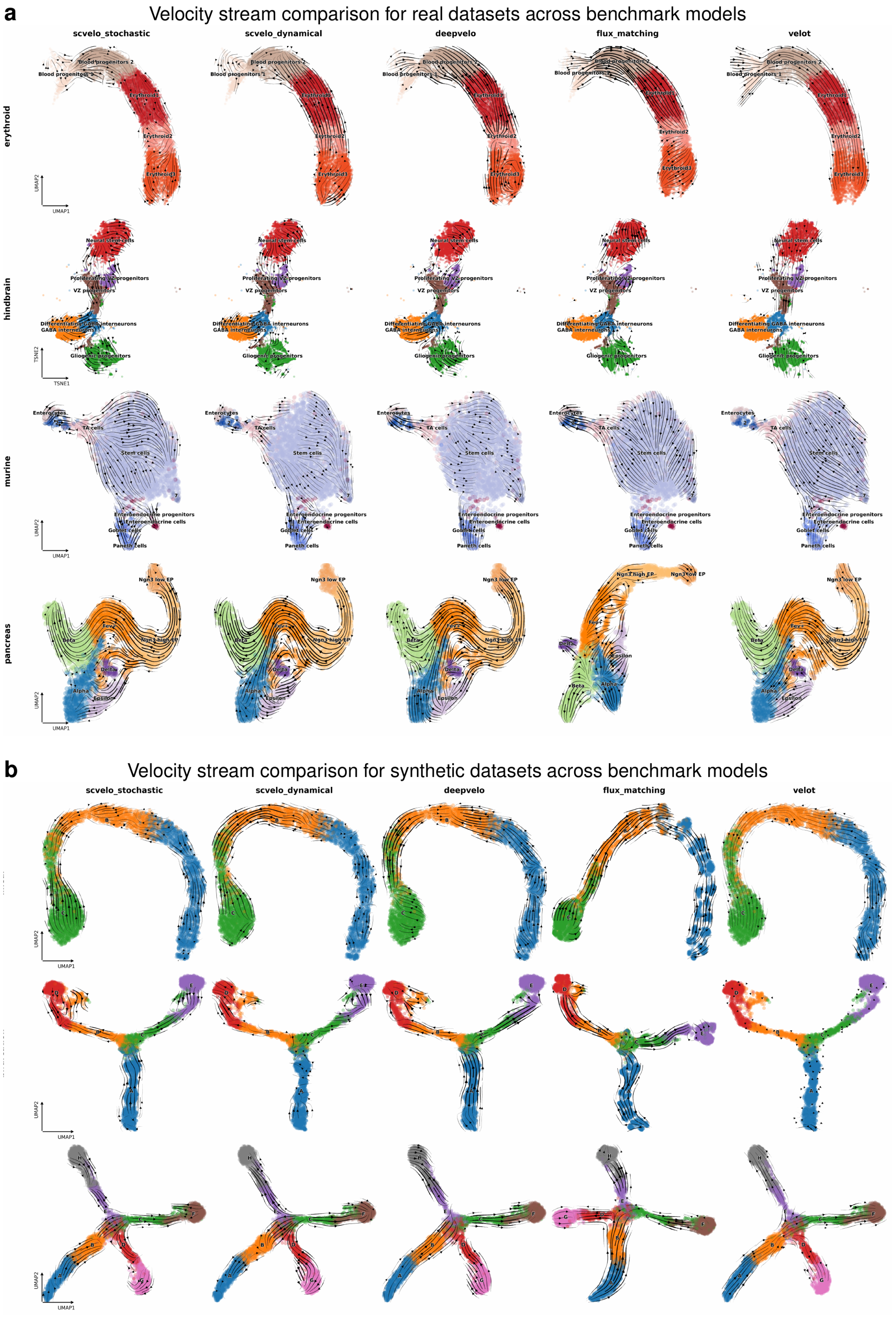

**Fig. S1 Velocity streams comparison across benchark models.** **a)** Results for the four real datasets (erythroid, hindbrain, murine, and pancreas), and the five benchmarked models. **b)** Results for the three synthetic datasets (linear, bifurcation, and trifurcation), and the five benchmarked models. By analysing the las column one can see how the VelOT method obtains much better and expected results regarding continuity and branching lineage compared to the other methods, where weird cycles and back-time flow appears; Flux Matching may re-compute UMAP coordinates, giving as a results different projections.

### S4 Supplementary results

#### S4.1 Detailed pancreas meta-state landscape

Figure S2 shows the per-cell soft membership map for each meta-state separately, providing a fine-grained view of how each meta-state is distributed across the manifold. M0, M5, and M8 are tightly localized in the progenitor pool; M7 and M10 occupy distinct positions at the first fate decision; M1, M2, M9, and M11 are the terminal $\beta$, $\varepsilon$, $\alpha$, and $\delta$ basins; M3 and M4 occupy the broader *Fev*$^+$ intermediate; M6 sits inside the cycling region of the manifold.

Figure S3 reports the full meta-state by cell-type contingency analysis as three complementary views: (a) raw contingency counts, (b) cell-type fractions within each meta-state, and (c) the $\log_2$ observed-over-expected enrichment heatmap with significance asterisks for FDR-corrected Fisher exact tests ($P < 0.05$ and $\log_2$ enrichment $> 0$). The coarse transition matrix (Fig. S3d) is row-stochastic and dominated by strong self-transitions in the terminal states (M3, M6, M7 and M8); each with self-transition $> 0.9$), with structured forward flows from M0 $\rightarrow$ M5 $\rightarrow$ M9 $\rightarrow$ M11 (transition probability $> 0.8$ for the last three) recapitulating the expected progenitor to branching to mature topology of endocrinogenesis.

A similar visualization can be seen in Figure S4a. The complementary view (Fig. S4b) inverts the contingency and shows, for each real cell type, what fraction of its cells were assigned to which meta-state. *Beta* cells split predominantly into M6 (the canonical *Beta* terminal state) and M10 (*Fev*$^+$ intermediates with some $\beta$-bias), reflecting the gradual maturation continuum from *Fev*$^+$ to *Beta*. *Fev*$^+$ cells distribute across M1 and M11 (the early branching state) and M9 (the *Delta*-leaning intermediate). *Alpha* cells concentrate in M3 (their terminal state) with a small fraction in M1 (early intermediates), and so on. This per-cell-type decomposition exposes the dynamical heterogeneity that the original cell-type labels collapse, and is a major qualitative advantage of the MetaFlow output over hard cell-type assignments.

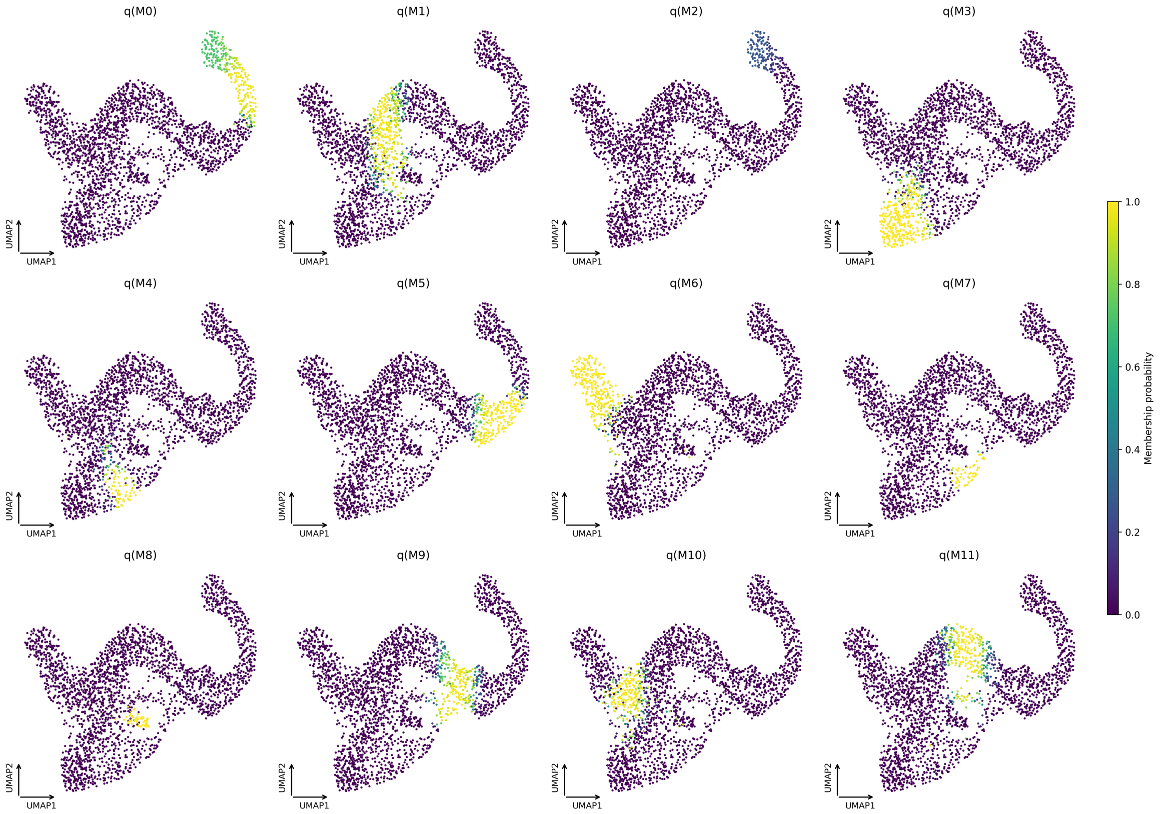

**Fig. S2 Per-cell soft memberships for each meta-state.** Each subplot shows the soft membership $q_{ik} \in [0, 1]$ of all cells to meta-state $k$ (M0 to M11) overlaid on the same UMAP coordinates. Color intensity encodes membership strength; black points correspond to cells with $q_{ik} \approx 0$. The smooth, contiguous patches of high membership confirm that meta-states are not random partitions but coherent regions of the manifold.

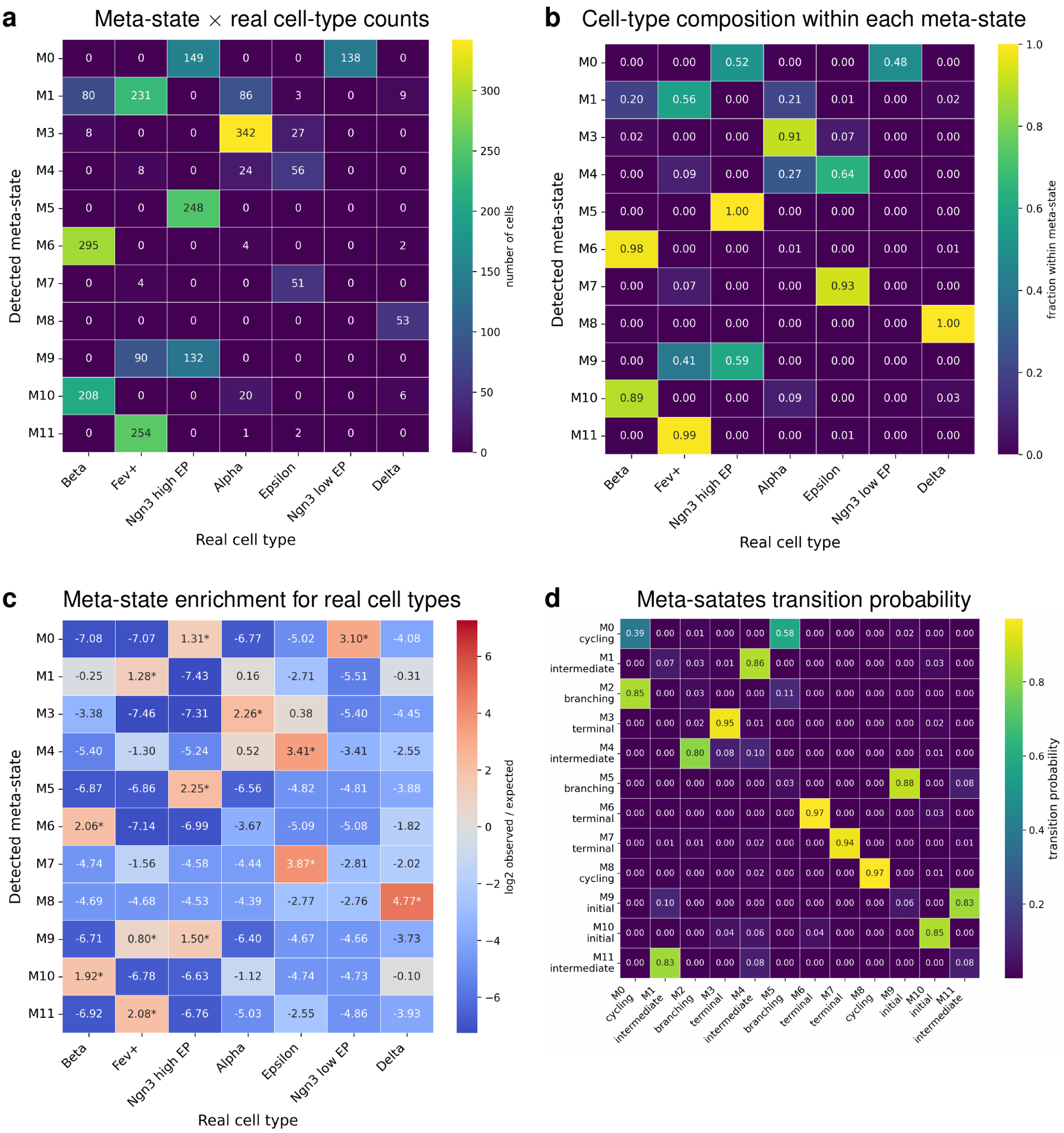

**Fig. S3 Full pancreas meta-state versus cell-type contingency analysis. a)** Raw contingency counts $n_{a,c}$ for the $12 \times 7$ table. **b)** Heatmap of cell-type fractions within each detected meta-state; each row sums to one. **c)** $\text{Log}_2$ observed-over-expected ratios from Fisher exact tests; asterisks mark cells significant at FDR $< 0.05$ with positive enrichment. The diagonal-like structure confirms that each meta-state is dominated by a single cell type while subdividing the largest types ($Fev^+$, $Ngn3^{high}$ EP, *Beta*) into multiple dynamical compartments. **d)** Rows are the current meta-state, columns the future meta-state at lag $\Delta t = 0.15$. Each row sums to one. Diagonal entries indicate self-transitions; off-diagonal entries indicate inter-state transitions. The matrix reveals a sparse forward structure consistent with developmental hierarchy: progenitor states (M0, M5, M8) have non-trivial off-diagonal entries to downstream branching states (M7, M10) which in turn flow to intermediates (M3, M4); mature terminal states (M1, M2, M9, M11) have self-transition probabilities $\geq 0.99$, indicating absorbing behavior.

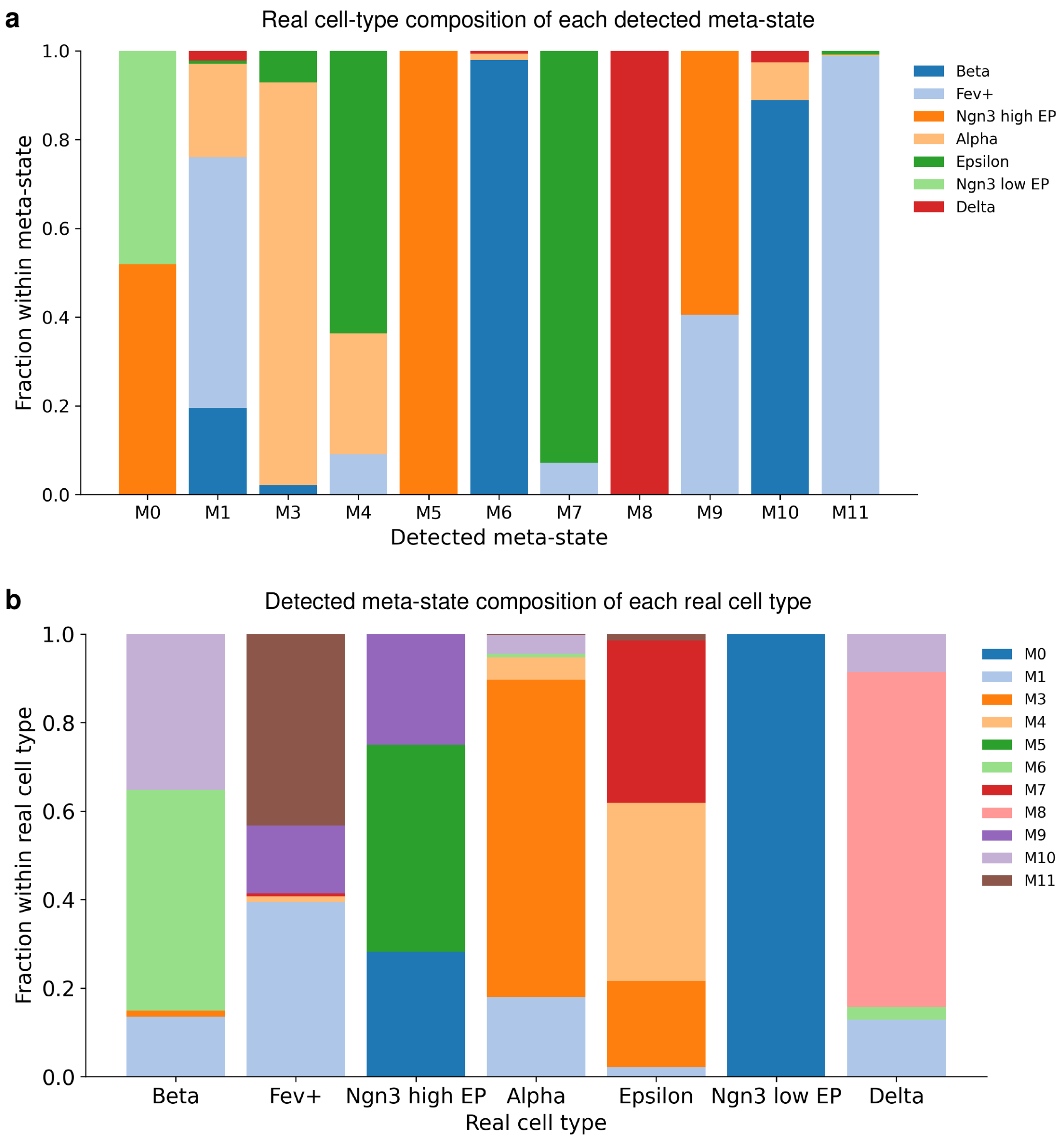

**Fig. S4 Bidirectional bar charts of the meta-state and cell-type contingency on the pancreas dataset. a)** Cell-type composition of each detected meta-state (bars are meta-states, partitioned by the seven cell types). **b)** Meta-state composition of each real cell type (bars are cell types, partitioned by the 12 meta-states). The two views are complementary: the top panel asks "*which cell types make up each meta-state?*" and the bottom asks "*which meta-states does each cell type spread across?*". Together they show that the meta-state partition is biologically interpretable (each meta-state has a clear cell-type dominant component) while also revealing sub-cell-type heterogeneity (each cell type, especially the intermediates, distributes across several meta-states).

#### S4.2 Pancreas meta-state gene programs along inferred paths

Figure S5 reports gene expression along three representative meta-state paths inferred from the pancreas VelOT-MetaFlow field: the $\beta$ path (M0→M7→M10→M3→M4→M9→M1), the $\alpha$ path (M0→M7→M10→M3→M4→M9), and the $\varepsilon$ path (M0→M7→M10→M3→M4→M9→M1→M11→M5→M8). Rows are selected genes grouped into known biological modules; columns are bins of pseudotime along the path. The expected programs are cleanly recapitulated: *Spp1*, *Sparc*, *Habp2*, *Tgm2* (early stem markers) decay at the root; *Ins1*, *Ins2*, *Nnat*, *Ppp1r1a*, *Sytl4* (mature $\beta$-cell genes) peak terminally; *Fev*, *Chgb*, *Cryba2*, *Hmgn3*, *Chga* (endocrine intermediate markers) peak in the middle. The meta-state summary heatmap (Fig. S6) compresses these per-path heatmaps into a single $K \times G$ matrix that summarizes the gene programs of each meta-state.

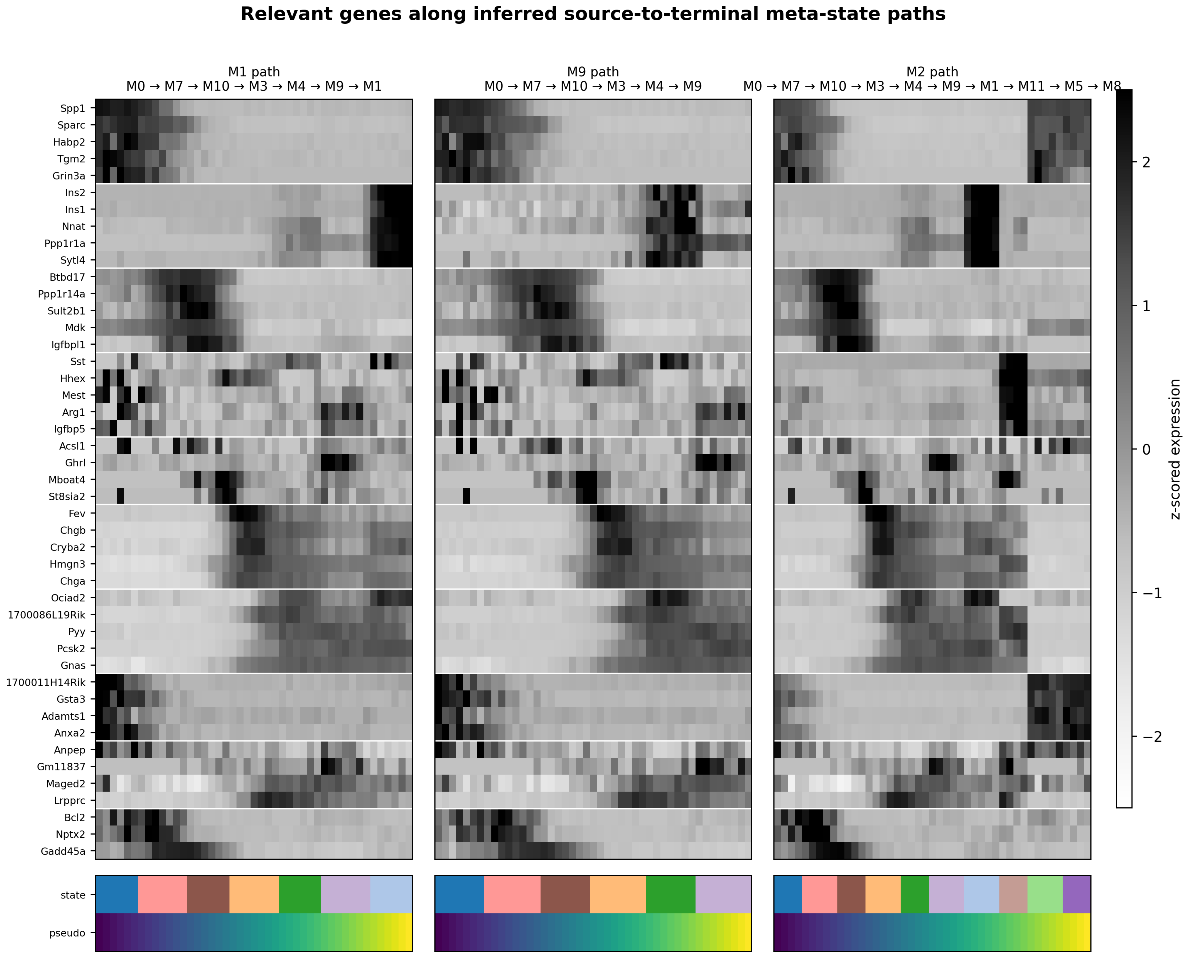

**Fig. S5 Pancreas gene expression along inferred VelOT-MetaFlow source-to-terminal paths.** Three paths from the most progenitor-like state (M0) to representative terminals (M1 $\beta$, M9 $\alpha$, and M8 $\varepsilon$) are shown side by side. Rows are genes grouped into eight biological modules (one block per module, separated by horizontal lines); columns are pseudotime bins along the path. Color encodes z-scored expression. Bottom color bars indicate the meta-state and pseudotime at each column position. The progressive activation of late markers (*Ins1, Ins2, Nnat, Pyy, Pcsk2, Gnas*) and the silencing of early markers (*Spp1, Sparc, Habp2, Tgm2*) along all paths confirms biological correctness of the MetaFlow trajectories.

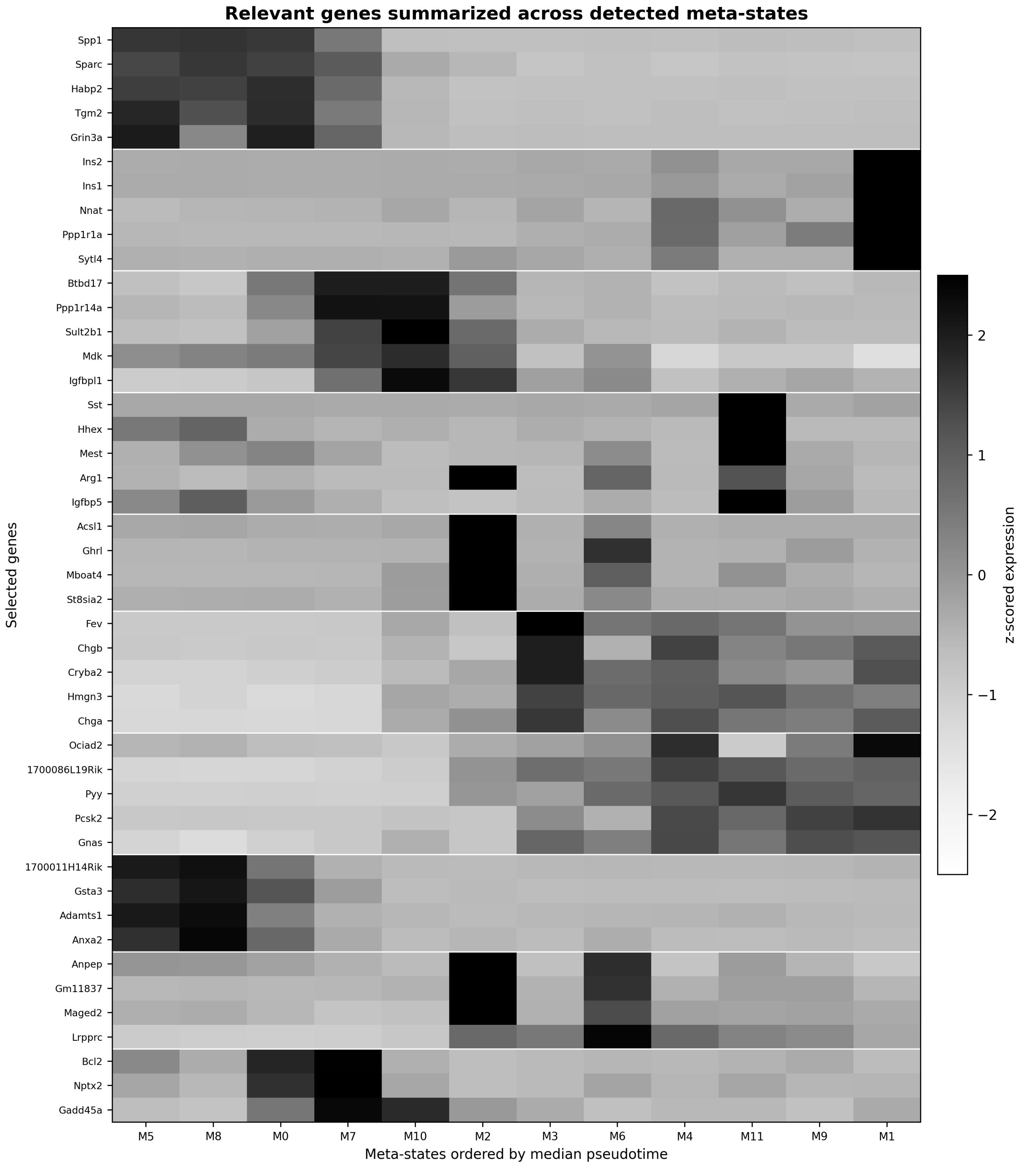

**Fig. S6 Per-meta-state gene expression summary on the pancreas dataset.** Rows are selected genes grouped into biological modules; columns are the 12 meta-states ordered by median pseudotime. Color is z-scored expression. The heatmap provides a compact, interpretable signature for each meta-state: M5 and M8 (*Ngn3*$^{low}$ EP) express early stem markers (*Spp1, Sparc, Tgm2, Grin3a*); M7 and M10 (*Ngn3*$^{high}$ EP) express committed endocrine markers (*Btbd17, Ppp1r14a, Sult2b1, Mdk, Igfbpl1*); M3 (*Fev*$^{+}$) expresses intermediate endocrine markers (*Fev, Chgb, Cryba2*); M9 (*Alpha*) shows *Arg1, Igfbp5* enrichment; M1 (*Beta*) is dominated by *Ins1, Ins2, Nnat.*

##### S4.3 Erythroid maturation: gene programs along inferred trajectories

The smoothed VelOT velocity field on the erythroid dataset (main Fig. 4) defines for each cell a downstream trajectory. Forward integration of the field from each Blood progenitor 1 cell yields paths that traverse all five clusters in the expected order. Figure S7 compiles the gene-program analysis along this trajectory. Top: a heatmap of ∼100 canonical erythroid markers (rows) along pseudotime (columns), grouped into five known programs (Stem/progenitor, Stem/progenitor 2, Erythroid priming, Membrane remodeling, Iron/heme synthesis); the trajectory recapitulates the canonical sequence *Runx1, Tal1, Lmo2* (stem) to *Meis1, Gata1* (priming) to *Car1, Tfrc, Slc4a1, Rhag, Hbb-bs* (membrane and iron/heme). Middle: single-gene expression curves for the canonical markers *Kit, Gata2, Gata1, Klf1, Tfrc, Alas2, Gypa, Hbb-bs* along pseudotime, color-coded by cell state; *Gata2* declines, *Klf1* rises, *Alas2* and *Gypa* explode at terminal cells, exactly the canonical maturation cascade. Bottom: violin plots of three VelOT-derived per-cell features (forward flow $\mathbf{v} \cdot \nabla\tau$, speed $|\mathbf{v}|$, local coherence) and Spearman correlations of marker genes with pseudotime, $\mathbf{v} \cdot \nabla\tau$, $|\mathbf{v}|$, and coherence. The strongest positive correlations of erythroid markers (*Alas2*, *Gypa*, *Hbb-bs*, $\rho \geq 0.79$) with pseudotime, and negative correlations of *Gata2* ($\rho = -0.56$) and *Kit* ($\rho = -0.13$), confirm that the VelOT-inferred trajectory tracks known erythroid biology.

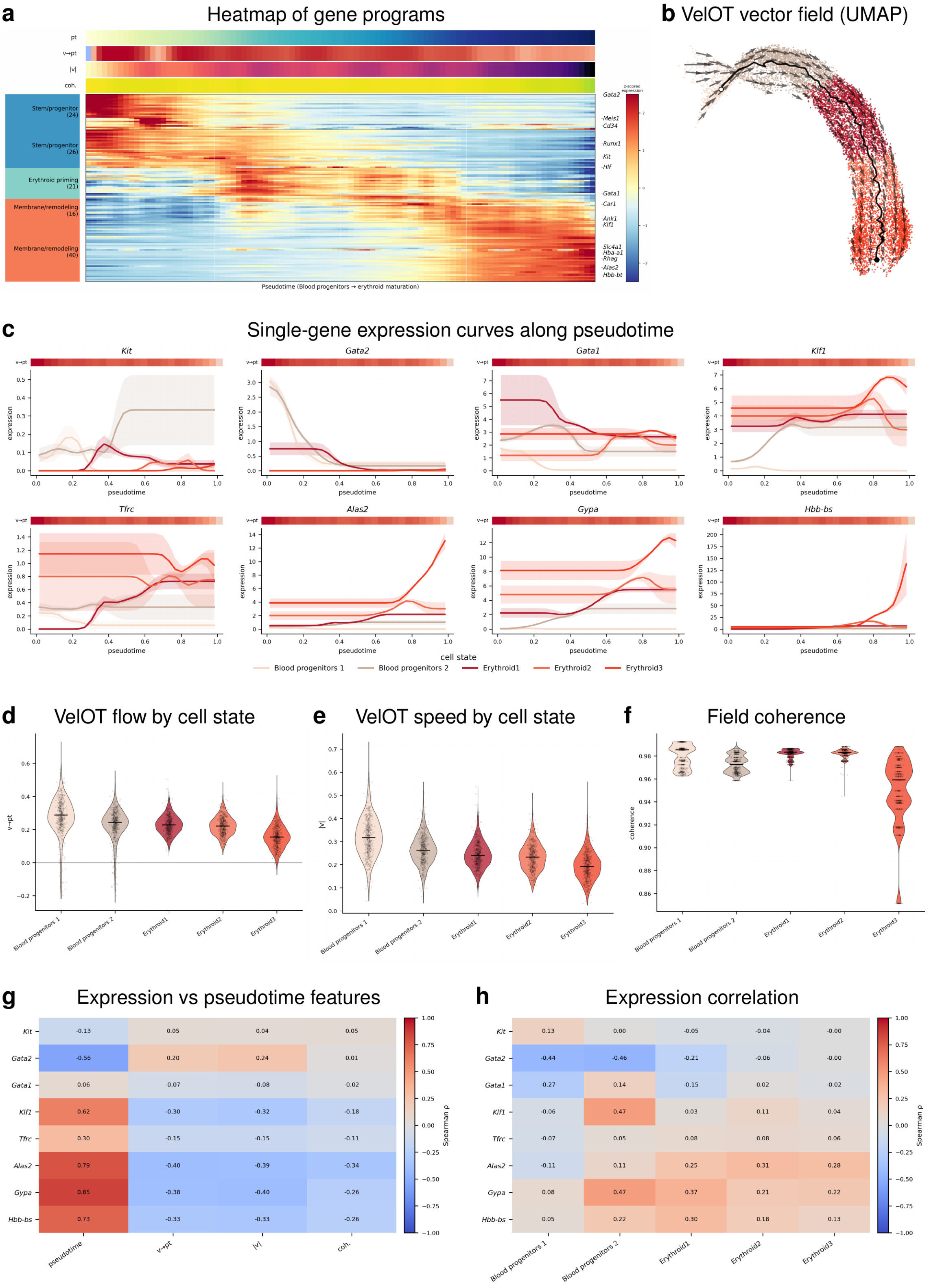

**Fig. S7 Full erythroid VelOT analysis: gene programs, vector-field flow, and feature associations along inferred trajectories. a)** Gene programs (Stem/progenitor 23 genes; Stem/progenitor 17 genes; Erythroid priming 30 genes; Membrane remodeling 19 genes; Iron/heme synthesis 23 genes) along pseudotime; rows are genes, columns are pseudotime bins, color is z-scored expression. **b)** UMAP VelOT vector field with inferred Blood progenitors to Erythroid maturation trajectory. **c)** Single-gene expression curves along pseudotime for the canonical markers *Kit, Gata2, Gata1, Klf1, Tfrc, Alas2, Gypa, Hbb-bs*; color encodes the five clusters. **d-f)** Violin plots of forward VelOT flow $\mathbf{v} \cdot \nabla\tau$, VelOT speed $|\mathbf{v}|$, and local vector-field coherence respectively for each cluster. **g)** Spearman correlations of selected genes with pseudotime, $\mathbf{v} \cdot \nabla\tau$, $|\mathbf{v}|$, and coherence. **h)** Spearman correlations of selected genes within each cell state.

#### S4.4 Per-cell distributions and statistical significance of the benchmark

Figure S8 shows per-cell distributions of CBDir and ICCoh and execution times for all five methods across the four real datasets, with pairwise Wilcoxon tests. The pattern is consistent: VelOT achieves the tightest distributions at the highest values for both CBDir and ICCoh on every dataset, and the differences are highly significant ($P < 10^{-4}$, denoted ****) for all method comparisons except the few pancreas pairs where DeepVelo also achieves near-perfect CBDir.

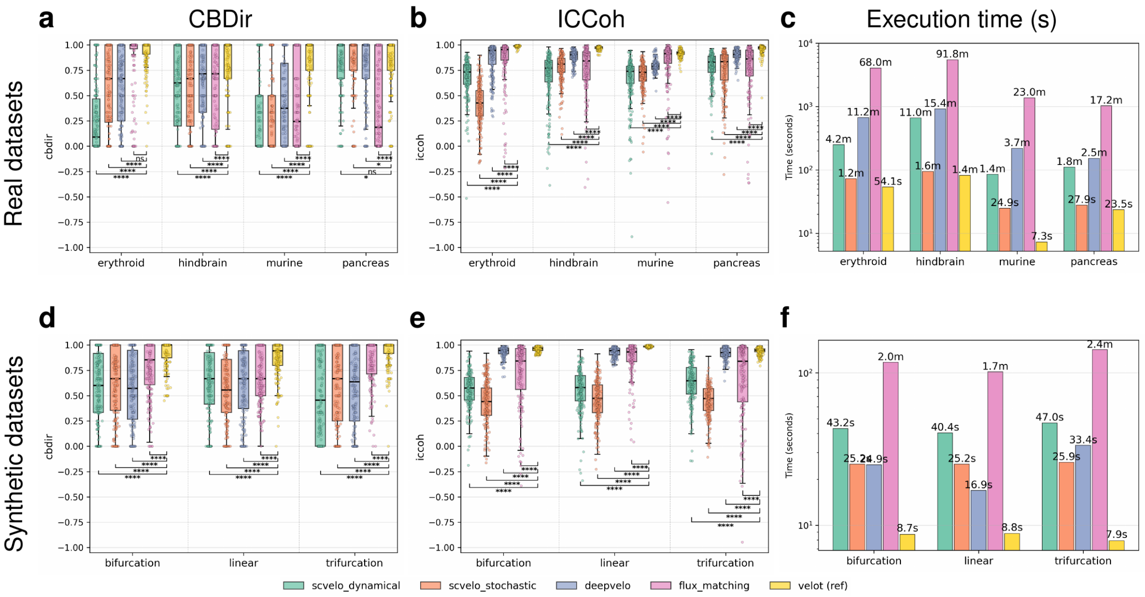

**Fig. S8 Per-dataset distributions and pairwise statistical significance of the RNA-velocity benchmark. a-c)** Box plots of per-dataset CBDir, ICCoh, and total execution time in seconds across the four real datasets (erythroid, hindbrain, murine intestinal organoid and pancreas) and the five benchmarked methods (scVelo dynamical, scVelo stochastic, DeepVelo, FluxMatching and VelOT shown as ref.). Statistical brackets denote two-sided Wilcoxon signed-rank tests with Benjamini-Hochberg correction; each * detones one level of significance of the *p-value*. ns: not significant (*p-value* $> 0.05$). **d-f)** Same results for the three synthetic datasets (linear, bifurcation, and trifurcation). VelOT achieves the most concentrated and highest distributions on every dataset.

##### S4.5 Oligodendroglioma streamlines

Figure S9 shows the VelOT-inferred streamlines on the IDH-mutant 1p/19q co-deleted oligodendroglioma scRNA-seq dataset [69], computed entirely from total counts (no spliced or unspliced inputs). The streamlines correctly recover the two canonical differentiation axes: *Stem-like* cells emit arrows towards both the AC-like (astrocyte-like) and OC-like (oligodendrocyte-like) regions, while a small *Unassigned* fraction sits at the apex of the bifurcation. This pattern matches the developmental hierarchy described in the original publication [69] and the broader neuro-oncology literature [70].

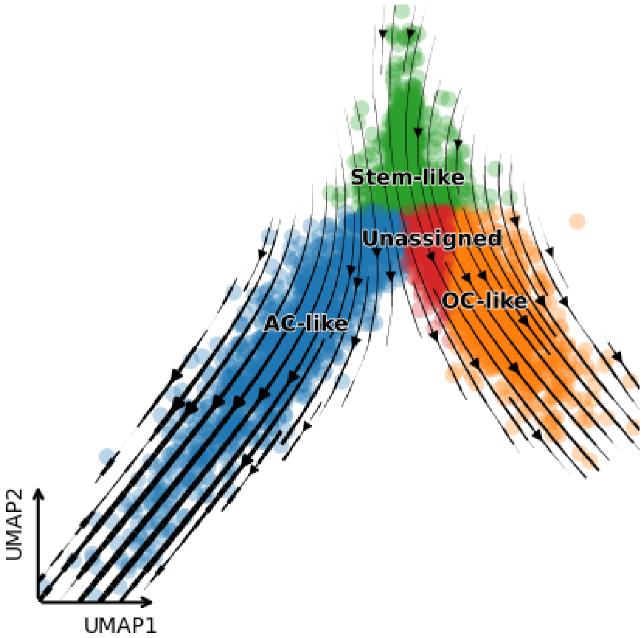

**Fig. S9 VelOT streamlines on adult IDH-mutant 1p/19q co-deleted oligodendroglioma scRNA-seq.** UMAP embedding of tumor cells classified into stem-like (green), AC-like (blue), OC-like (orange), and unassigned (red) following the original cell-type labels of Tirosh *et al.* [69]. VelOT streamlines were computed from total mRNA counts only, without any spliced or unspliced inputs. The vector field correctly recovers the two canonical differentiation axes (stem-like to AC-like and stem-like to OC-like) with the unassigned fraction at the bifurcation apex.

### S5 Synthetic bifurcation simulations, scaling analyses, and ablation experiments

#### S5.1 Synthetic bifurcation design and evaluation metrics

The implementation controls used a synthetic bifurcation dataset with one progenitor compartment (Root) and two terminal branches (Branch 1 and Branch 2). The geometry was generated as a curved two-dimensional manifold with controlled bifurcation structure, then embedded in a higher-dimensional feature space with a known forward direction from Root toward both terminal fates. This design gives access to a ground-truth qualitative topology while remaining close to the geometry encountered in single-cell differentiation maps: cells are unevenly sampled, branches can be imbalanced, and local neighborhoods around the bifurcation contain ambiguous fate information.

The analyses report complementary metrics rather than a single scalar score. CBDir measures whether the inferred cell-wise velocity points toward the correct local branch direction. ICCoh measures local coherence of inferred vectors within neighborhoods. Future alignment measures agreement between the inferred forward displacement and the expected terminal direction after short forward integration. Terminal reach records the fraction of forward trajectories ending in Root, Branch 1, or Branch 2. Fate entropy measures the diversity of terminal routing, with approximately one bit expected for a balanced two-branch fate split and lower values when the field routes cells predominantly to one terminal. Per-cell confidence summarizes the fraction and strength of local OT assignments supporting each velocity estimate; the fraction of zero-confidence cells is used as a failure-mode diagnostic when sampling is sparse or the graph becomes disconnected.

#### S5.2 Hardware timing and hardware-invariant velocity fields

The hardware comparison evaluates whether acceleration changes only runtime or also affects inferred dynamics. Three regimes were compared: GPU execution, single-thread CPU execution (CPU-1), and a multi-worker CPU regime (CPU-8) in which independent window-pair computations are distributed while low-level BLAS/OpenMP oversubscription is avoided. The total runtime decomposition shows that the smooth neural field stage is the dominant accelerated component, whereas window construction, UMAP projection, and metric computation contribute little to total runtime. The GPU achieved the fastest wall-clock time on the tested bifurcation problem and the smallest overall runtime variability. Importantly, the qualitative and quantitative vector fields remained visually and statistically aligned across hardware regimes, showing that hardware selection changes execution speed but not the biological interpretation of the inferred flow.

#### S5.3 Cell-number scaling and stage-wise complexity

Scaling was assessed by increasing the number of cells while retaining the same bifurcation structure. The analysis separates total wall-clock time, stage-level runtime, amortized cost per cell, number of OT window pairs, GPU memory footprint, and quality stability. Over the tested range, total runtime increases much more slowly than the number of cells because fixed overheads and parallel stage execution dominate small instances, whereas the OT and metric stages become increasingly important at larger sample sizes. The OT stage shows the steepest empirical scaling and is therefore the expected bottleneck for substantially larger datasets. GPU memory grows only mildly in this experiment because the implementation processes local windows rather than materializing a full all-to-all transport matrix.

#### S5.4 Ablation design and hyperparameter families

The ablation analysis was organized around four interpretable families. The windowing family modifies how cells are paired across pseudotime windows, including overlap, window size, and unsupervised spatial pre-grouping. The OT-cost family modifies the penalties used to discourage biologically implausible couplings, including time and kNN penalties. The smoothing-loss family tests whether the neural smoother needs explicit smoothness, curl, or divergence constraints. The smoothing-network family changes the architecture or conditioning of the neural field, including hidden width, number of training epochs, pseudotime conditioning, and kNN smoothing scale.

Across these families, the strongest negative control is the raw-OT field without the neural MLP smoother. This condition runs quickly but produces poor CBDir and ICCoh, demonstrating that local OT displacements alone do not form a coherent global velocity field. In contrast, most changes to cost penalties, windowing, and architecture remain close to the baseline in the high-quality regime. These results support the interpretation that VelOT is robust to moderate hyperparameter variation but depends critically on smoothing local transport signals into a coherent vector field.

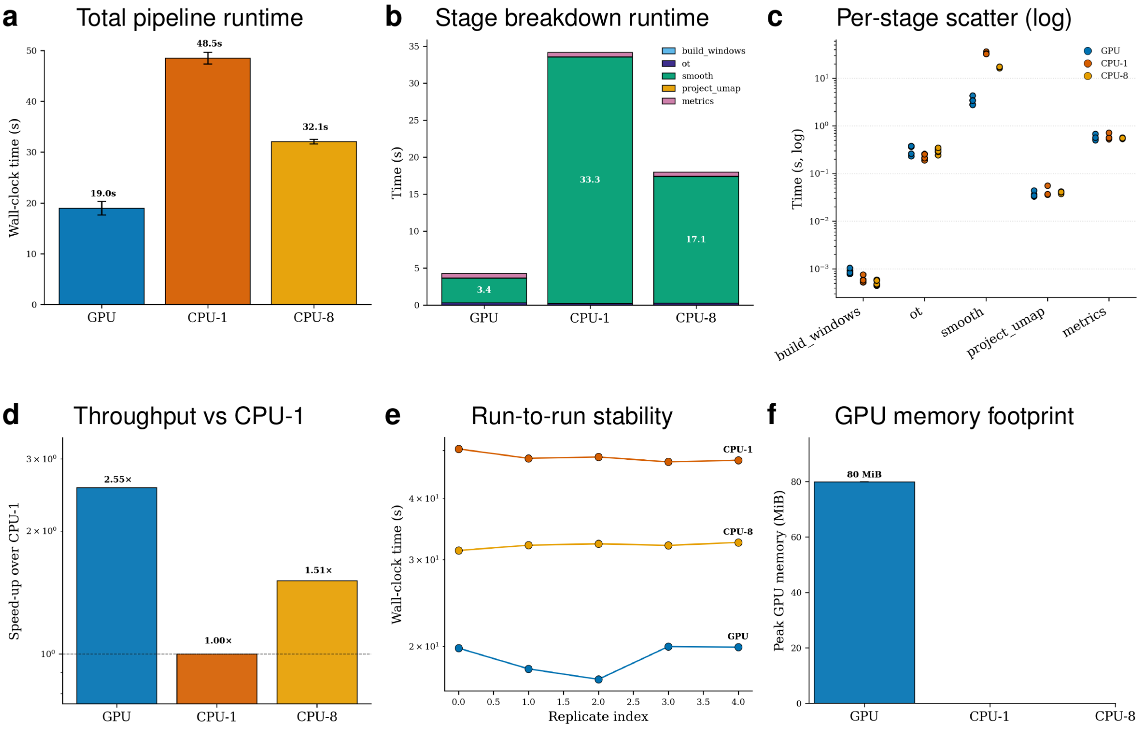

**Fig. S10 Hardware timing on the synthetic bifurcation dataset.** All results are shown for GPU, CPU-1, and CPU-8. GPU execution is fastest overall, CPU-8 improves over CPU-1, and the peak GPU memory footprint remains small for this benchmark.

#### S5.5 Perturbation robustness, terminal routing, and confidence diagnostics

Perturbation sweeps were designed to test three distinct failure modes. Geometry-noise sweeps distort the cell manifold while preserving the intended bifurcation topology. Branch-imbalance sweeps change the relative density of the terminal branches, mimicking rare-fate and over-represented-fate settings. Pseudotime-corruption sweeps perturb temporal ordering, which directly tests whether VelOT remains stable when the root-to-terminal ordering is noisy. The results show that global quality metrics degrade smoothly under geometry noise, but terminal routing and fate entropy are more sensitive to branch imbalance and pseudotime corruption. Confidence diagnostics are therefore important complementary readouts: high global CBDir or ICCoh does not necessarily guarantee balanced recovery of rare fates.

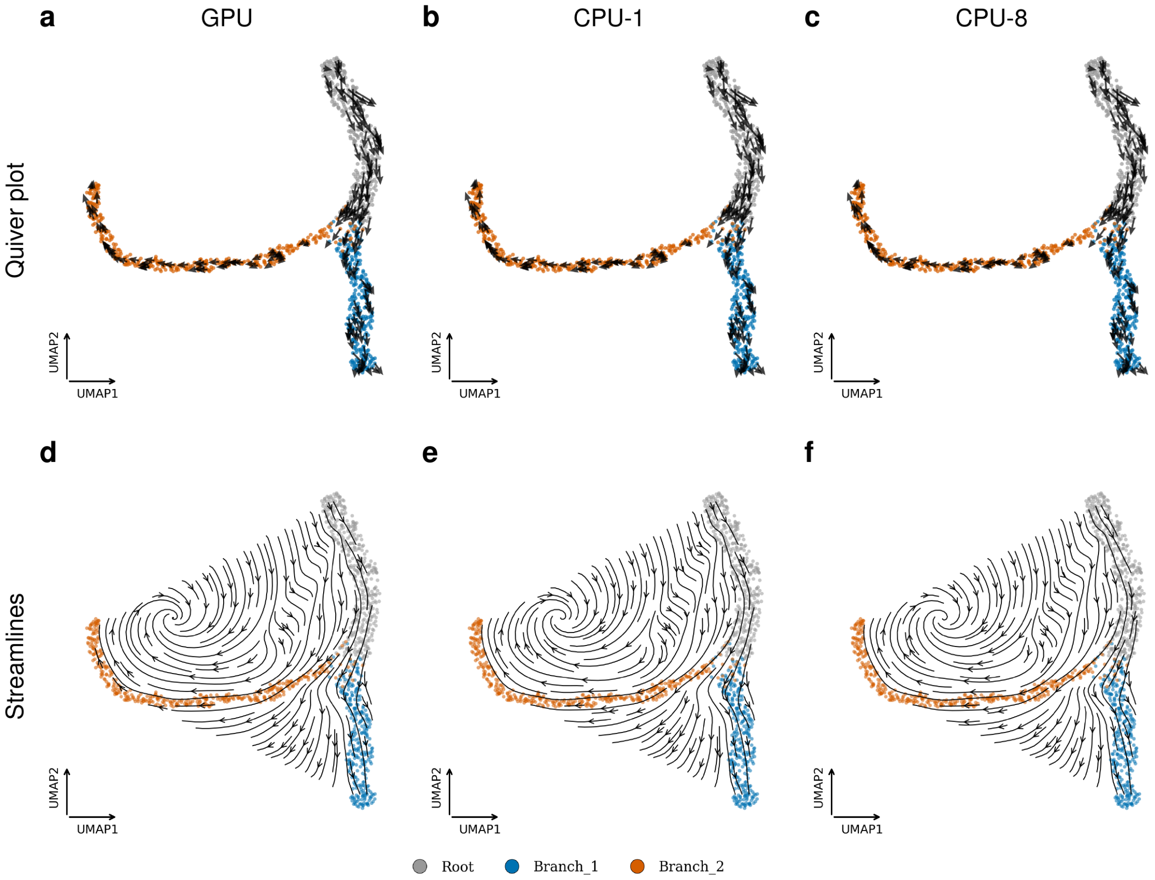

**Fig. S11 Velocity-field equivalence across hardware regimes. a-c)** Quiver plots of the VelOT vectors projected to UMAP coordinates for the GPU, CPU-1, and CPU-8 computations respectively. **d-f)** Streamline visualizations of the VelOT field in UMAP coordinates for the GPU, CPU-1, and CPU-8 computations respectively. The three fields are qualitatively indistinguishable, with all regimes preserving the same bifurcation topology and branch routing.

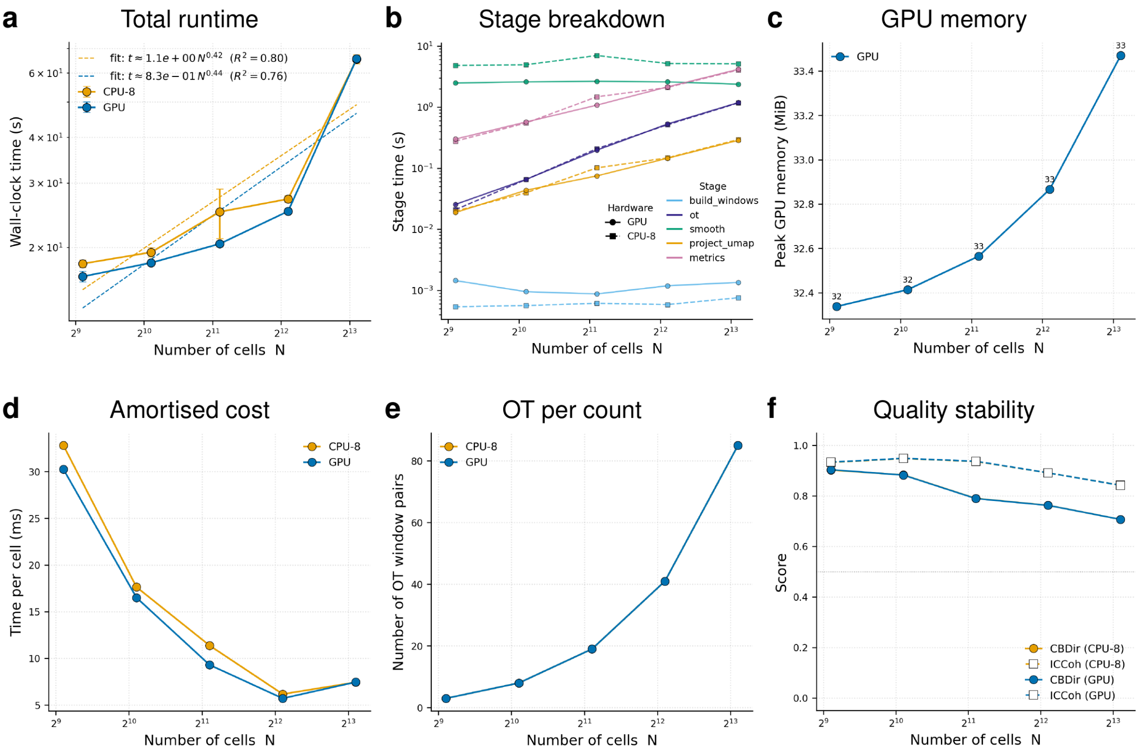

**Fig. S12 Cell-number scaling of runtime, memory, and quality. a-f)** Total runtime, stage breakdown, GPU memory footprint, amortized time per cell, OT window-pair count, and quality stability are shown as a function of cell number (dataset size). GPU and CPU-8 retain similar quality at matched cell number, while runtime differences are driven by stage-specific implementation costs.

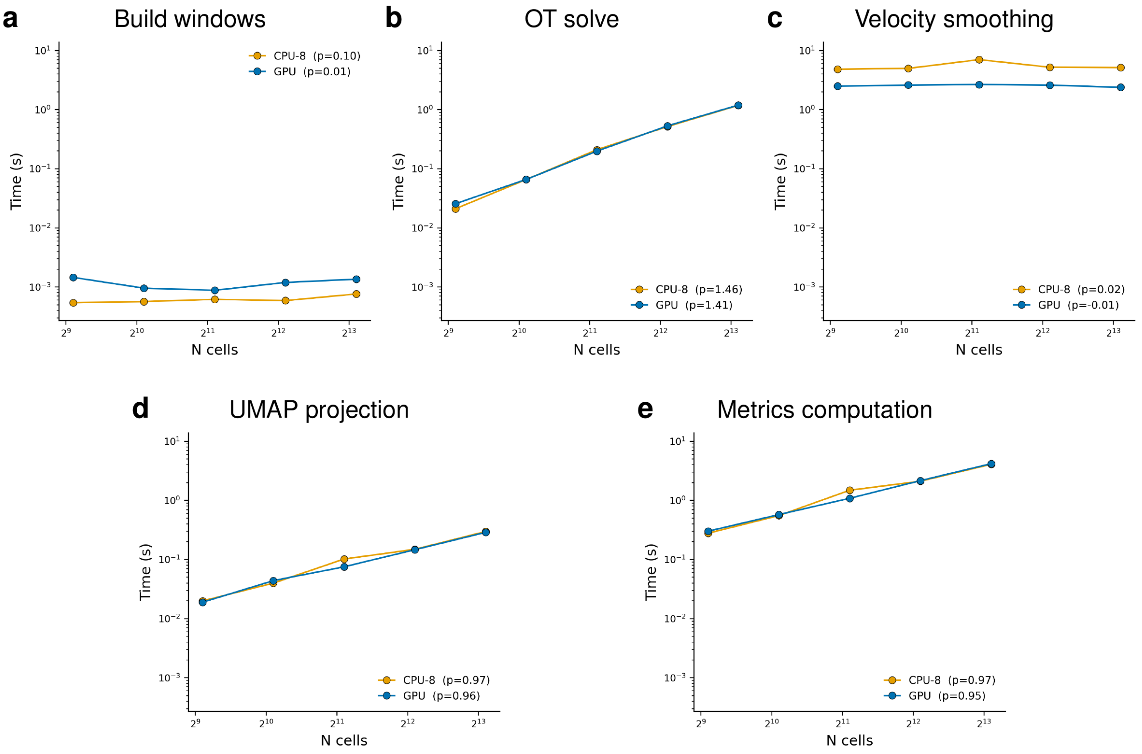

**Fig. S13 Stage-wise scaling on log-log axes. a-e)** Time specific computation cost in seconds as a function of the number of cells (N) of the dataset for the different tasks of the VelOT pipeline; window construction, OT, neural smoothing, UMAP projection, and metric computation. The OT and metric stages show the strongest dependence on cell number, whereas neural smoothing remains comparatively flat across the tested range.

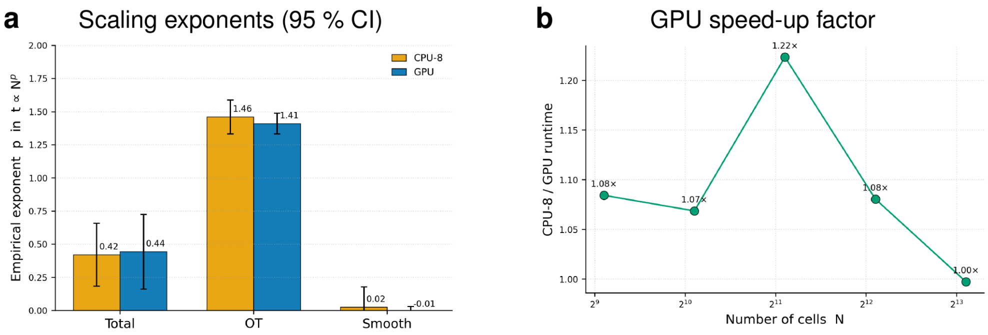

**Fig. S14 Empirical scaling exponents and GPU speed-up factor. a)** Fitted exponents for total runtime, OT runtime, and smoothing runtime under GPU and CPU-8 execution. **b)** GPU speed-up factor relative to CPU-8 as a function of cell number (N). The GPU advantage is largest for small-to-medium inputs and narrows when CPU-side OT and data orchestration dominate.

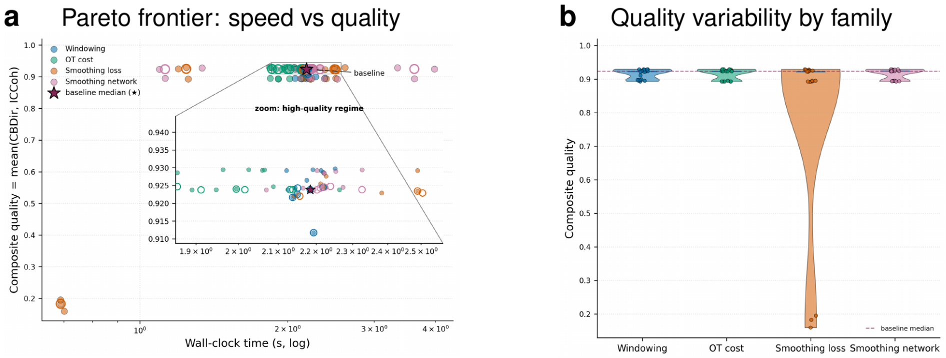

**Fig. S15 Ablation speed-quality Pareto frontier. a)** Composite quality pareto frontier plot against wall-clock time across hyperparameter families. Most ablations cluster near the high-quality baseline regime, while the raw-OT/no-MLP condition is fast but fails to preserve coherent vector-field quality. **b)** Composite quality violin plots across families.

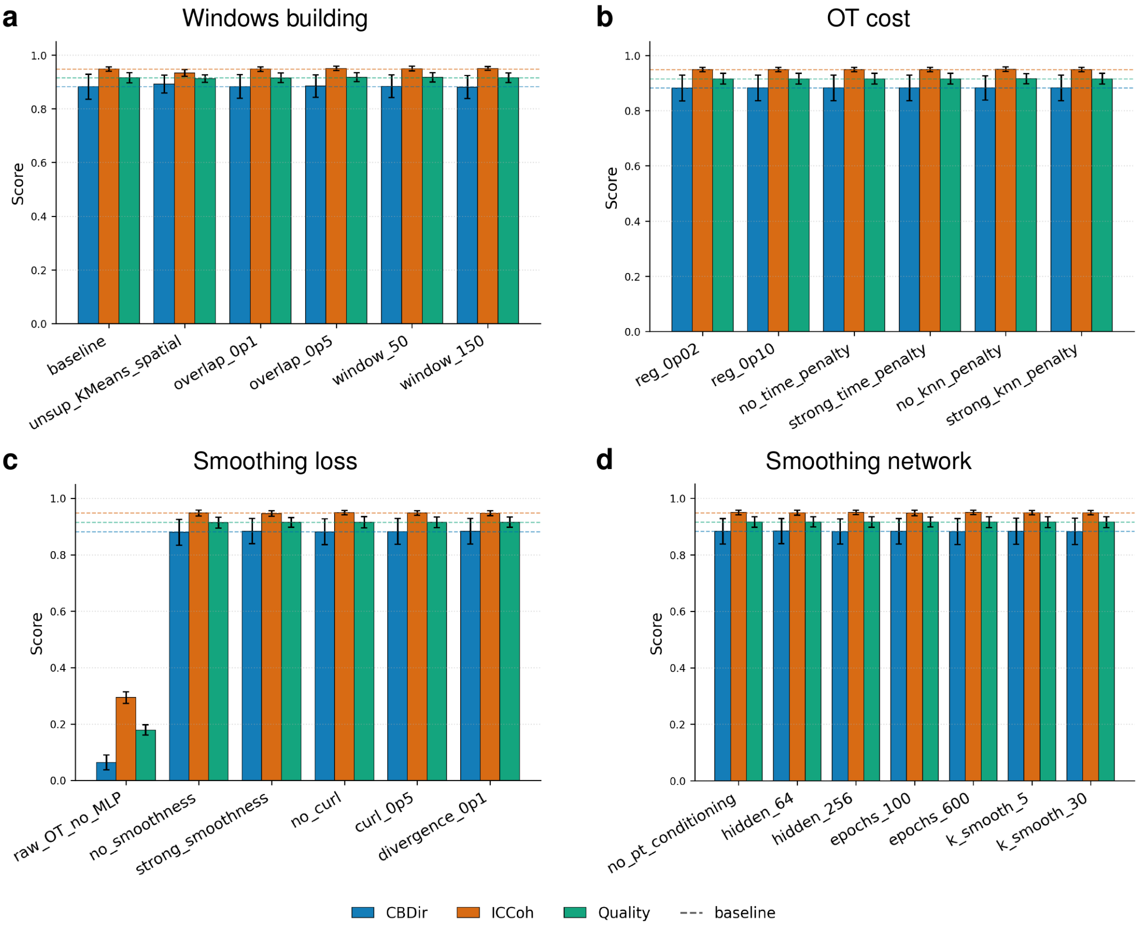

**Fig. S16 Per-family ablation metrics.** Mean CBDir, ICCoh, and composite quality are shown with variability across seeds for **a)** windowing construction, **b)** OT-cost computation, **c)** smoothing-loss, and **d)** smoothing-network ablations. Dashed lines indicate baseline levels. The neural smoother is the most critical component, while most windowing and OT-cost variants remain close to baseline.

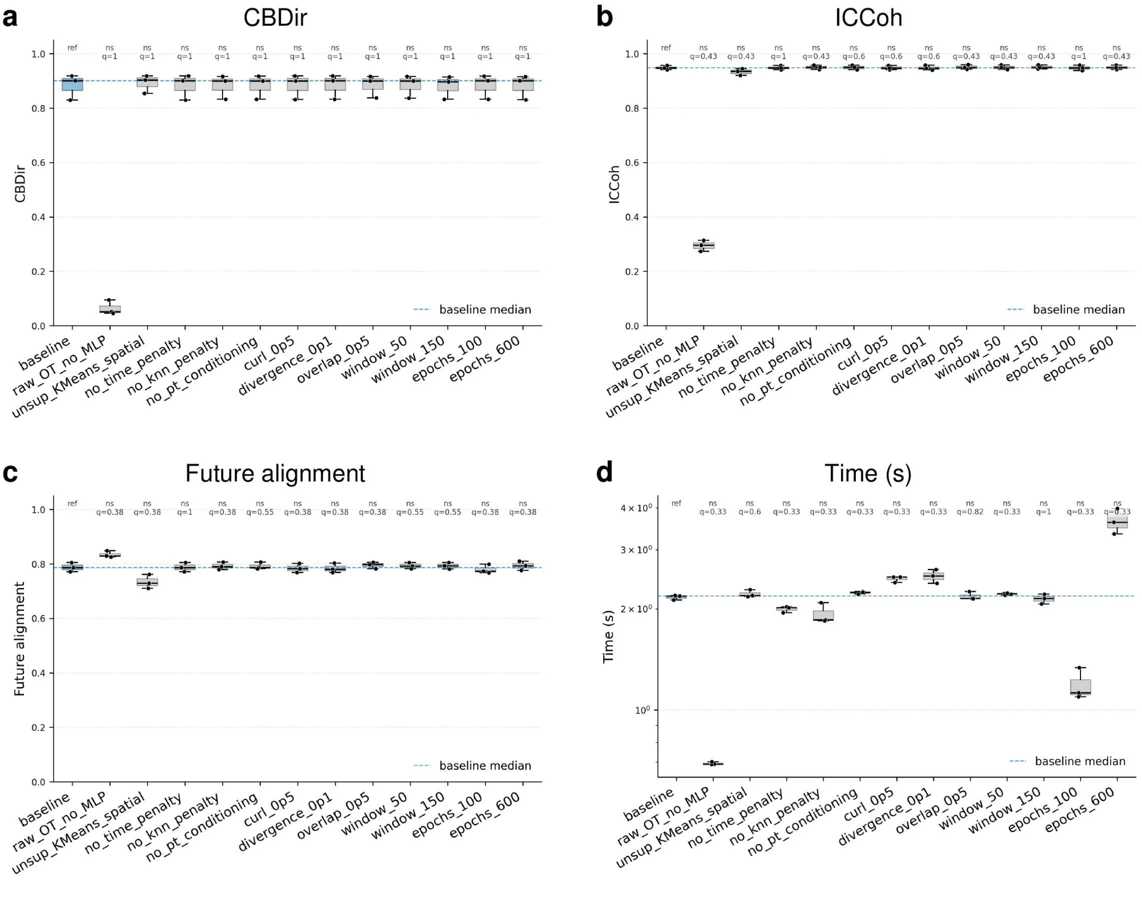

**Fig. S17 Exact paired sign-flip tests for curated ablations. a-d)** CBDir, ICCoh, future alignment, and runtime respectively are compared against the baseline across matched seeds using paired sign-flip tests with BH-corrected $q$ values. The raw-OT/no-MLP control is the only condition that strongly disrupts CBDir and ICCoh, whereas most curated ablations remain statistically close to baseline in quality.

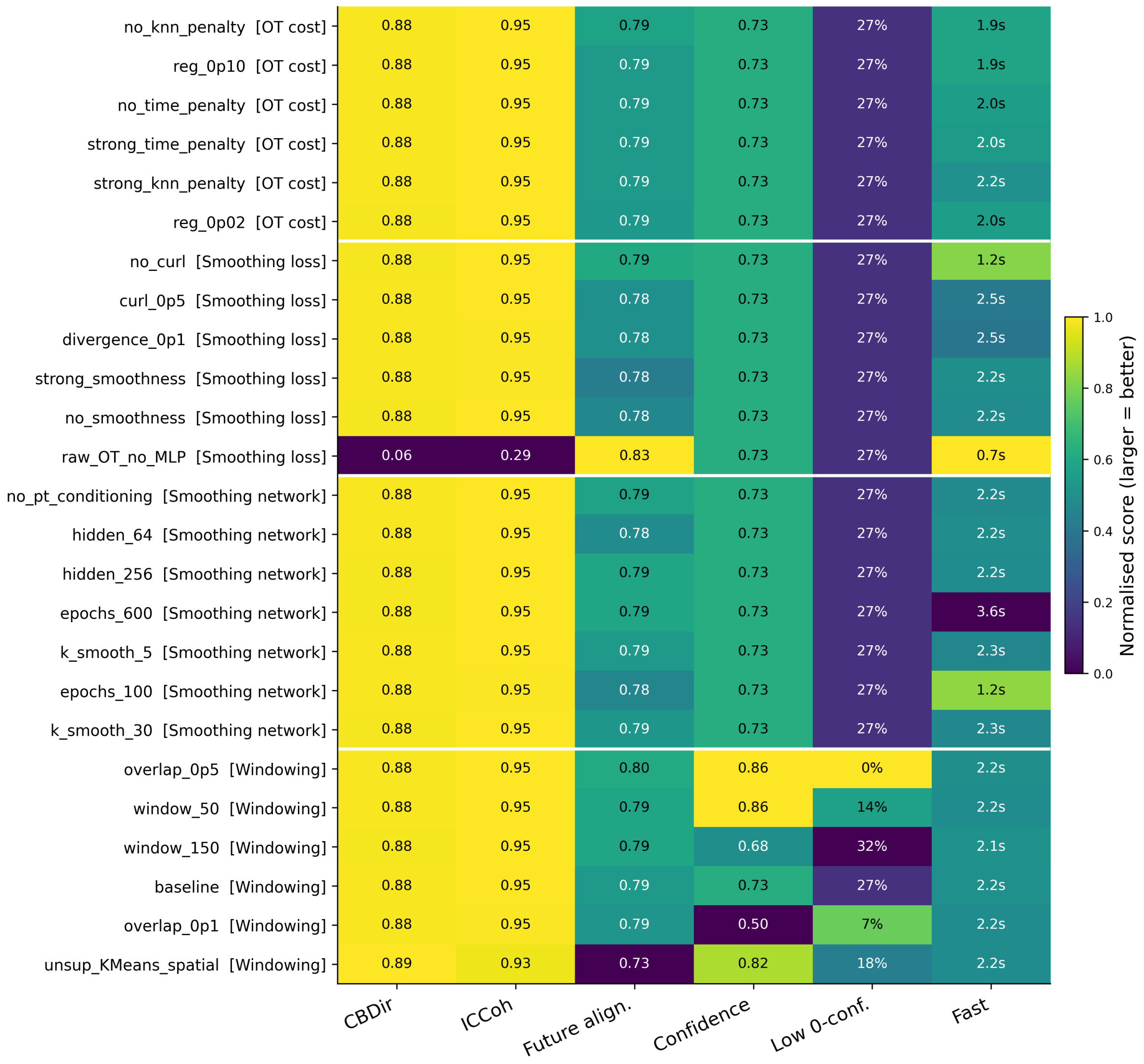

**Fig. S18 Ablation metric heatmap.** Rows correspond to ablation configurations grouped by family, and columns summarize CBDir, ICCoh, future alignment, confidence, low-confidence fraction, and speed. Values are normalized so that higher values are better, with raw values annotated in each cell. The heatmap highlights the failure of raw OT without MLP smoothing and the stability of most regularized configurations.

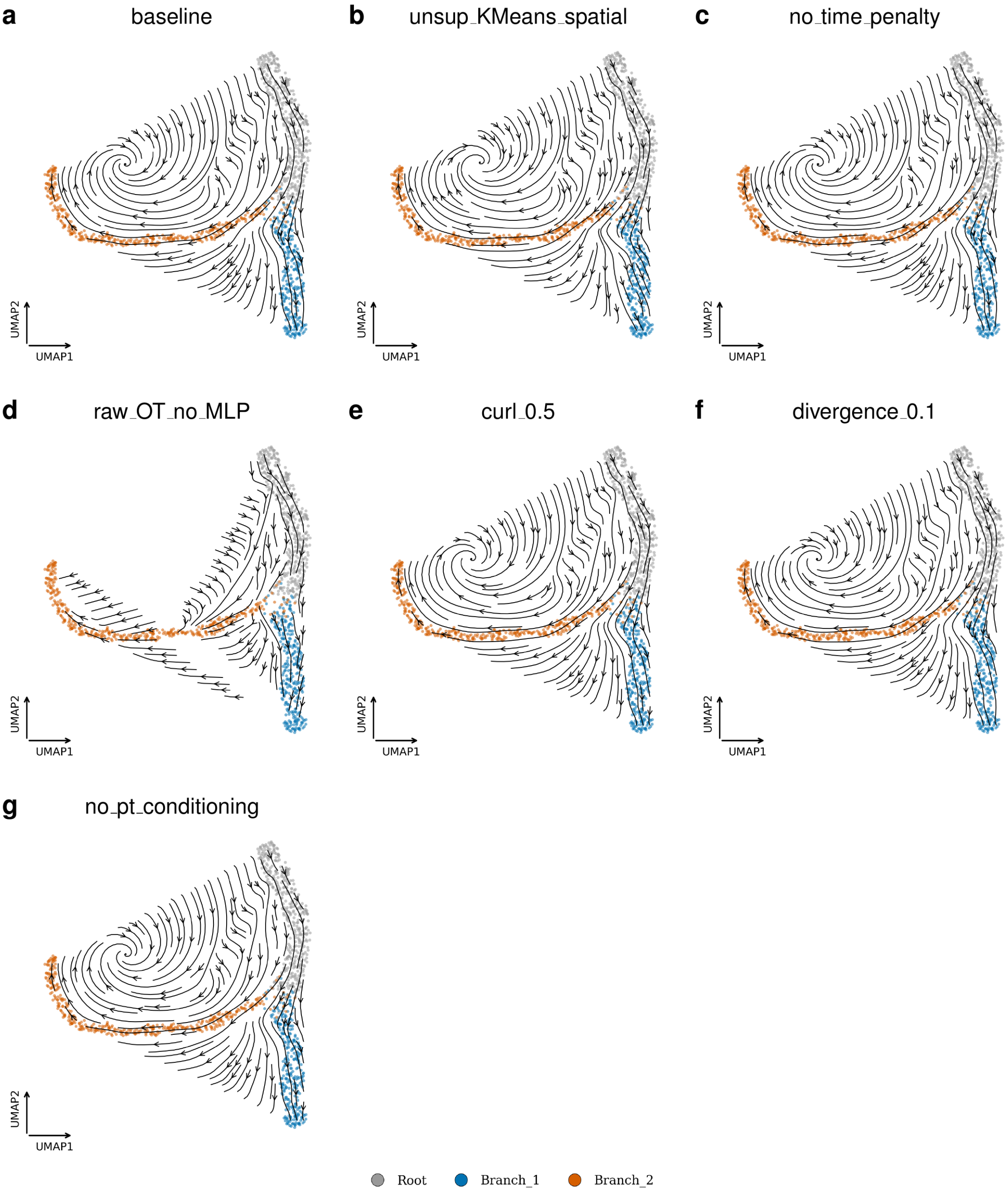

**Fig. S19 Representative velocity fields across ablations. a)** Streamline visualization of the VelOT field for the baseline and **b-g)** for different selected ablations in UMAP coordinates. Most regularized variants preserve the same root-to-branch topology, whereas the raw-OT/no-MLP control produces a fragmented and less coherent field.

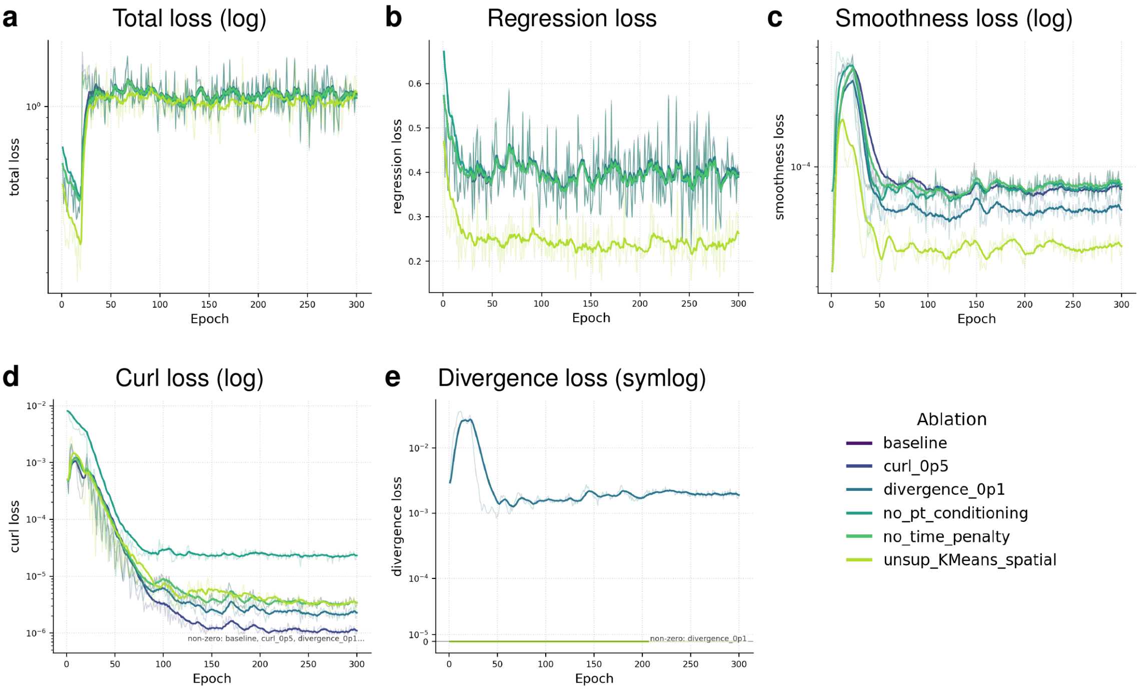

**Fig. S20 Smoothing convergence curves. a-e)** Raw and exponentially averaged training curves are shown for total loss, regression loss, smoothness loss, curl penalty, and divergence penalty. The regression term stabilizes after an early transient, while the curl and divergence penalties decay to low values when activated.

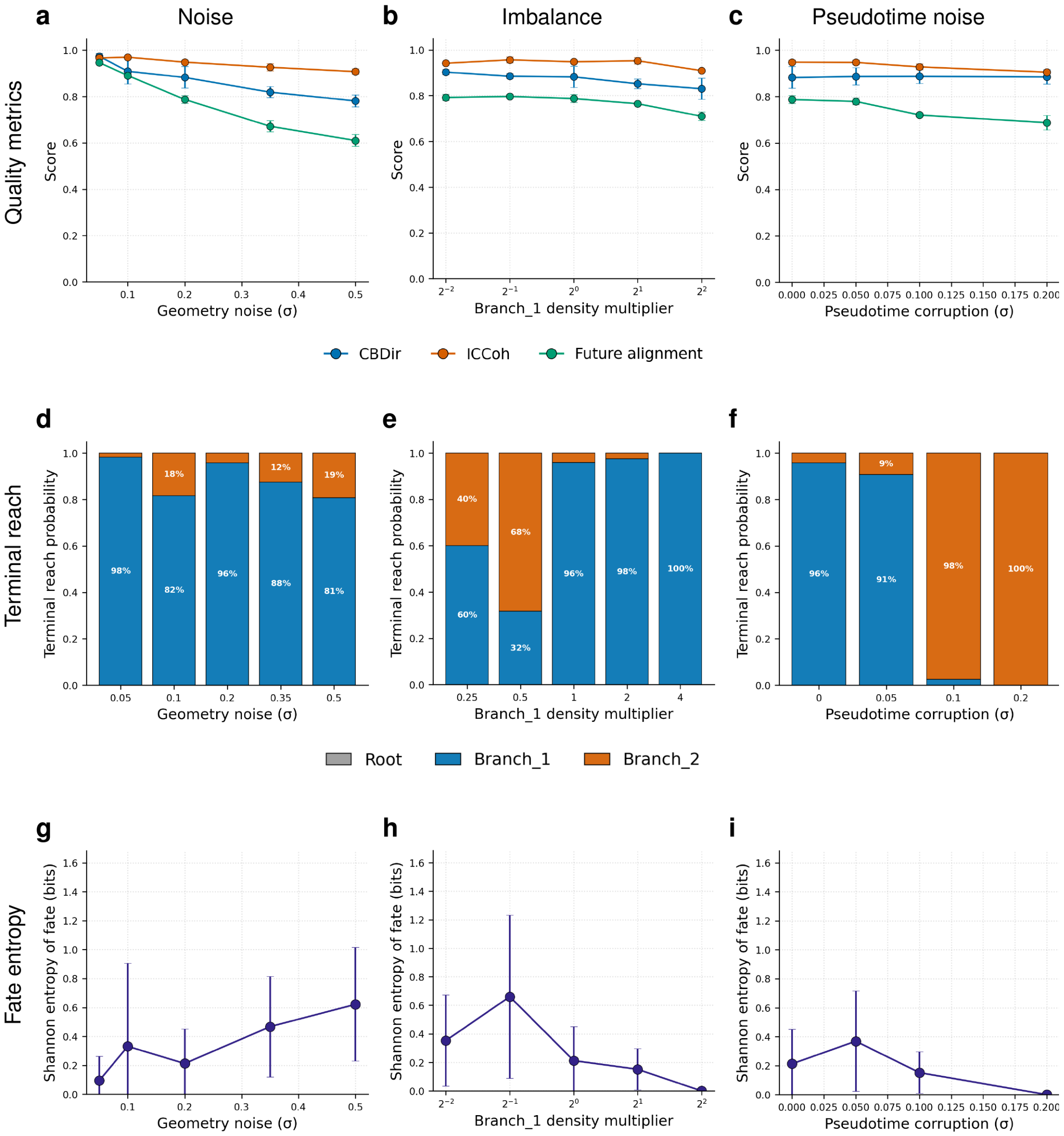

**Fig. S21 Robustness across perturbation sweeps. a-c)** Noise, imbalance and pseudotime noise quality metrics shown for geometry noise, branch-density imbalance, and pseudotime corruption. Geometry noise primarily decreases quality gradually, whereas imbalance and pseudotime corruption mainly alter terminal routing. **d-e)** Terminal reach probability across cell-type clusters for geometry noise, branch-density imbalance, and pseudotime corruption. While terminal reach probability is stasble across different levels of geometry noise, it is more sensitive to branch-density imbalance and pseudotime noise corruption. **d-i)** Fate entropy for geometry noise, branch-density imbalance, and pseudotime corruption.

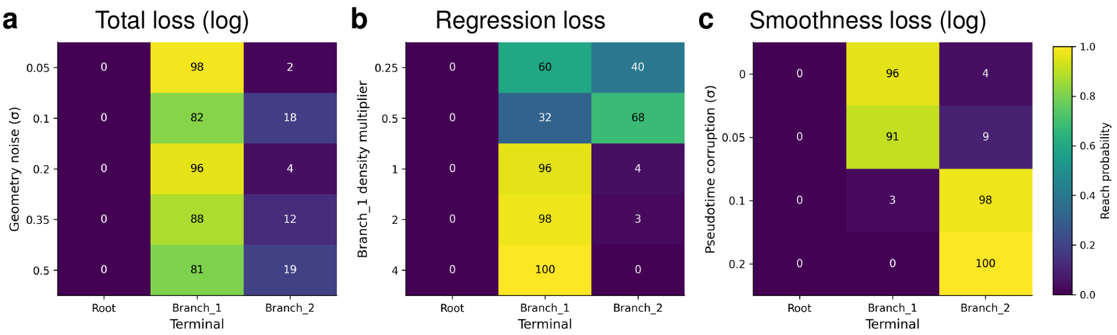

**Fig. S22 Terminal reach matrices across perturbation sweeps. a)** Total loss (log-scale), **b)** regression loss, and **c)** smoothness loss (log-scale) heatmaps. Each heatmap reports the probability that forward-integrated trajectories terminate in Root, Branch 1, or Branch 2 across increasing perturbation levels. The matrices reveal rare-fate under-recovery and pseudotime-induced routing bias that may not be obvious from global quality metrics alone.

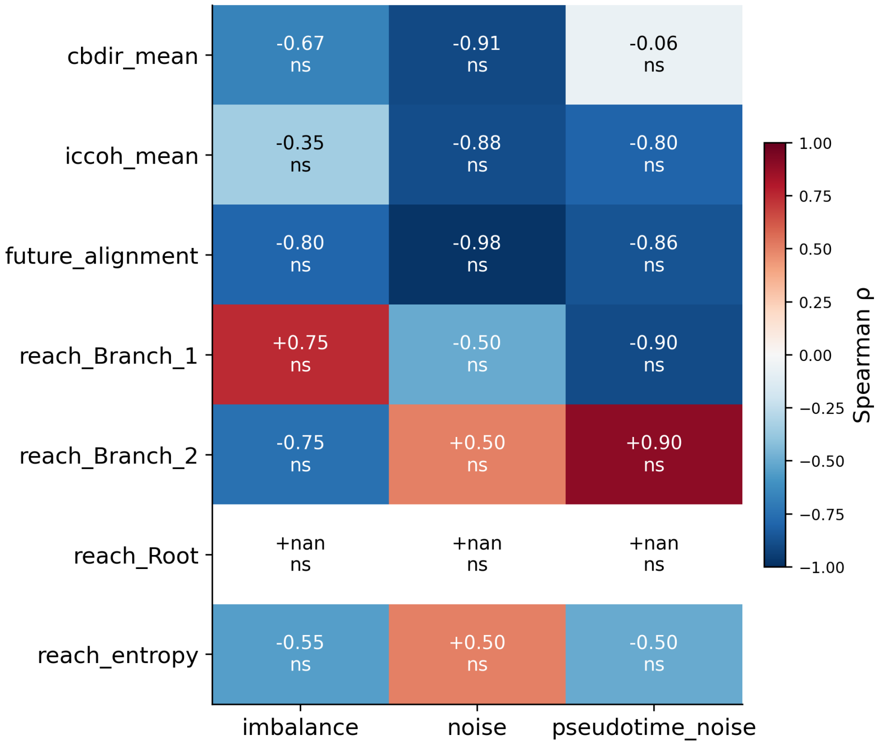

**Fig. S23 Monotonic-trend tests for robustness metrics.** Spearman correlations with BH correction summarize monotonic trends across imbalance, geometry noise, and pseudotime noise. The analysis separates quality metrics from terminal reach and entropy, emphasizing that topology-level routing diagnostics can change even when global quality remains relatively stable.

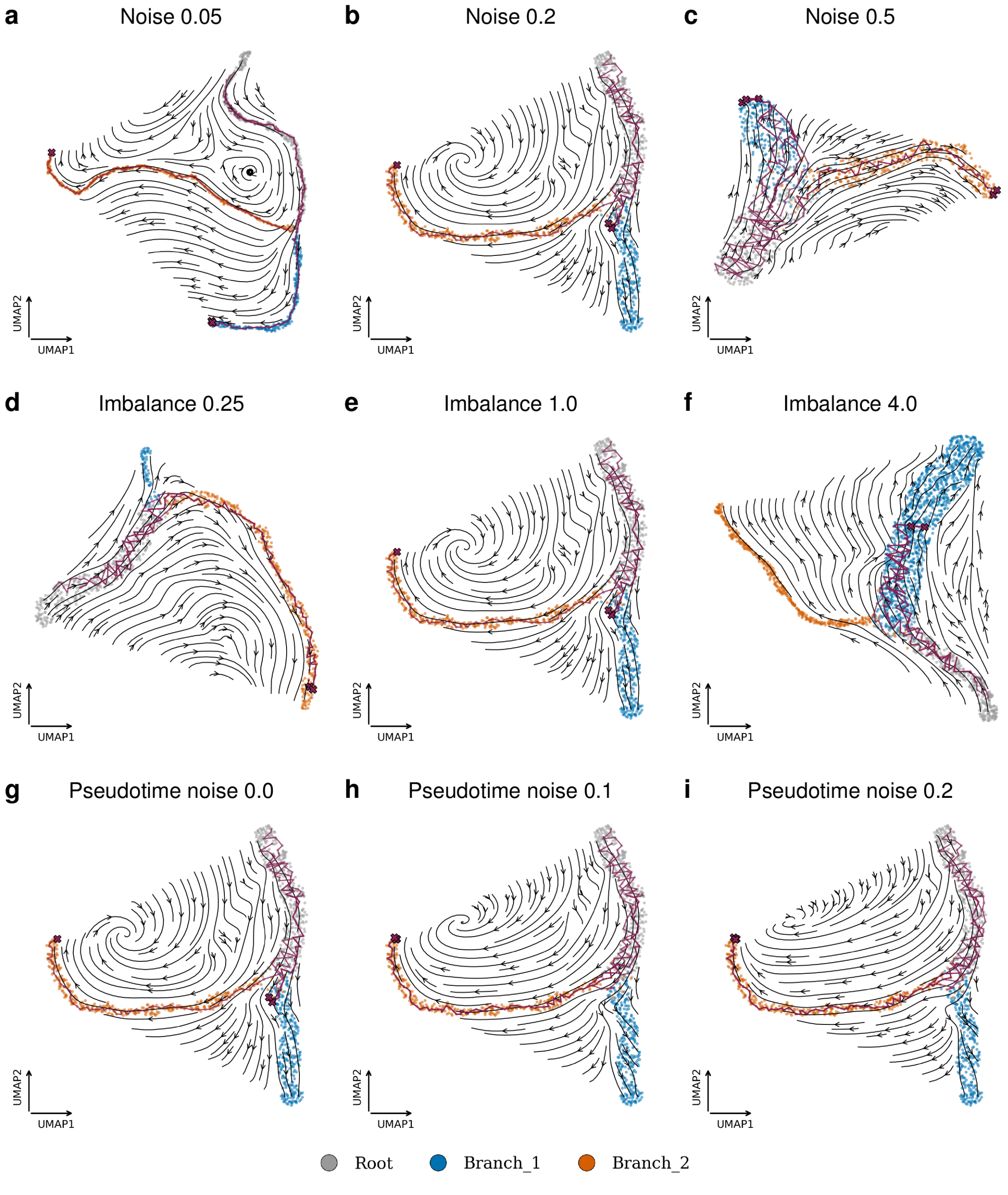

**Fig. S24 Representative VelOT fields with integrated forward trajectories from Root. a-i)** Streamlines from the VelOT field and integrated trajectories are shown in UMAP coordinates for representative geometry-noise, imbalance, and pseudotime-corruption settings. Moderate perturbations preserve the expected two-branch topology, whereas stronger pseudotime corruption biases route selection.

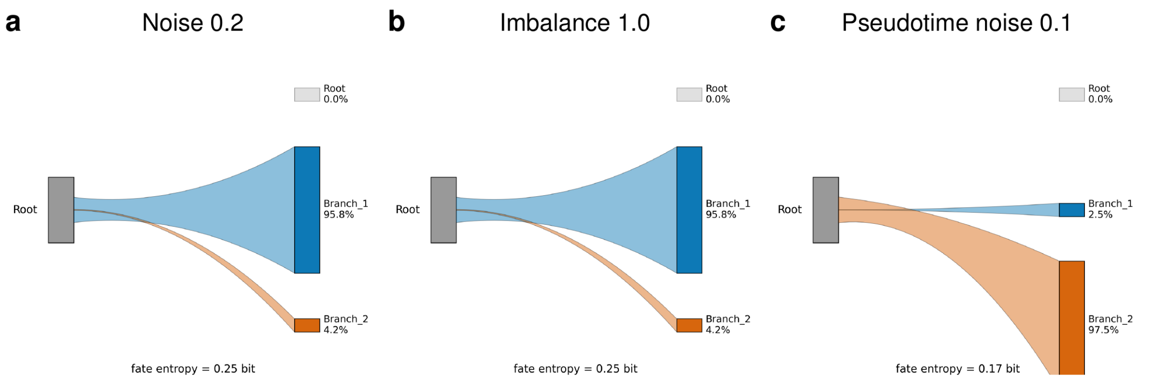

**Fig. S25 Alluvial routing diagrams from Root to terminal states.** Representative routing diagrams for **a)** 0.2 noise level, **b)** 0.1 fraction of imbalance and **c)** pseudotime corruption level of 0.1. The diagrams show how the fraction of trajectories reaching Branch 1 or Branch 2 changes under selected perturbation settings. The fate entropy annotation summarizes how balanced or collapsed the terminal routing is.

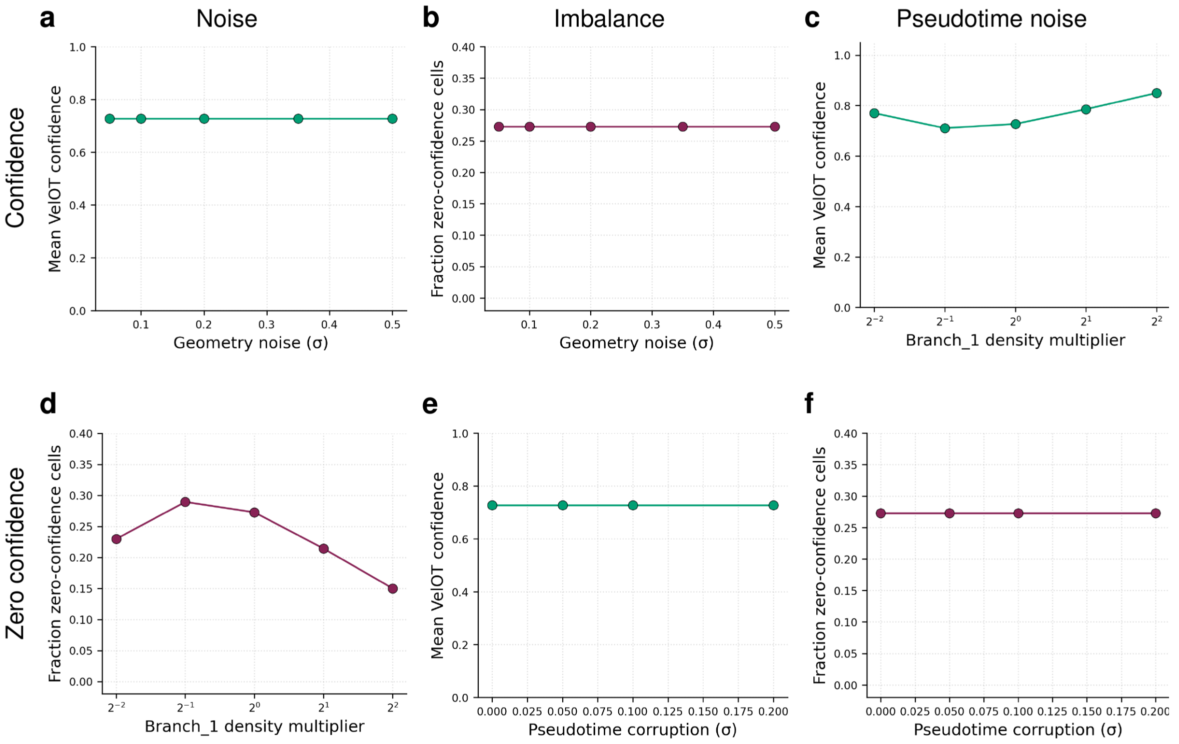

**Fig. S26 Per-cell confidence diagnostics across perturbation sweeps. a-c)** Mean VelOT confidence under geometry noise, branch imbalance, and pseudotime corruption respectively. **d-f)** Fraction of zero-confidence cells for the same case studies. Confidence is stable for geometry and pseudotime noise in this experiment, but branch imbalance changes the fraction of cells with zero confidence, supporting its use as a practical rare-fate diagnostic.

### S6 Implementation, outputs, and recommendations

#### S6.1 Stored outputs in the AnnData object

The VelOT pipeline writes its outputs to standard slots of the AnnData object. Table S2 lists the most important keys.

**Table S2**: Key outputs stored in `adata` after running VelOT and VelOT-MetaFlow.

| Location | Key | Meaning |
|---|---|---|
| *Preprocessing and core VelOT outputs* | | |
| obs | `velot_pseudotime` | Normalized pseudotime $\tau_i \in [0, 1]$ |
| obs | `velot_confidence` | Per-cell OT confidence $c_i \in [0, 1]$ |
| obsm | `velot_velocity_latent` | Smoothed velocity in PCA space $\mathbf{v}_i \in \mathbb{R}^D$ |
| obsm | `velot_velocity_umap` | Locally projected velocity in 2D embedding |
| *VelOT-MetaFlow preprocessing and diagnostics* | | |
| obs | `velot_metaflow_time_bin` | Discrete pseudotime bin $b_i \in \{0, \ldots, B - 1\}$ |
| obs | `velot_metaflow_divergence` | z-scored divergence $\hat{\nabla} \cdot \mathbf{V}_i^*$ |
| obs | `velot_metaflow_emb_curl` | z-scored embedding curl $\lvert\omega_i\rvert$ |
| obs | `velot_metaflow_ot_alignment` | Cosine alignment $a_i$ |
| obsm | `velot_metaflow_future_latent` | Future state $\mathbf{Z}^{\text{future}}$ |
| obsm | `velot_metaflow_velocity_final` | Final blended velocity $\mathbf{V}^*$ |
| *VAMPFlow outputs* | | |
| obs | `velot_vampflow_state` | Hard assignment $\arg\max_k q_{ik}$ |
| obs | `velot_vampflow_committor_M{t}` | Per-cell committor to terminal $t$ |
| obsm | `velot_vampflow_soft_states` | Soft memberships $q_{ik}$ |
| uns | `velot_vampflow_transition_matrix` | $\mathbf{T}^{\text{VAMP}} \in \mathbb{R}^{K \times K}$ |
| uns | `velot_vampflow_state_summary` | State-level DataFrame |
| *PAGA-like connectivity* | | |
| uns | `{key}_paga_like_connectivity` | Dict with all matrices below |
| | `.confidence` | PAGA confidence $\text{conf}_{ab}$ |
| | `.enrichment` | Degree-corrected enrichment $\text{enr}_{ab}$ |
| | `.zscore` | Permutation $z$-score $z_{ab}$ |
| | `.pvalue` | Empirical $p$-value $p_{ab}$ |
| | `.qvalue` | BH-corrected $q$-value |
| | `.directed_flux` | Group flux $F_{ab}$ |
| | `.directionality` | Directionality $D_{ab}$ |

#### S6.2 Hyperparameter table

Table S3 lists the default hyperparameters used throughout the paper. All values are exposed in the `velot` YAML configuration files and can be cross-validated on a held-out subset of the data.

#### S6.3 Practical recommendations

Based on extensive use of the pipeline on the benchmark datasets and on internal cohorts, we recommend the following.

1. *Velocity quality first.* Inspect the alignment map before interpreting meta-states. A median alignment greater than 0.3 across the whole data is a healthy threshold; lower values typically reflect a poor root specification or excessive regularization.
2. *VAMP-2 score as a convergence diagnostic.* Monitor the VAMP-2 score during training. A plateau within the first 600 epochs indicates good convergence; oscillation suggests that the learning rate or batch size should be tuned.
3. *Selection of $K$.* Prefer $K \in [6, 10]$ for typical scRNA-seq datasets. Very large $K$ (greater than 15) tends to fragment biologically coherent states.
4. *PAGA q-values.* Report edges with $q_{ab} < 0.10$ only; edges with $q_{ab} < 0.01$ and $\text{conf}_{ab} > 0.30$ define the robust backbone of the transition graph.

**Table S3** Default hyperparameters of the VelOT pipeline and VelOT-MetaFlow module used in the manuscript analyses. Values should be treated as approximate reproducibility defaults rather than universal optima. It is recommended to analyse each dataset in order to exploit the pipeline performance by using appropriate parameters.

| Symbol | Name | Default | Role |
|---|---|---|---|
| $D$ | PCA components | 10 to 50 | Latent space dimension |
| $k$ | kNN neighbors | 30 | Manifold and smoothing kNN graph |
| $W$ | Window size | 80 | Cells per temporal window |
| $f_{\text{overlap}}$ | Window overlap | 0.5 | Temporal window overlap fraction |
| $\varepsilon$ | Sinkhorn regularization | 0.05 | Entropy strength in OT |
| $I_{\text{Sink}}$ | Sinkhorn iterations | 80 | Default Sinkhorn loop iterations |
| $\lambda_{\text{time}}$ | Time penalty | 1.0 | OT cost backward-pseudotime weight |
| $\lambda_{\text{knn}}$ | kNN penalty | 0.5 | OT cost off-manifold weight |
| $\lambda_{\text{smooth}}$ | Smoothness weight loss | 0.5 | MLP kNN smoothness penalty |
| $\lambda_{\text{curl}}$ | Curl weight loss | 0.5 | MLP curl penalty |
| $\lambda_{\text{divergence}}$ | Divergence weight loss | 0.0 | MLP divergence penalty |
| $T_{\text{MLP}}$ | MLP epochs | 300 | VelOT MLP training epochs |
| $\eta$ | Learning rate | $10^{-3}$ | AdamW learning rate |
| $T_{\text{flow}}$ | Flow epochs | 1,200 | CFM training epochs (MetaFlow) |
| $T_{\text{VAMP}}$ | VAMP epochs | 1,200 | VAMPFlow training epochs |
| $\lambda_{\text{bal}}$ | Balance weight | 0.20 | Meta-state balance regularizer |
| $\lambda_{\text{sharp}}$ | Sharpness weight | 0.02 | Meta-state sharpness regularizer |
| $\lambda_{\text{orth}}$ | Orthogonality weight | 0.02 | Meta-state orthogonality regularizer |
| $B$ | Pseudotime bins | 8 | OT bin marginals |
| $\Delta t$ | Forward step | 0.15 | Forward propagation timestep |
| $n_{\text{perm}}$ | Permutations | 200 | PAGA null-model permutations |
| $T$ | Softmax temperature | 0.25 | Directed-edge sharpness |
| $\delta_{\text{conf}}$ | PAGA edge threshold | 0.15 | Minimum confidence for edge inclusion |

5. *Cross-validation with cell-type labels.* Run the Fisher exact-test contingency analysis (main Fig. 6c,d) systematically. Strong, sparse enrichment (one or two cell types per meta-state with FDR $< 0.05$) indicates a biologically valid partition; broad, weak enrichment suggests the meta-state count $K$ should be reduced.
6. *Embedding choice.* VelOT operates in PCA space, but the visualization uses UMAP. The local Jacobian projection makes velocity directions robust to UMAP non-linearities; nevertheless, for very small datasets ($N < 500$), consider visualizing on PHATE [27] or diffusion-map components.

[19] D. Klein, T. Uscidda, F. J. Theis, and M. Cuturi. Entropic (Gromov) Wasserstein flow matching with applications to single-cell genomics. *Advances in Neural Information Processing Systems*, 37,

2024.

[20] C. Bunne et al. Learning single-cell perturbation responses using neural optimal transport. *Nature Methods*, 20(11):1759–1768, 2023. doi: 10.1038/s41592-023-01969-x.

[21] Y. Lipman, R. T. Q. Chen, H. Ben-Hamu, M. Nickel, and M. Le. Flow matching for generative modeling. In *The Eleventh International Conference on Learning Representations*, 2023.

[22] R. T. Q. Chen, Y. Rubanova, J. Bettencourt, and D. K. Duvenaud. Neural ordinary differential equations. In *Advances in Neural Information Processing Systems*, volume 31, 2018.

[23] A. Tong et al. Improving and generalizing flow-based generative models with minibatch optimal transport. *Transactions on Machine Learning Research*, 2024. URL https://openreview.net/forum?id=CD9Snc73AW.

[24] M. S. Albergo and E. Vanden-Eijnden. Building normalizing flows with stochastic interpolants. In *The Eleventh International Conference on Learning Representations*, 2023.

[46] I. Loshchilov and F. Hutter. Decoupled weight decay regularization. In *International Conference on Learning Representations*, 2019.

[57] D. P. Kingma and J. Ba. Adam: a method for stochastic optimization. In *International Conference on Learning Representations*, 2015.
